# Mechanism of single-strand annealing from native mass spectrometry and cryo-EM structures of RAD52 homolog Mgm101

**DOI:** 10.1101/2025.10.26.684629

**Authors:** Carter T. Wheat, Zihao Qi, Miqdad Hussain, Katerina Zakharova, Vicki H. Wysocki, Charles E. Bell

## Abstract

RAD52, the primary single-strand annealing (SSA) protein in humans, forms undecameric rings that bind ssDNA within a narrow, positively-charged groove. Whether RAD52 anneals two complementary ssDNAs on the same ring in *cis*, or between two ring-ssDNA complexes in *trans,* is unknown. Here, we determined cryo-EM structures of Mgm101, a RAD52 homolog from yeast mitochondria, in complexes with ssDNA, a duplex intermediate of annealing, and B-form dsDNA product. In all states, Mgm101 forms a closed nonadecameric ring that binds the backbone of the first ssDNA at the base of the narrow groove. The second complementary strand binds directly on top of the first to form an extended, unwound, and circular duplex intermediate of annealing. The third complex captures apparent B-form DNA product bound to a novel β-hairpin motif located on top of the Mgm101 ring, above the primary DNA-binding groove. Mass photometry and native mass spectrometry confirm and further elucidate the complexes formed in solution. Altogether, our data reveal the full SSA pathway of Mgm101 and suggest it anneals two complementary ssDNAs on the same ring in *cis*. Structural conservation with RAD52 suggests it is likely to use a similar *cis* mechanism of annealing.

## Introduction

DNA damage by endogenous and exogenous agents must be repaired to maintain genomic stability^1^. Among the most toxic DNA lesions are double-stranded DNA breaks (DSBs), which can be repaired by several pathways depending on the cell cycle^2^. Homologous recombination (HR), deployed during S/G2 phases, uses a sister chromatid as a template for highly accurate repair^1^. In M/G1 phases where no such template is available, cells predominantly use non-homologous end-joining (NHEJ) to process DNA termini and directly ligate ends, often with short alterations^1,3^. In cases where the DSB is flanked by directly repeated sequences, the cell can use a third pathway called single-strand annealing (SSA)^3^. In both HR and SSA the DNA ends are resected to generate long 3’ overhangs coated with replication protein A (RPA)- a crucial intermediate. In HR, BRCA2 facilitates exchange of RPA for RAD51 to promote strand invasion into a homologous donor template^4–8^. In SSA, RPA is alternatively replaced by RAD52, which can directly anneal the complementary regions in an ATP-independent manner^9,10^. After cleavage of the non-complementary overhangs and strand ligation, the DSB is repaired, but with loss of genetic information. Inactivation of RAD52 is synthetically lethal with deficiencies in BRCA1 or BRCA2, and has thus been targeted for treatment of breast and ovarian cancers^11–14^.

RAD52 is a 418 amino acid protein with an N-terminal domain for DNA binding and oligomerization (DBD) and a largely disordered C-terminal domain (CTD) for mediating interactions with multiple proteins including RPA and RAD51^15–19^. Crystal structures of the DBD reveal an undecameric ring with two separate binding sites for DNA^20,21^. The inner site is located within a deep and narrow positively-charged groove that binds the backbone of ssDNA in an extended conformation with its bases exposed for homology recognition^21^. The outer site, located at the periphery of the groove, binds dsDNA or ssDNA in a helical conformation that bridges two neighboring RAD52 rings in a crystal structure^21^. Exactly how these two sites function together in annealing is currently unknown. Different models have been proposed to explain the RAD52 annealing mechanism, including a *cis* process in which both ssDNA strands are paired on the same ring^9,22^, or a *trans* process in which annealing occurs between two ssDNA-loaded rings^21,23,24^. To date, a structure of RAD52 bound to two complementary DNA strands simultaneously has yet to be determined, leaving these models up for debate. A more complete understanding of RAD52’s interactions with DNA during SSA is needed to better inform design of effective inhibitors for cancer treatment.

RAD52 is a member of the single-strand annealing protein (SSAP) family, which includes a diverse group of proteins from bacteriophage that have been exploited in powerful methods for bacterial genome engineering including recombineering and MAGE (multiplex automated genome engineering)^25,26^. Despite limited sequence homology, RAD52 and phage SSAPs share a conserved core fold comprised of a β-β-β-α motif and β−hairpin motif that form the inner DNA binding site. Like RAD52, phage SSAPs also contain a C-terminal domain for interacting with partner proteins^18,27,28^, including a phage-encoded exonuclease and the host single-stranded DNA binding protein (SSB)^27,28^. In contrast to the closed ring assembly of RAD52, bacteriophage SSAPs Redβ from bacteriophage λ and RecT from a prophage of *Listeria innocua* (LiRecT) form left-handed helical filaments^29,30^. Both Redβ and RecT have been structurally captured with an extended and unwound duplex DNA intermediate that suggests a *cis* mechanism of annealing^29,30^. RAD52 does not form helical filaments, and whether it adheres to the *cis*-annealing mechanism observed for phage SSAPs is currently unknown.

Here, we turn to a distant RAD52 homolog from yeast called Mgm101. Mgm101 is more closely related in sequence to human RAD52 than to the phage SSAPs, sharing 17% sequence identity with RAD52, 15% identity with Redβ, and 13% identity with LiRecT over the DBD. However, as Mgm101 forms both rings and filaments like the phage SSAPs^31,32^, we hypothesize it may provide a key functional and evolutionary link between the two. *Mgm101* was initially discovered from a temperature-sensitive mutation that resulted in the loss of half of the mitochondrial DNA (mtDNA) at each cell division^33^. Mgm101 is a component of the mitochondrial nucleoid that is essential for mtDNA replication and repair of oxidative DNA damage^31,33,34^. Interestingly, Mgm101 is also present in the nucleus where it is involved in repair of interstrand crosslinks (ICLs) in the Fanconi-Anemia pathway, as well as telomere elongation in *rad52-*defective cells^35,36^. An in vivo assay demonstrated Mgm101 performs SSA within yeast mitochondria to mediate tandem repeat recombination^31^. In vitro, Mgm101 anneals naked ssDNAs as well as ssDNA pre-complexed with Rim1, the mitochondrial SSB protein^31^. Analytical ultracentrifugation (AUC), size-exclusion chromatography (SEC), and small-angle x-ray scattering (SAXS) suggested Mgm101 forms oligomers of ∼14 subunits in the absence of DNA^31,37^. Negative stain TEM shows Mgm101 forms both well-ordered rings and highly compressed helical filaments without DNA^31,37^. The latter accumulate upon storage of concentrated protein at 4 °C and are speculated to be an inactive storage form of the protein. Upon addition of ssDNA, Mgm101 forms clusters of heterogenous and poorly structured filaments^31^. To date, there are no high-resolution structures of Mgm101, leaving questions of its mechanism, assembly, and similarity to RAD52 and phage SSAPs unknown.

Here, we determined four cryo-EM structures of Mgm101, each representing a snapshot along a full SSA pathway. The structures include Mgm101 alone, and in complexes with ssDNA, a duplex intermediate of annealing, and B-form DNA product. Mgm101 forms a closed nonadecameric (19-mer) ring that binds the backbone of ssDNA within a narrow positively-charged groove, and base pairs the second complementary strand directly on top of the first within the groove to form an extended, unwound, and circular duplex intermediate. A fourth structure suggests that the annealed DNA product is then released from the groove as B-form DNA and captured by a novel positively-charged C-terminal β-hairpin motif at the upper and outer rim of the ring. We also observe lock-washer assemblies, but these do not contain DNA. The DNA-binding properties and structures captured by cryo-EM are further supported in solution by mass photometry (MP) and native mass spectrometry (nMS). Overall, our results reveal key structural snapshots along the Mgm101 SSA pathway including ssDNA capture, homologous pairing, and product binding, suggesting a unifying *cis* annealing mechanism for RAD52 family SSAPs.

## Results

### Solution Assembly of Mgm101 and its DNA Complexes

MP was used to investigate the solution assembly of Mgm101 alone and with DNA. MP correlates scattered light from single biomolecule collisions with a coverslip to determine mass. MP of Mgm101 at 100-500 nM monomer revealed a single species at 547-551 kDa that most closely matches a 19-mer, which would have a theoretical mass of 541 kDa (Fig. 1a and Supplementary Fig. 1). All mass measurements detected by MP, including replicates, are reported in Supplementary Table 1. The low mass region is not shown in Fig. 1 because the molecular weight of monomeric Mgm101 (28.4 kDa) is below the mass detection limit of MP (30-40 kDa) and the low mass region is susceptible to cumulative histogram artifact peaks^38,39^. Full histograms and replicates are shown in Supplementary Fig. 1.

**Fig. 1.**
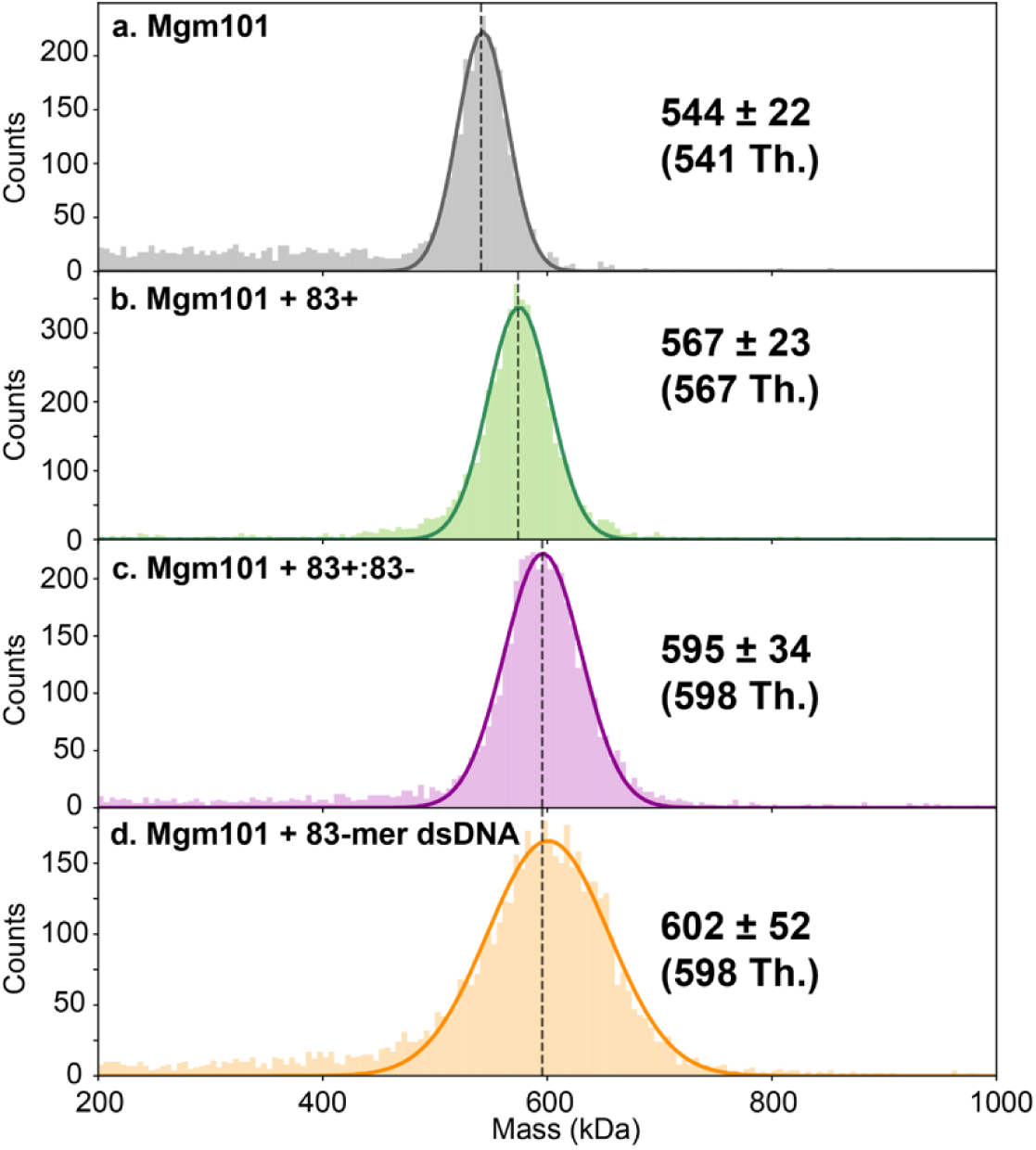
Mass photometry (MP) of Mgm101 alone and in DNA complexes reveals a 19-mer in all states. Example mass histograms of Mgm101 with no DNA **(a)**, 83+ ssDNA **(b)**, two complementary ssDNAs (83+:83-) added sequentially **(c)**, and pre-formed 83+/83- dsDNA **(d)**. The fitted mass and sigma for each peak are shown to the right, and the theoretical (Th.) mass is shown below. Each measurement was performed at 500 nM Mgm101 monomer with a stoichiometric amount of each DNA added (four nucleotides or bp per Mgm101 monomer). Each mass histogram is representative of three independent replicates, shown fully in Supplementary Figs. 1 and 2. Notice that MP can detect and quantify the mass shift as each strand of ssDNA is added, and that Mgm101 remains a 19-mer in all DNA-bound states.

MP was further used to investigate the assembly of Mgm101 with DNA. Assuming a stoichiometry of 4 nucleotides per Mgm101 monomer like RAD52^21^, we used an 83-mer oligonucleotide (83+) that was slightly more than long enough to occupy all 19 monomers of the Mgm101 oligomer (sequences of all oligonucleotides used in this study are provided in Supplementary Table 2). This produced a correspondingly larger species to match a complex of one 19-mer Mgm101 and one copy of 83+ (Fig. 1b and Supplementary Fig. 2a). We observed similar results with the slightly longer 87+ ssDNA (Supplementary Fig. 2b). When an equivalent amount of the complement strand (83-) was added to the pre-formed Mgm101 complex with 83+ (designated as 83+:83-), a species matching a complex of one 19-mer Mgm101 with two DNA strands was observed (Fig. 1c and Supplementary Fig. 2c). However, because the resolution in the 500-600 kDa range is around 66 kDa^38^, it is not discernible if the complex has two complementary strands (83+/83-) or two copies of the same strand (two 83+ or two 83-). Lastly, addition of pre-formed 83-mer dsDNA to Mgm101 resulted in a species matching a complex of one 19-mer Mgm101 and one copy of dsDNA (Fig. 1d and Supplementary Fig. 2d).

To examine the Mgm101 DNA complexes with higher mass accuracy, we employed nMS, which uses electrospray ionization (ESI) under conditions that maintain non-covalent interactions and oligomeric equilibrium near physiological ionic strength^30,40,41^. nMS of Mgm101 alone at 10 μM monomer produced a species clearly corresponding to a monomer, and two larger species corresponding to an 18-mer and a 19-mer (Fig. 2a). Experimental and theoretical masses for all species detected by nMS are reported in Supplementary Table 3. A concentration-dependent shift from monomer to 18/19-mer was observed, as samples at 20 and 1 μM Mgm101 produced lower and higher amounts of the monomer than at 10 μM, respectively (Supplementary Fig. 3). The 18-mer could conceivably arise from dissociation of a monomer from the 19-mer induced by the voltages used for desolvation, but tests with different solvent conditions and fragmentation methods suggest that the monomer and 18-mer are naturally present in solution (Supplementary Fig. 4).

**Fig. 2.**
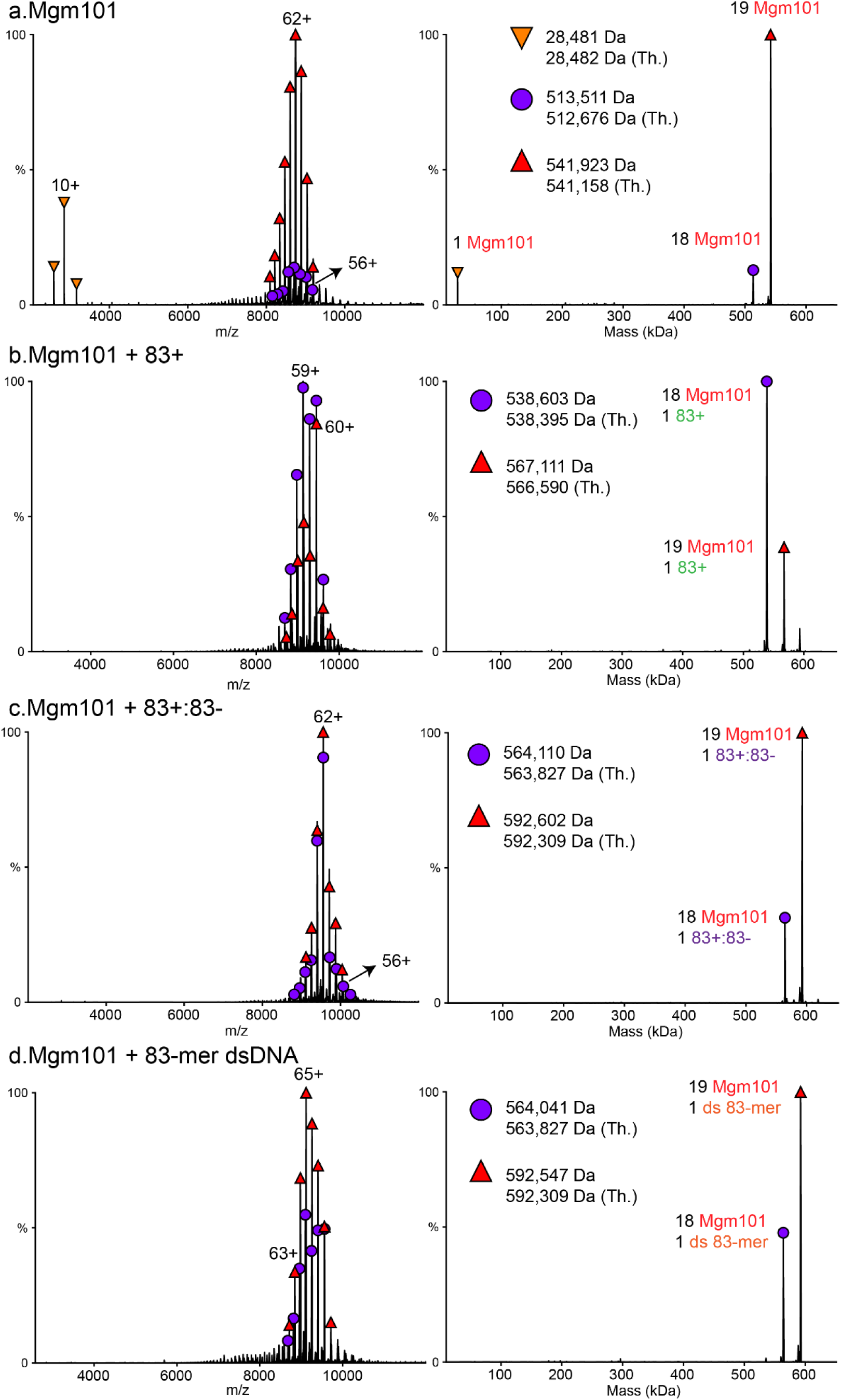
Native MS of Mgm101-DNA complexes. Representative raw native mass spectrum (left) and its corresponding deconvolved zero-charge mass spectrum (right) for (**a**) 10 μM Mgm101, (**b**) 10 μM Mgm101 with 40 μM nt 83+ ssDNA, (**c**) 10 μM Mgm101 with 40 μM nt 83+ and 83-ssDNA added sequentially, (**d**) 10 μM Mgm101 with 40 μM bp of pre-formed 83-mer dsDNA. The peak for each complex is labeled with its identified components based on matching the observed mass to the theoretical (Th.) mass of all possible complexes, as detailed by the symbols. Note that the observed masses are routinely slightly higher than the theoretical masses due to incomplete desolvation of folded species. Aside from this incomplete desolvation, the native MS measurements are accurate to within approximately 0.2% of the theoretical mass value.

We next performed nMS on 10 μM Mgm101 monomer mixed with a stoichiometric amount of 83+ ssDNA and again observed species consistent with 18 and 19-mer assemblies, each with a single copy of 83+ (Fig. 2b). Similar results were seen with four additional ssDNAs (87+, 83-, 75+, 75-) although for 75+ and 75-, additional peaks were observed for 18- or 19-mers with two copies of the ssDNA (Supplementary Fig. 5). The latter species could conceivably arise from attempts at annealing at sites of partial complementarity. When the complementary 83-strand was added to the pre-formed Mgm101-83+ complex, species corresponding to 18 and 19-mer assemblies with two strands were observed (Fig. 2c), supporting the conclusion from MP that the Mgm101 oligomer can form a *cis* complex with two complementary strands of ssDNA. This was confirmed by using complementary pairs of slightly different lengths to increase the mass difference (83-:87+, 87+:75-, and 75-:83+) (Supplementary Fig. 6), which unambiguously identified the complexes as having exclusively one copy of each strand. These observations are strongly suggestive of a *cis* annealing mechanism in which the two strands of ssDNA are paired on a single 18 or 19-mer Mgm101 oligomer. We further tested the concentration dependence of the *cis* annealing complex by performing the experiment at 10-fold lower concentration (1 μM Mgm101 monomer and 4 μM nucleotides of each ssDNA). The same 18-mer and 19-mer complexes with two complementary ssDNA strands were observed (Supplementary Fig. 7). We also performed controls in which two non-complementary ssDNAs, or a 2- or 3-fold excess of the same ssDNA were added to Mgm101 (Supplementary Figs. 8-10). This resulted in mixtures of multiple different species including complexes with only one ssDNA, or two copies of the same ssDNA. This was in stark contrast to the experiments with two complementary strands where a dominant species containing one copy of each strand was consistently observed. We also added a 2-fold excess (80 μM) of 87+ to Mgm101, followed by addition of one equivalent (40 μM) of the complementary 75-strand, to test if the complementary strand could displace the excess 87+ strand (Supplementary Fig. 11). Indeed, 18- and 19-mer complexes with two complementary strands (87+:83-) emerged as the dominant species. Finally, we performed nMS on 10 μM Mgm101 mixed with an equivalent amount (4 bp/monomer, or 40 μM bp) of pre-formed 83-mer dsDNA (Fig. 2d). Peaks corresponding to the 18- or 19-mer bound to one copy of the 83-mer were observed. Similar results were seen for the 75-mer pre-formed duplex (Supplementary Fig. 12). While the conformation of the dsDNA when bound to Mgm101 cannot be determined from this experiment, this result indicates the possibility of two distinct binding sites on Mgm101, one for ssDNA, and another for dsDNA.

In summary, the nMS data indicate that Mgm101 exists in a two-state equilibrium between a monomer and a large oligomer of 18 or 19 subunits. While the ratio of 18-mer to 19-mer varies across the different experiments (Supplementary Fig. 13), both species appear to bind and anneal DNA. No intermediate assemblies were observed (i.e., oligomers smaller than 18-mers) in either the presence or absence of DNA, nor were species of monomeric Mgm101 bound to DNA observed. The data are thus consistent with a cooperative process in which Mgm101 monomers assemble on ssDNA to form an 18 or 19-mer, which can then bind a second ssDNA strand to form a complex with two complementary strands.

### Cryo-EM structures of Mgm101

To visualize Mgm101 and its complexes with DNA, we used cryo-EM. We prepared three grids of Mgm101 incubated at 37 °C with stoichiometric amounts of: (1) 83+ ssDNA, (2) 75+ and 75-added sequentially (75+:75-), and (3) pre-formed 83+/83- dsDNA. From the first two grids we obtained structures of Mgm101 with ssDNA, and with a two-strand (duplex) DNA annealing intermediate, as expected. To our surprise, the second grid also yielded a structure with apparent B-form dsDNA product. The third grid did not yield a structure with dsDNA as expected, and instead resulted in two structures of apo (unbound) Mgm101: one as a closed 19-mer ring, and another as an apparent 18-subunit open-ring lock-washer. Single particle workflows for all structures are shown in Supplementary Figs. 14 to 16, and data collection and refinement statistics are shown in Supplementary Table 4.

In all DNA-bound states, Mgm101 forms a closed 19-mer ring that has a deep, narrow, and circular positively-charged groove around its circumference (Fig. 3a). As expected, Mgm101 adopts the same SSAP core fold seen in RAD52 and the phage SSAPs, consisting of a central pair of α-helices (α2-α3) flanked by a three-stranded antiparallel β−sheet (β3-β5) on one side and a β-hairpin (β1-β2) on the other (Fig. 3b and Supplementary Fig. 17). The α2-α3 helices form the base of the DNA binding groove, while the β3-β5 sheet and β1-β2 hairpin form its inner and outer walls, respectively. In contrast to the structural conservation of the core fold and DNA binding groove, Mgm101 also possesses a novel C-terminal β−hairpin motif (β6-β7) located above the outer rim of the DNA-binding groove, herein called the C-lobe (Figs. 3a and 3b)^21,29,30^. Residues 1-75 of Mgm101 are disordered in all structures, except for a short β-strand (β0) formed by residues 48-58 that packs onto β3 at the inner surface of the ring in the duplex intermediate structure, to extend the β3-β5 sheet inward (Supplementary Fig. 18).

**Fig. 3.**
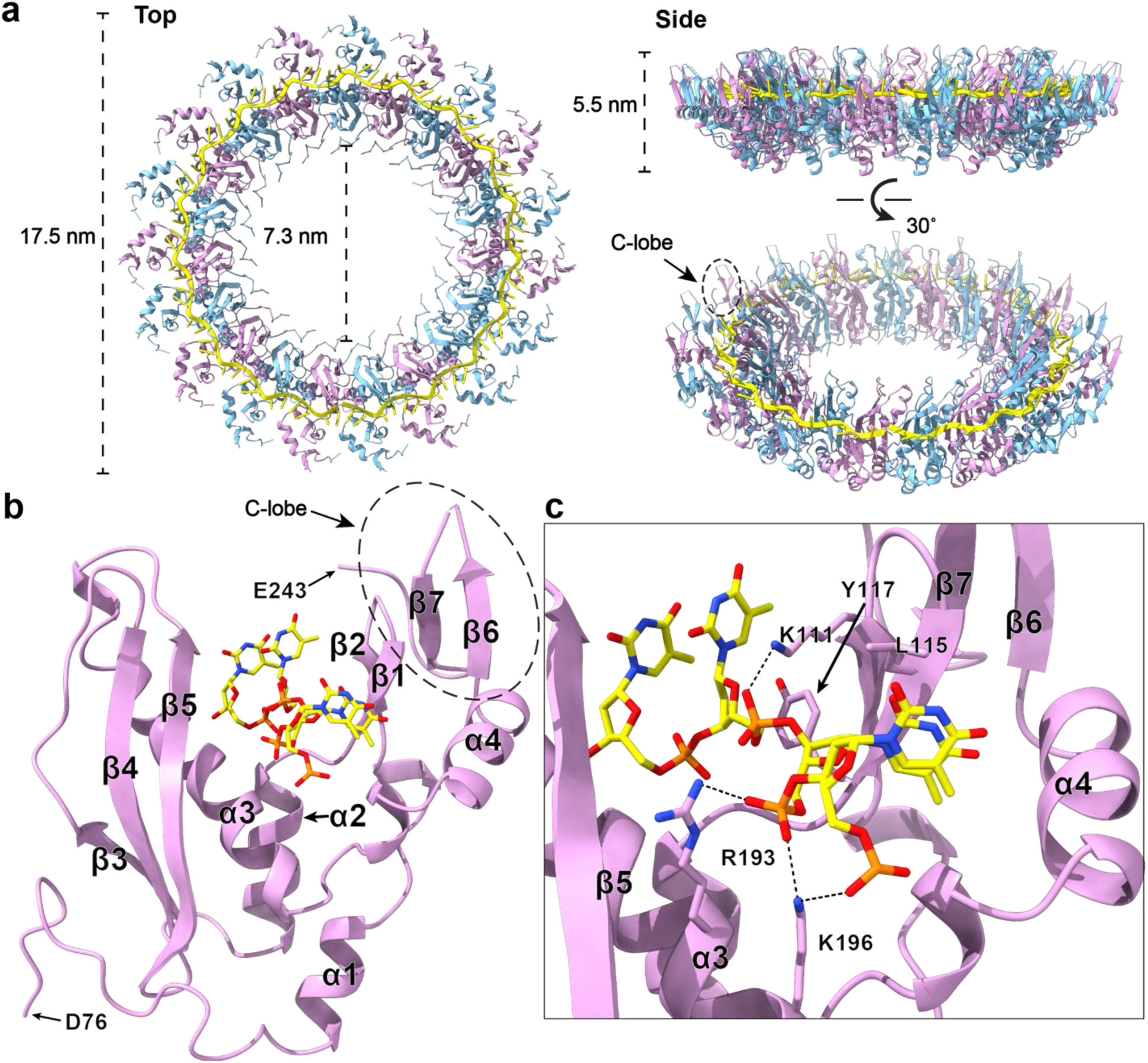
Cryo-EM structure of Mgm101 with 83+ ssDNA. **(a)** 19-mer Mgm101 ring (with alternating subunits in plum and cyan) bound to ssDNA (yellow) in top, side, and oblique views. The location of the C-lobe is shown within a dashed circle in the oblique view (only one is circled for clarity). **(b)** Mgm101 monomer bound to 4 nucleotides of ssDNA. The C-lobe is again depicted within a dashed circle, encompassing β6-β7 and the remainder of the C-terminus. **(c)** Close-up of (b), highlighting residues contacting the ssDNA.

In the complex of Mgm101 with 83+, the ssDNA is bound with its sugar-phosphate backbone buried deeply at the base of the groove atop α2-α3, and its bases facing upwards, exposed for homology recognition (Figs. 3b, 3c and Supplementary Fig. 19). The ssDNA adopts an extended repeating conformation consisting of segments of four nucleotides with close to B-form base stacking, disrupted by an irregular twist caused by insertion of the β1-β2 hairpin that separates the bases by 12 Å (Fig. 3c). The sugar-phosphate backbone is coordinated by a network of interactions including ion pairs with R193 and K196 from α3 and K111 from β2 (Fig. 3c). The side chains of several other residues, including E120 of α2, E184, S188, and N189 of α3, and W205 in the loop connecting α3 and α4 form H-bonding interactions with the phosphates. Two apolar residues of the β1-β2 hairpin, L115 (β1) and Y117 (β2) wedge into the DNA to stack with a base and contact a backbone ribose, respectively (Fig. 3c). All 10 of the residues that directly contact the ssDNA are among the most highly conserved residues in Mgm101 from ConSurf analysis (Supplementary Fig. 20).

From the grid containing Mgm101 incubated with 75+ and 75-sequentially, we obtained two structures: one of a complex with a duplex DNA annealing intermediate bound within the main DNA-binding groove (Supplementary Fig. 21), and another with apparent B-form DNA bound at the top of the ring to multiple successive C-lobes (Supplementary Fig. 22). The complex of Mgm101 with duplex annealing intermediate adopted essentially the same structure described above for the ssDNA complex, except for the appearance of clear density for a second ssDNA strand bound directly on top of the first (Figs. 4a and 4b, and Supplementary Fig. 21b). The first, or ‘inner’ strand is bound in essentially the same location and conformation as in the ssDNA complex. The second, ‘outer’ strand is fully Watson-crick (WC) base paired with the first, and adopts a similar extended, 4-nucleotide repeating conformation such that 4-nucleotide segments from each strand align to a backbone RMSD of 1.5 Å. The outer strand is bound at a slightly wider radius than the inner strand (Fig. 4a), indicating that its backbone conformation is slightly more extended. Together, the two strands form a fully base-paired duplex that is circular and completely unwound (Supplementary Fig. 21b).

**Fig. 4.**
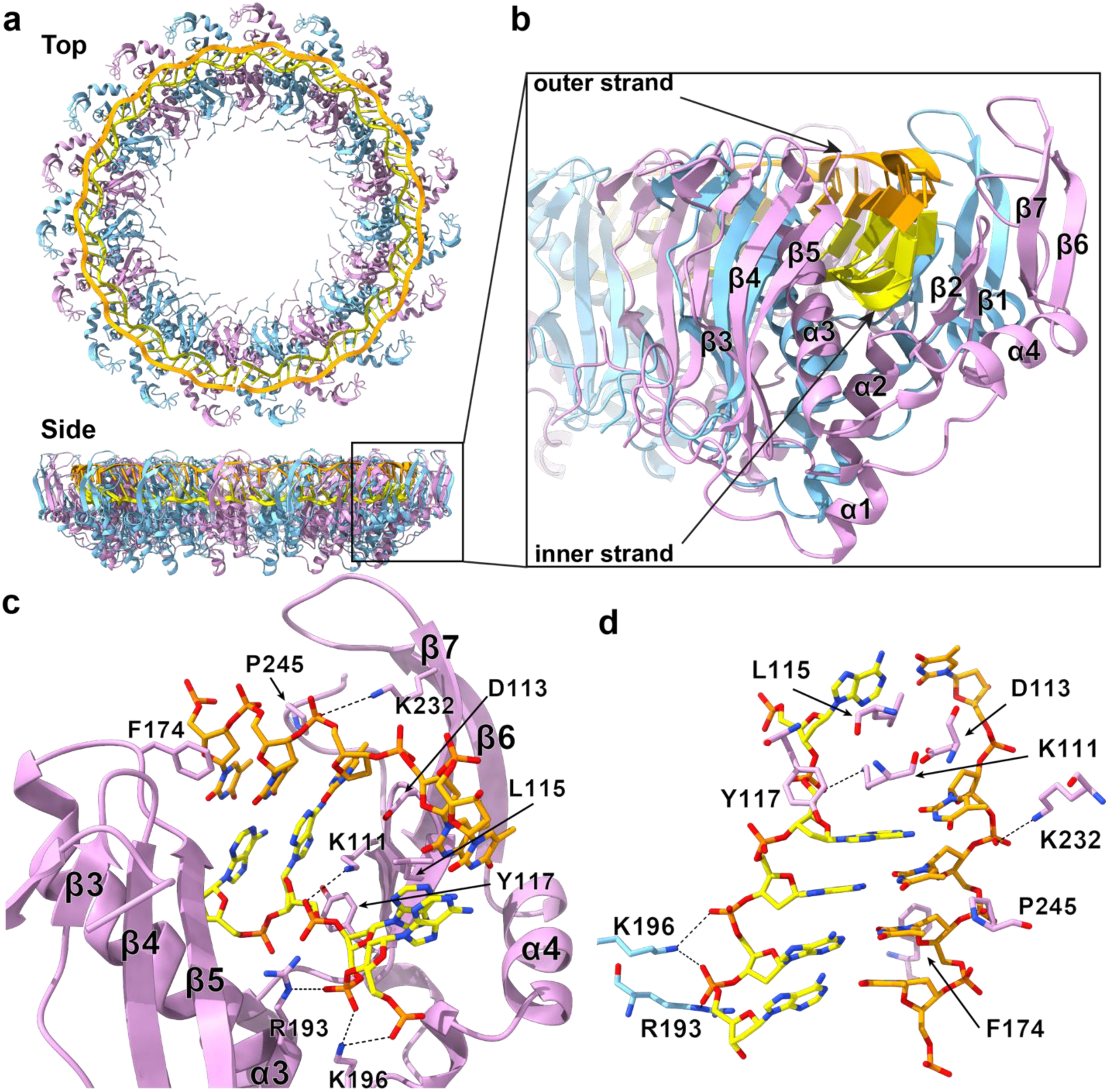
Cryo-EM of Mgm101 in complex with 75-mer duplex annealing intermediate. **(a)** Cartoon representation of the 19-mer ring of Mgm101 with alternating subunits in plum and cyan, 75+ ssDNA (added first) in yellow, and 75-ssDNA (added second) in orange. **(b)** Cutaway view looking into the narrow DNA binding groove. **(c)** Inner and outer strand Mgm101 interactions, with ion pairs and H-bonds shown as dashed lines. Notice that while the Mgm101 interactions with the inner strand are extensive (not all are shown for clarity), the interactions with the outer strand are sparse, such that it is held in the complex predominantly by WC base pairs with the inner strand. **(d)** 5 bp of the duplex intermediate (yellow and orange). Residues from the neighboring Mgm101 monomer are in cyan. As in the ssDNA structure, K111, L115, and Y117 from the β1-β2 hairpin wedge into the bases to distort the B-form conformation.

While the inner strand forms the same set of extensive interactions with Mgm101 described above for the ssDNA complex, the outer strand forms only limited interactions with the protein and is held in place almost exclusively by WC pairs with the inner strand. The closest Mgm101 interactions that do occur with the outer strand include D113 of the β1-β2 hairpin, which stacks against a base of the outer strand, F174 from the loop connecting β5 and α3, which packs against a ribose, K232 from β6, which forms a distant (∼ 5 Å) ion pair with a phosphate, and P245 near the C-terminus, which packs against the bases (Fig. 4c). Three of these residues, D113, F174, and P245 are highly conserved (Supplementary Fig. 20).

As was also the case for the ssDNA complex, the C19 symmetry averaging led to density corresponding to fully circular DNA with no discernable base identity or termini. Therefore, we modeled the inner strand as dA76, the outer strand as dT76, and arbitrarily placed the 5’-termini of each strand at the first and last subunits of the 19-mer. Although the inability to assign the sequence from the density for the bases complicates the conclusion that WC base pairs are formed (and thus that homologous pairing has occurred), the density clearly shows the opposing bases approach one another closely (Fig. 4d and Supplementary Fig. 21b), which can only occur if correct WC base pairs are formed.

The complex of Mgm101 with apparent B-form DNA also contained a 19-mer ring, but instead of having density for two DNA strands bound in the inner groove, it has weaker but distinct tube-shaped density resembling B-form dsDNA located above the DNA binding groove (Fig. 5 and Supplementary Fig. 22). This new density appears on only one side of the ring, where it contacts the C-terminal β6-β7 hairpins of ∼5 consecutive Mgm101 subunits (Fig. 5a). 35 bp of B-form dsDNA was modeled into this density at low map contour level to fit the overall shape, although the major and minor grooves are not clearly defined enough to fit with atomic-level precision. This presumed B-form DNA is close to a group of four consecutive positively-charged residues (K229-R230-K231-K232) on the loop connecting β6 and β7 (Fig. 5c), but due to the 6-8 Å resolution of this region of the map (Supplementary Fig. 22a), specific interactions cannot be clearly defined. While these four residues are only moderately conserved (Supplementary Fig. 20), a triple alanine mutation of K229, R230, and K231 was non-functional at 37 °C^42^.

**Fig. 5.**
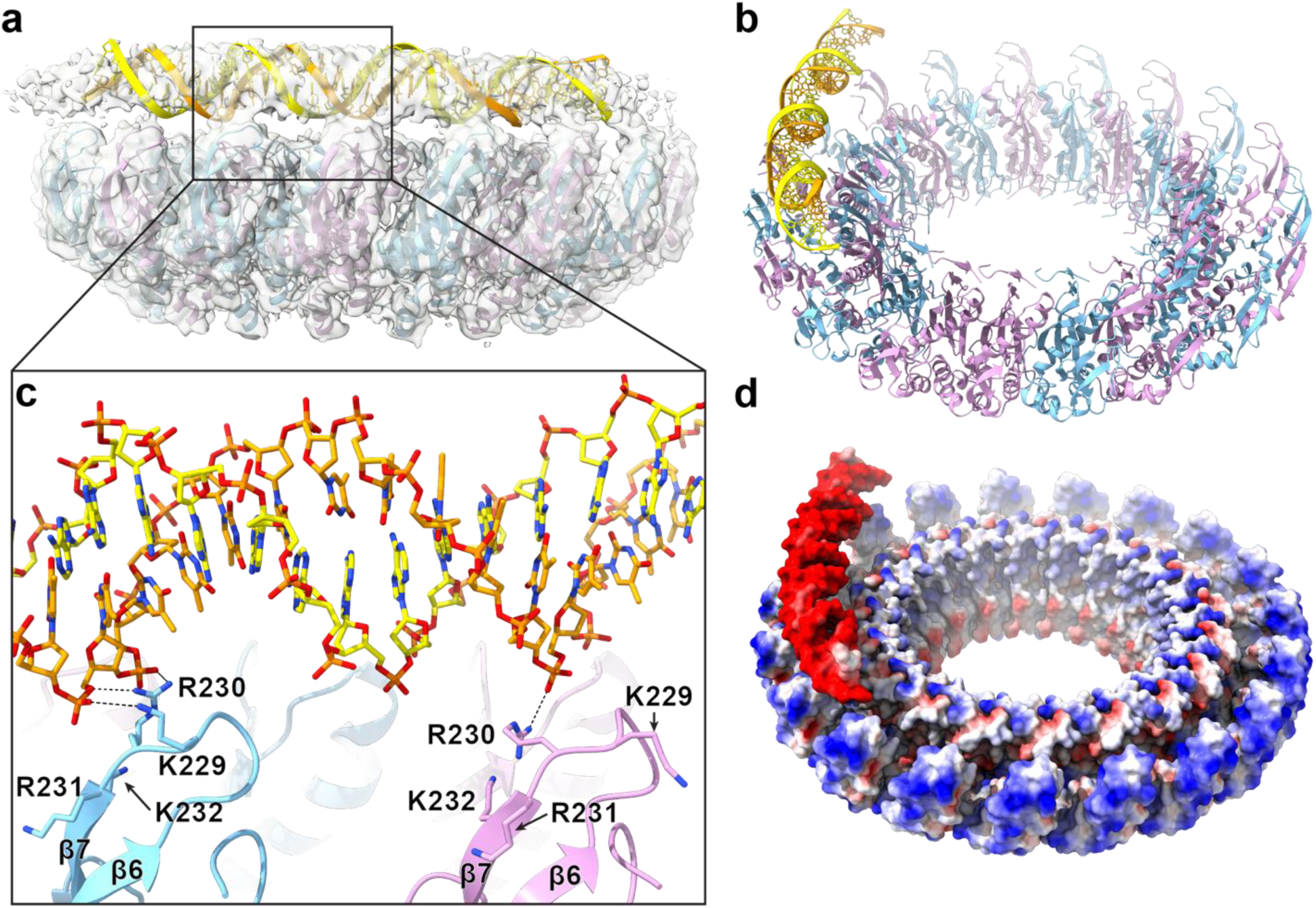
Structure of Mgm101 bound to apparent B-form dsDNA. **(a)** Side-view of the 19-mer Mgm101 ring (alternating subunits in plum and cyan) and 35 bp of dsDNA (orange, yellow) modeled into the tube-shaped density. **(b)** Oblique view of the complex rotated 90° with respect to (a). **(c)** Close-up view of the region boxed in (a), showing the proximity of the dsDNA backbone to the outer β6-β7 hairpins. Notice that K229 and R230 are poised to form potential ion pair interactions with the sugar-phosphate backbone and R231 and R232 are nearby. **(d)** Electrostatic surface representation of the complex shown in (b). The outer β6-β7 hairpins are abundant in positive charge (blue), both on the top and the side of the ring.

Although the functional significance of this apparent complex with B-form DNA has not been firmly established, it could result from annealed duplex intermediate dissociating from the inner DNA binding groove where the two strands were paired, folding into B-form DNA, and binding to an outer DNA binding site formed by the C-lobes. Along these lines, it is worth emphasizing that this structure did not contain DNA bound at the inner site.

While we hypothesize that the B-form dsDNA observed in the structure described above was annealed within the groove and subsequently captured at the outer site by the C-lobes, it is conceivable that dsDNA could either form in solution before binding to Mgm101, or anneal on the ring, fully disassociate, and then re-bind. Consistent with these possibilities, data herein from MP, nMS, and prior data from gel-shift assays^31^ show Mgm101 binds weakly to pre-formed dsDNA. To visualize dsDNA binding to Mgm101, cryo-EM samples were prepared by incubating Mgm101 with pre-formed 83-mer dsDNA for 10 minutes at 37 °C. The resulting cryo-EM images revealed two distinct types of particles: 19-mer rings like those described above (27.5% of particles refined to high-resolution) and an 18-subunit open lock-washer form (72.5% of final particles). Single particle analysis for each of these resulted in high resolution structures (Supplementary Fig. 16), but curiously neither contained bound DNA, either as ssDNA or dsDNA. Analysis of the 19-mer closed ring particles led to a 2.69 Å reconstruction to yield an apo structure (Supplementary Fig. 23). The apo ring structure superimposes onto the above structures with ssDNA and duplex intermediate to global RMSD values of 0.76 Å and 0.58 Å, respectively, indicating that no large-scale conformational changes in Mgm101 occur upon inner or outer strand binding.

Single particle analysis of the open lock-washer form (without symmetry averaging) led to multiple reconstructions after 3D-classification, two of which refined to 2.96 Å and 2.97 Å resolution (Supplementary Fig. 16). The two reconstructions appeared nearly identical, and so a full atomic model was built and refined for only the 2.96 Å reconstruction (Fig. 6 and Supplementary Fig. 24). The map showed strong density for 16 subunits at the central portion of the lock-washer, and weaker density for two additional subunits at the ends (Supplementary Fig. 24a). Even weaker density extending beyond the 18-subunit lock washer was observed, suggesting the possible presence of additional subunits to form a short but greater than 1-turn helix. However, inspection of the individual particles and resulting 2D classes showed only lock washers (Fig. 6c), suggesting that the extra density beyond the 18-subunit lock-washer likely arises from imperfect particle alignment (where single particle alignment of the apparent 18-subunit oligomer is off by a subunit). Continuous modeling by 3D variability analysis indicates the lock-washer undergoes a significant degree of motion (continuous disorder), particularly at its terminal subunits (Supplementary Movies 1-4).

**Fig. 6.**
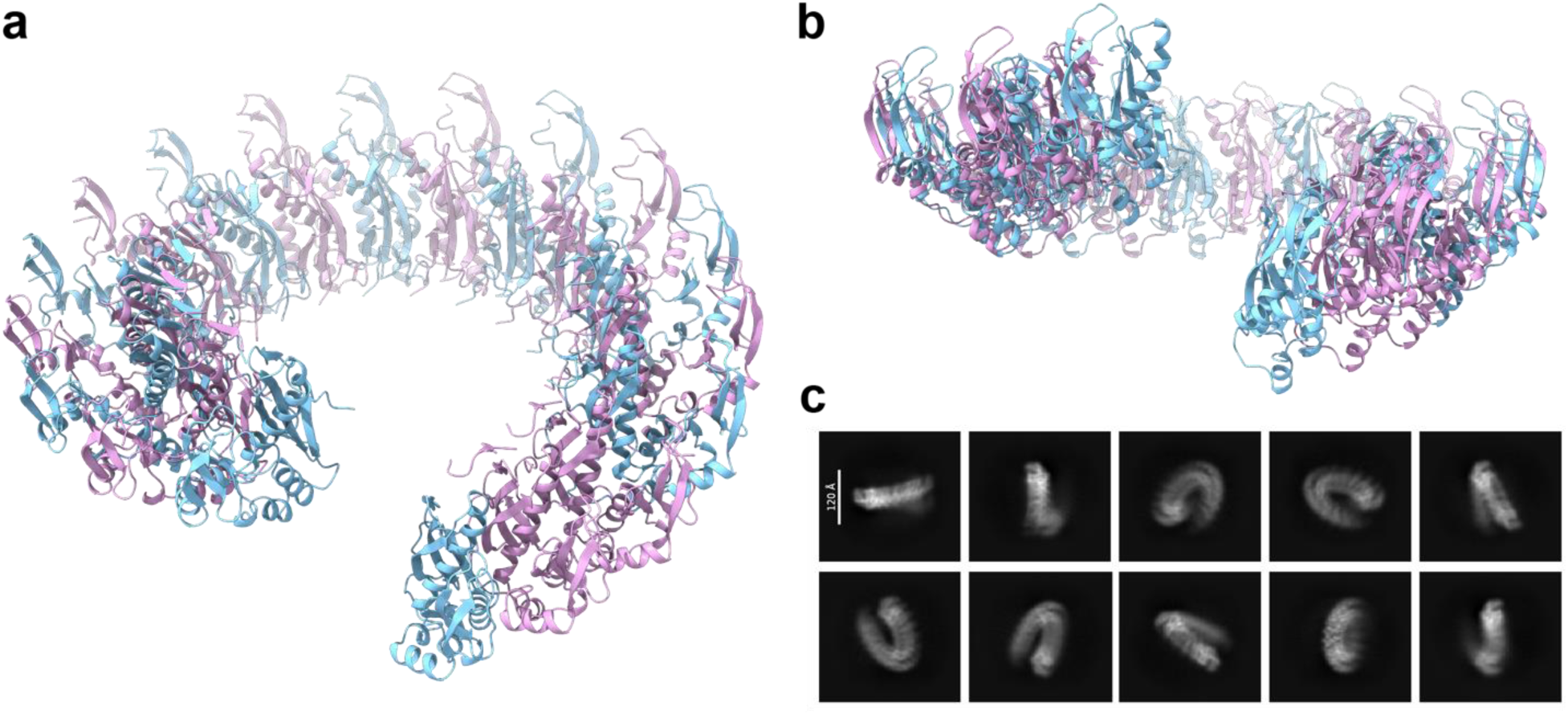
The open ring lock-washer structure of Mgm101. **(a)** Oblique view and **(b)** side view of the 18-subunit open lock-washer Mgm101 structure, where alternating protomers are colored in plum and cyan. **(c)** Selected 2D class averages of the particles corresponding to the lock-washer assembly.

Assuming the lock-washers do contain only 18-subunits, they could conceivably account for the 18-mers observed by native MS (all of the closed ring structures seen by cryo-EM contain exclusively 19 subunits). However, the 18-mers observed by nMS could bind DNA, whereas the 18-mer lock-washers observed by cryo-EM do not. Both the inner and outer DNA-binding sites are still available on the lock washer, but in slightly different, non-planar arrangements on adjacent subunits. Overall, the lock-washers contain a very similar structure and packing of the Mgm101 monomers as seen in the closed ring form, but apparently do not bind DNA, leaving their functional significance uncertain. The 18-subunit lock washers were also present in the data set collected for 75+:75-added sequentially (Supplementary Fig. 15), where again they did not contain bound DNA.

### Mgm101 DNA binding affinity and mutational analysis

For a mutational analysis to test the importance of residues making key interactions in the structures, we first developed a quantitative DNA-binding assay for wild-type (WT) Mgm101. To measure the affinity of Mgm101 for ssDNA, we performed fluorescence polarization (FP) titrations with three different 5’-fluoroscein-labeled ssDNA probes: a poly-dT 48-mer (dT48), a 48-mer of naturally occurring sequence (48+), and the 83+ oligo with a naturally occurring sequence used in nMS and cryo-EM (Fig. 7a). Mgm101 bound cooperatively to all three ssDNAs, with the highest affinity for 83+ (apparent *K_d_* = 360 ± 85 nM), and slightly weaker affinity for dT48 (apparent *K_d_* = 480 ± 130 nM) and 48+ (apparent *K_d_* = 540 ± 110 nM) (Table 1). The slightly higher affinity for dT48 compared to 48+ could arise from its lack of secondary structure. We also measured the affinity of Mgm101 for double-stranded versions of each of the three ssDNAs annealed with their respective complement strands (Fig. 7b). On average, the affinity for each dsDNA was 3.8-fold weaker than for the respective ssDNA, and the affinity for the longer 83-mer dsDNA was 2-fold higher than for the two 48-mers. Overall, this DNA-binding behavior is consistent with previous gel-shift assays of Mgm101 with 50-mer oligonucleotides that reported apparent *K_d_*values of 192 nM for ssDNA and 1.1 μM for dsDNA^31^.

**Fig. 7.**
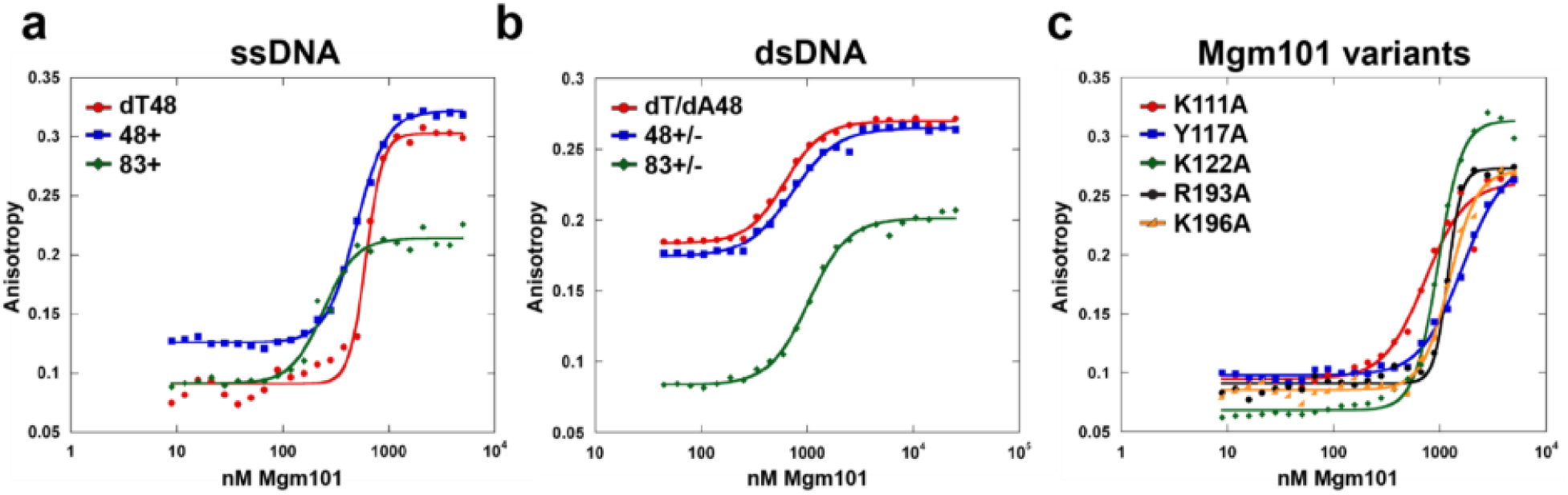
Binding of Mgm101 to different lengths of ss and dsDNA measured by fluorescence polarization (FP). 50 nM of 5’ Fluorescein labeled single stranded dT48, 48+, and 83+ (**a**), or double stranded dT/dA48, 48+/-, and 83+/- (**b**) was incubated with serially diluted WT Mgm101 at 8-5000 nM (a), or 44-25000 nM (b) at 25 °C for 30 minutes. (**c**) 50 nM of 83+ ssDNA was incubated with serially diluted site-directed variants of Mgm101 at 8-5000 nM. Fluorescence polarization was measured in triplicate at 25 °C.

**Table 1.** DNA binding affinity and Hill coefficients from fluorescence polarization titrations^1^.

| DNA | DNA type | Mgm101 variant | Apparent $K_d$ (nM) | Hill coefficient ($n$) | N |
| --- | --- | --- | --- | --- | --- |
| dT48 | ss | WT | $480 \pm 130$ | $4.2 \pm 1.2$ | 4 |
| 48+ | ss | WT | $540 \pm 110$ | $3.1 \pm 0.36$ | 5 |
| 83+ | ss | WT | $360 \pm 85$ | $3.7 \pm 1.1$ | 13 |
| dT48/dA48 | ds | WT | $2100 \pm 1500$ | $1.23 \pm 1.05$ | 3 |
| 48+/48- | ds | WT | $2200 \pm 1500$ | $1.5 \pm 0.64$ | 3 |
| 83+/83- | ds | WT | $1040 \pm 430$ | $2.8 \pm 1.2$ | 3 |
| 83+ | ss | K111A | $760 \pm 160$ | $2.6 \pm 0.81$ | 4 |
| | | Y117A | $1700 \pm 130$ | $1.9 \pm 0.33$ | 3 |
| | | K122A | $520 \pm 42$ | $3.4 \pm 0.28$ | 3 |
| | | R193A | $990 \pm 160$ | $4.1 \pm 1.8$ | 4 |
| | | K196A | $1200 \pm 100$ | $3.2 \pm 0.04$ | 3 |

<sup>1</sup>Apparent $K_d$ values are reported as the mean and standard deviation from at least three independent experiments (three separate titrations on different days). The far-right column (N) gives the number of replicates.

Based on the residues making key contacts with ssDNA in the structures, six site-directed mutants of Mgm101 were constructed and the variant proteins purified to measure their effects on DNA binding (Fig. 7c, Table 1). The mutations included K111A and Y117A at residues of the β1-β2 hairpin that wedge into bases, R193A and K196A at residues of α3 that contact the phosphates of the inner ssDNA strand at the base of the groove, and K122A of α2, which is on the outer surface of the Mgm101 ring, away from the ssDNA, to serve as a control. Lastly, F131A, at a residue buried in the hydrophobic core that is conserved with RAD52, was included as a negative control to test for protein folding and stability defects. All the mutated residues, including K122 and F131, are among the most highly conserved residues of Mgm101 (Supplementary Fig. 20).

As expected, the F131A variant could be purified as the MBP-fusion but was completely insoluble after proteolytic removal of the MBP tag, indicating a defect in folding and/or stability. The other five variants were fully soluble during purification and reached concentrations ranging from 4.2 to 132 mg/mL, suggesting that the mutations did not disrupt folding. The Y117A variant had the largest effect, decreasing ssDNA affinity by 4.7-fold. This residue is at the base of the β1-β2 hairpin and wedges into the DNA to stack with a ribose of the inner strand. The K196A and R193A variants had the next largest effects, decreasing affinity for ssDNA by 3.3- and 2.7-fold, respectively. This makes sense as these two positively-charged residues form close ion pairs with the phosphates of the inner strand. K111A of the β1-β2 hairpin reduced DNA binding affinity to a lesser extent, by 2.0-fold, which was somewhat unexpected as K111 wedges into the bases and contacts a phosphate. Lastly, the K122A variant had the mildest effects, reducing ssDNA binding by only 1.4-fold, consistent with it being distant from the DNA. Overall, the FP binding data is consistent with the structures, supporting the conclusion that K111, Y117, R193, and K196 are important for binding ssDNA. None of the single amino acid substitutions completely abolish ssDNA-binding, as the interaction involves a network of at least 10 conserved residues of the protein, such that simple truncation of each side chain (to alanine) is only partially disruptive.

### Correlation of Mgm101 structures with prior genetic and mutational data

In addition to the six structure-based mutations we tested above, a wealth of prior genetic and mutational data on Mgm101, informing on no less than 30 residues of the protein (Supplementary Table 5), can now be interpreted with a detailed structural framework. Early studies identified two temperature sensitive mutants, Mgm101-ts1 (P119S)^33^ and Mgm101-ts2 (D109N)^34^. P119, at the beginning of α2, sits right on top of one of the phosphates of the inner ssDNA, where the N-end of α2 forms a favorable charge-dipole interaction with the phosphate (Fig. 4a). The effects of the conservative D109N mutation are harder to rationalize structurally, as D109 is on the surface of β1 (opposite from the DNA) and makes only a loose ion pair with R240 of the C-lobe. Zuo *et al.* performed extensive deletion analysis and concluded that residues 77-241 of Mgm101, formed by N- and C-terminal deletions of 76 and 6 residues, respectively, are the minimal functional core that can complement the Mgm101-ts1 mutant^434243^. This agrees with our observation that D76 is the first ordered residue in the structure (aside from β0). They also performed extensive random mutagenesis of this core region and identified 16 mutations (at 15 different residues) that abolished activity in vivo. Although too numerous to detail here, a summary is provided in Supplementary Table 5, and essentially all the mutations can be easily rationalized by our structures, including four at residues that make key contacts with the inner ssDNA (G114D, Y117N, S188G, and K196E).

Mbantenkhu *et al.* were the first to purify large quantities of the Mgm101 protein and biochemically characterize it^31^. They tested three mutations based on conservation with RAD52, including N128A, F131A, and F231A, all of which strongly disrupted Mgm101 activity in vivo. N128 on α2 makes close H-bond interactions at the subunit interface. F131 at the end of α2 is buried deeply in the hydrophobic core. F213 of α4 is buried in the core at the base of the bottom of the ring. The N128A mutation was the least severe of the three, consistent with its being at a subunit interface as opposed to in a hydrophobic core. Nardozzi *et al.* mutated two vicinal cysteines, C194 and C195, finding that both are critical for function^37^. These residues of α3 are buried in the hydrophobic core, anchoring α3 so the adjacent R193 and K196 residues on either side are firmly positioned to contact the phosphates of the inner ssDNA.

Finally, in a later study, Mbantenkhu *et al.* mutated 10 residues of the C-lobe, and tested for activity both in vivo and in vitro, finding significant defects for most^42^. Four mutations with the largest defects (Group 1: K231A, W235A, R237A, Y246A), did not allow mtDNA maintenance in vivo. Another four of the mutations (Group 2: K229A, R230A, K248A, Y244A), affected mtDNA maintenance under stress conditions. The last two mutations (Group 3: F222A, K247A) had no defects in vivo. Three of the residues, K231 of Group 1, and K229 and R230 of Group 2, are positively-charged residues poised to contact the B-form DNA in our putative product complex. Two of the residues, Y246 of Group 1, and Y244 of Group 2, are within 5 Å of the outer ssDNA strand. The two Group 3 mutations are on the surface of the C-lobe, away from the DNA, consistent with their having no observed defects in vivo.

The authors purified the four Group 1 protein variants and found that R237A eluted in the void volume of an SEC column, K231 eluted at a larger size than WT, W235A eluted as a monomer, and Y246A eluted like WT. The authors tested for ssDNA-binding using a large circular ssDNA substrate, and found defects in ssDNA-binding, particularly for Y246A. Defects in ssDNA-binding are difficult to rationalize from the structure, as these residues of the C-lobe are distant from the inner ssDNA site. The generally positively-charged nature of the C-lobe could endow it with a weaker, non-specific affinity for the long ssDNA substrate. Interestingly, as a control the authors purified the Y117A protein variant, based on the conservation of this residue and known role for ssDNA binding in RAD52, and found that it surprisingly had normal affinity for the circular ssDNA substrate. Y117 directly contacts the 83+ and 75+ ssDNAs in our structures (at the inner site), and our purified Y117A variant had the strongest defects for ssDNA-binding in our FP DNA-binding assay. It may be that circular ssDNA substrates are not able to access the deep inner ssDNA-binding site on the Mgm101 ring. Altogether, the mutational data from this prior study focused on the C-lobe can largely be rationalized from our structures.

### Comparison with RAD52

Despite sharing only 17% sequence identity, the Mgm101 and RAD52 monomers align to an RMSD of 2.2 Å for 131 aligned pairs of Cα-atoms (Fig. 8a). Their DNA binding grooves are nearly identical in shape, size, and electrostatic charge distribution. At the base of the inner groove, Mgm101 coordinates the backbone of the inner strand in a similar manner as RAD52 (Fig. 8b), with ion pairs from R193A and K196A on α3 of Mgm101 conserved as R153 and R156 on α3 of RAD52 (Supplementary Fig. 25). RAD52 has an additional ion pair from K152 of its α3 that is not conserved in Mgm101. Both proteins use the β1-β2 hairpin to wedge into bases of the inner strand, where K111, L115, and Y117 of Mgm101 are structurally equivalent to R55, V63, and Y65 in RAD52. This wedging results in a 12 Å separation of bases in Mgm101, which is strikingly conserved as 11 Å in RAD52. The remaining 4-nt segments of ssDNA bound by Mgm101 and RAD52 retain a base separation and backbone conformation close to B-form DNA. Both proteins also use the same portions of their monomers to pack into similar ring-shaped oligomers, and many of the residues making key contacts at the inter-subunit interface are conserved (Supplementary Fig. 25).

**Fig. 8.**
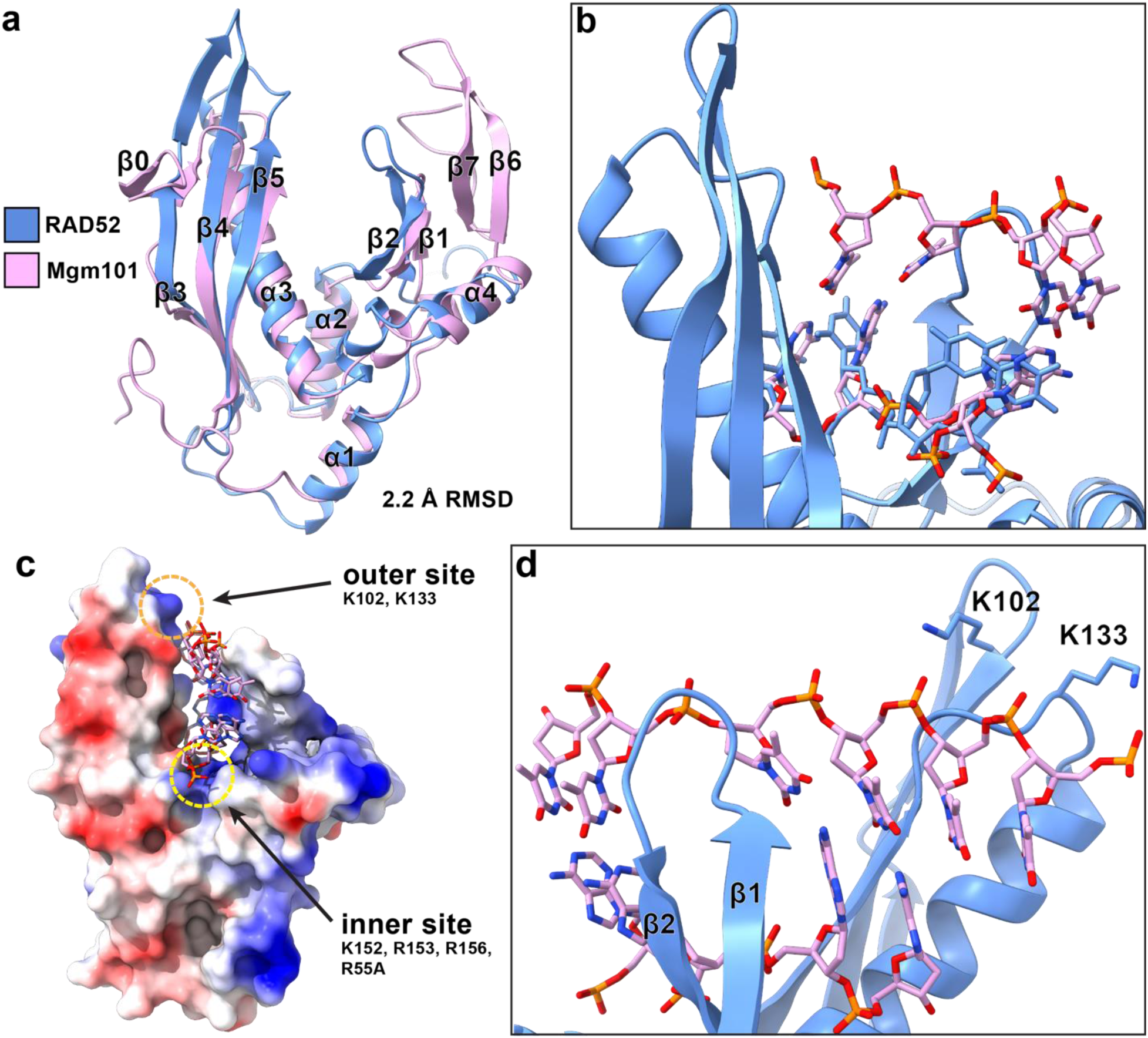
Structural alignment of Mgm101 with RAD52 suggests possible RAD52 interactions with a duplex intermediate. **(a)** Structure-based alignment of Mgm101 (residues 49-56 and 76-247, plum) and RAD52 (residues 25-208, blue). **(b)** RAD52-ssDNA complex (blue) superposed with 4 bp of annealed duplex DNA (plum) from the Mgm101 structure. **(c)** Electrostatic surface representation of monomeric RAD52 and the aligned duplex from (b), where the RAD52 inner and outer ssDNA binding sites are in yellow and orange dashed circles, respectively. **(d)** Rotated view of (c), showing how RAD52 outer site residues K102 and K133 could form ion pairs with phosphates of an outer strand.

**Fig. 9.**
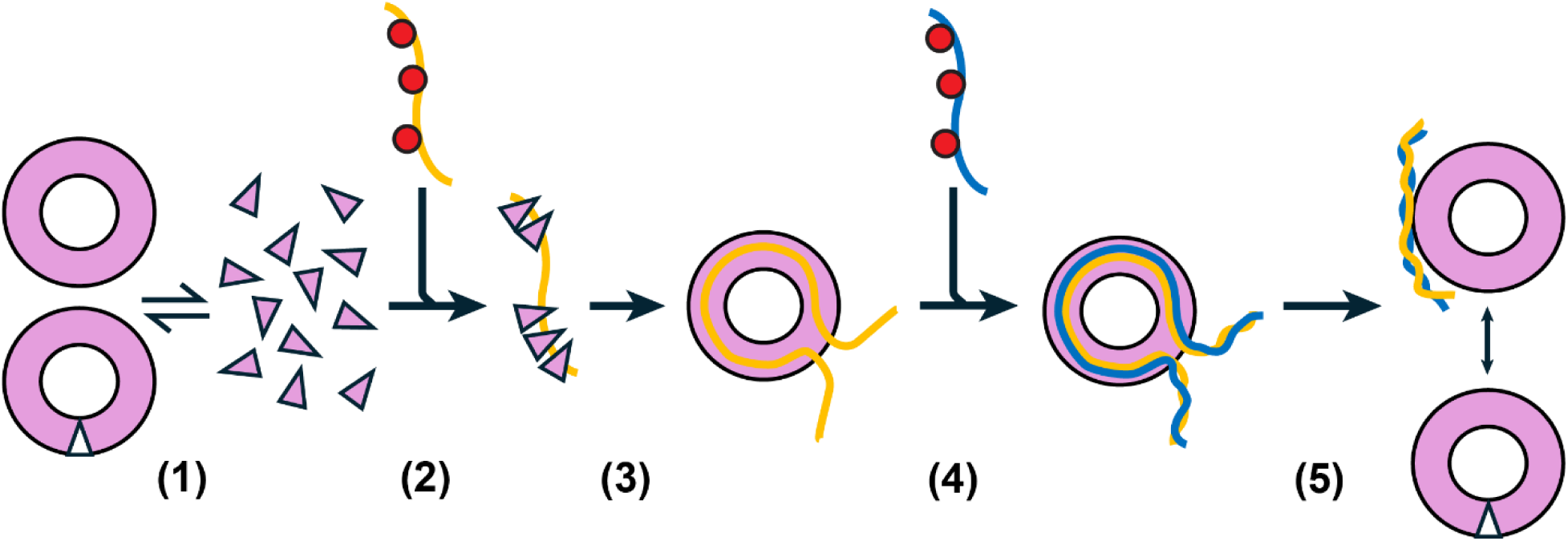
Model for single-strand annealing by Mgm101. **(1)** Mgm101 monomers are in equilibrium with 18/19-mers. **(2)** Mgm101 monomers assemble cooperatively on ssDNA, possibly coated with Rim1 (red circles) to **(3)** form a 19-mer ring-ssDNA complex. **(4)** Other ssDNAs can bind to the ring and sample the first ssDNA. When a match is found, a stable duplex intermediate is formed. Complementary ssDNA regions outside the groove are already aligned and free to fold into B-form dsDNA. **(5)** The paired ssDNAs within the groove eventually leave, fold into B-form dsDNA, and bind to the outer site at the Mgm101 C-lobe to drive the annealing reaction forward. 18-mer lock washers of Mgm101 without DNA are added to either end of the pathway, but their role is unclear. Native MS indicates that 18-mers can bind ssDNA in step 3, and complementary ssDNAs in step 4, but such 18-mers were not captured by cryo-EM. It is unclear if the 18-mers observed by native MS are closed rings or open lock washers.

Although RAD52 has not been captured with a second complementary strand as demonstrated here for Mgm101, a structural alignment shows that the inner ssDNA strand is bound in a highly conserved manner, and that the groove on RAD52 could easily accommodate a second strand, with minimal conformational changes at the tips of the β3-β5 sheet and β1-β2 hairpin (Fig. 8b). Residues of RAD52 that could potentially contact the second strand include K102 (between β3 and β4) and K133 (between β5 and α3) (Fig. 8d). Mutations at each of these residues have previously been shown to reduce affinity for dsDNA, and both residues contact the short segment of helical ssDNA bound at the outer site of RAD52 in a crystal structure^21,44^.

Despite these similarities there are some significant differences. As noted above, the C-terminal β6-β7 hairpin that is poised to make key interactions with the B-form dsDNA in Mgm101 is not present in RAD52, which instead has at the C-terminal end of its DBD an unstructured 20 amino acid linker followed by an α5 helix that packs against α2 of the neighboring subunit. While RAD52 has a largely disordered 200 amino acid C-terminal region for interacting with partner proteins, Mgm101 has no additional residues beyond its structured DBD, and instead has a largely disordered N-terminal tail of 75 amino acids (Supplementary Fig. 18a and 18b). While RAD52 has an unstructured N-terminal tail of 24 amino acid residues, there is no detectable sequence similarity between the two. A short segment of the N-terminal tail of Mgm101 binds as β0 on the inside of the ring to form potentially stabilizing inter-subunit interactions, and no such interaction occurs in RAD52 (Supplementary Fig. 18b and 18c).

## Discussion

SSA is a fundamental biological activity used in all cells for multiple aspects of DNA replication and genome maintenance. SSA is catalyzed by a diverse class of proteins known as SSAPs. In general, SSAPs bind preferentially to ssDNA and anneal two complementary strands of ssDNA in an ATP-independent manner. To date, at least three distinct families of SSAPs have been identified: the DdrB family found in bacteria including *Deionococcus radiodurans*, the ICP8 family found in oncogenic dsDNA viruses, and the RAD52 family found in eukaryotes and bacteriophage. Proteins of the DdrB and ICP8 families have at their core an SSB-like OB fold for binding ssDNA^45,46^. Crystal structures of DdrB show it forms a pentameric ring that can bind ssDNA and stack face to face with another such ring-ssDNA complex to mediate annealing in *trans*^47^. However, the two ssDNAs were only partially annealed at short complementary segments, and later stages of annealing are uncertain. ICP8 has been structurally captured as a monomer^46^, as a double 9-mer ring^48^, and as a bipolar filament^49^. Which of these forms are relevant to annealing is unclear, as no high-resolution structures have been determined with two complementary ssDNAs to suggest a clear annealing mechanism.

The RAD52 family uses a different (non-OB) core fold and appears to be the largest and most diverse. Classification of RAD52 family members has been historically challenging due to limited sequence conservation^26,50–52^, but the cryo-EM revolution and powerful new structure prediction methods have helped map subtle phylogenetic relationships. Five distinct sub-families of RAD52 SSAPs have recently been proposed as Redβ, RecT, Erf, Sak3, and RAD52^26^. While structures of RAD52 with ssDNA have been determined, how RAD52 pairs and anneals the second complementary strand is currently unknown. Structures of phage SSAPs Redβ and RecT, which form left-handed helical filaments, show how the complement ssDNA strand pairs directly on top of the first within the same groove on a single oligomer in *cis* to form an extended and unwound duplex intermediate of annealing^29,30^. While the resulting annealing models are compelling, they have yet to be confirmed for RAD52, which forms closed oligomeric rings instead of continuous helical filaments.

Here, we have determined five cryo-EM structures of a RAD52 homolog from yeast mitochondria called Mgm101. The structures provide snapshots of a more complete SSA pathway than has so far been established for other SSAPs. Based on these structures and complementary oligomerization and DNA binding analysis from MP and nMS, we propose a complete mechanistic model for Mgm101 annealing. In the absence of ssDNA, Mgm101 exists in solution as monomers in equilibrium with 18- or 19-mer rings or lock-washers. At the lower concentrations likely to exist in vivo, Mgm101 is predominantly monomeric. When ssDNA is introduced, such as by end-resection of a DSB, Mgm101 monomers assemble on the ssDNA cooperatively to form a 19-mer ring-ssDNA complex. In this initial substrate complex, the first ssDNA is bound deeply within the narrow groove at the inner site via extensive interactions with its sugar-phosphate backbone, leaving its bases exposed for homology recognition. This initial substrate complex provides a template for interacting with and sampling other ssDNA in the cell, such as the resected ssDNA from the other end of the break, or the ssDNA at the lagging strand of a replication fork. As potential ssDNA sequences enter the narrow groove to test for pairing, they form few interactions with Mgm101 itself, such that stable binding is dictated almost exclusiverly by formation of correct WC base pairs with the first ssDNA. When a sequence match is found, a stable duplex DNA intermediate of annealing is formed, as seen in our structure with a 75-mer duplex, which is extended and fully unwound. The annealed duplex product must then eventually leave the DNA binding groove, which could simply be driven by the force of the strands favoring the B-form conformation, which is too wide to fit in the narrow groove. Alternatively, Mgm101, or another protein could somehow actively displace the strands. One possibility is suggested by our structure of Mgm101 with what appears to be B-form dsDNA product. As the cellular ssDNA substrates are likely to be considerably longer than the 75-mers that fit in the groove on the 19-mer ring, once the initial pairing and alignment is formed within the inner groove, the remaining ssDNA strands outside the groove would be juxtaposed in perfect alignment, and free to fold into B-form DNA. The newly annealed B-form DNA would then be well-positioned to bind to the outer C-lobe regions of Mgm101, which could sequester and stabilize the duplex product as it is formed. As this association is presumably weak compared to ssDNA-binding, the B-form DNA would eventually be released as the DSB repair pathway or replication proceeds.

While this model is consistent with the available data, several important questions remain. For example, what is the importance of the size of the ring (11-mer for RAD52 vs. 19-mer for Mgm101) or the type of structure formed (closed ring vs. filament vs. lock washer)? The full range of oligomers observed for RAD52 family SSAPs varies considerably, suggesting that the exact oligomer size is not fundamental to the annealing activity. Moreover, recent RAD52 data indicates that at endogenous protein concentrations (∼ 20 nM), it exists as a monomer that can bind and anneal DNA as short 3-5 subunit clusters as opposed fully formed 11-mer rings^22^. Similar studies suggest the same is true for the bacteriophage Redβ SSAP^53^. In this view, the large oligomers observed at higher experimental concentrations used for structure determination may not be the functional species, making the distinction between an 11-mer and a 19-mer ring, or a ring and a filament irrelevant. In the case of Mgm101, the assembly of the 18- or 19-mer appears to be highly cooperative, more so than for RAD52. Even at the lower (nM) concentrations used for MP, intermediate species (between the monomer and 18- or 19-mer) were not observed, either for the protein alone or with DNA. In general, the highly cooperative nature of the SSAP-ssDNA interaction, together with cellular crowding phenomena and other uncertainties, obscures the size of the functional oligomers in cells considerably.

Another important question concerns the role of the novel C-lobe at the upper and outer rim of the Mgm101 ring. Mutational data and sequence conservation suggest the C-lobe is indeed critical for function, both in vitro and in vivo^42^. Based on our structure with apparent B-form DNA, we hypothesize the C-lobe binds and stabilizes the dsDNA product as the annealing reaction progresses on longer ssDNA substrates. However, other roles are conceivable, including (1) electrostatic attraction and steering of ssDNA substrates into the inner groove at the initiation and/or pairing steps, (2) binding of the initial ring-ssDNA complex to dsDNA to promote D-loop formation, (3) binding to dsDNA regions of the replication fork, as recently seen for RAD52^54^, (4) association of Mgm101 with the mitochondrial DNA within the nucleoid, or (5) yet to be identified roles in rolling circle replication of mitochondrial DNA^55^, inter-stand crosslink repair^35^ or telomere maintenance in the nucleus^36^. Our current data do not distinguish among these possibilities, and multiple roles are possible. While some of these roles could explain why the C-lobe is not found in RAD52 or the phage SSAPs, RAD52 contains a similar secondary outer site for binding dsDNA, located at the tips of α3 and β3-β5^44^. RAD52 has indeed been shown to promote D-loop formation, as has a homolog of Mgm101 from *C. parapsilosis*^32,44^. Thus, the functions of the Mgm101 C-lobe could still be relevant to the entire RAD52 family of SSAPs but adopted by a different structural element of each protein.

A key question for all SSAPs is if the second strand of ssDNA that is paired with the first is free in solution or bound by an SSB. RAD52 interacts with RPA, and cryo-EM structures have captured a putative RAD52-RPA annealing complex, although the complex did not contain two strands of ssDNA paired^9^. For bacteriophage Redβ, in vivo annealing at the lagging strand of the replication fork requires association with the host (*E. Coli*) SSB protein^28^. For Mgm101, yeast mitochondria contain an SSB called Rim1 that is closely related to bacterial SSBs^56^. Mgm101 can anneal ssDNA coated with Rim1^31^, but a direct protein-protein interaction has not yet been demonstrated. For RAD52 and Redβ, the interactions with RPA and SSB occur within a region that is C-terminal to the conserved DBD. Mgm101 does not have a C-terminal region beyond its DBD and instead contains a largely disordered N-terminal tail of 75 amino acids. The N-terminal tail of Mgm101 is required for in vivo activity but its sequence is not conserved. Further studies are needed to demonstrate if a physical Rim1-Mgm101 interaction occurs, and if so the regions of each protein involved.

While the N-terminal tail of Mgm101 is a likely candidate for Rim1 binding, it seems to also be required for folding and/or oligomer stabilization. Deletion of the N-terminal 75 residues eliminate function in vivo^43^, and attempted purification of the 75-residue N-terminally truncated protein led to insoluble protein (data not shown). Our structures indicate that a short segment of the N-terminal region forms a beta strand (β0) that packs onto the inner surface of the ring to a site that bridges neighboring subunits. The geometry of this interaction would indicate that the inner hole on the ring is occupied with the remaining disordered portions of the N-terminal regions of the 19-subunits. Further studies are needed to uncover the role of the N-terminal tail in the context of this unusual structural arrangement.

nMS experiments indicated the presence of both 18-mer and 19-mer forms of Mgm101 that could bind and anneal DNA. An open ring or lock-washer form of Mgm101 that appears to consist of 18 subunits was observed in both the annealed duplex DNA and pre-formed dsDNA cryo-EM datasets. While the lock-washer could explain the 18-mers observed by nMS, none of the lock washers observed by cryo-EM contained bound DNA. Native MS can kinetically trap transient species upon transfer into the gas phase, including species that may not emerge as abundant in cryo-EM. Such examples have been demonstrated for other proteins^41,57–60^. Interestingly, an open ring 10-mer form of RAD52, one subunit short of the closed 11-mer ring seen in cryo-EM, was recently discovered and found to have higher annealing activity than the 11-mer, particularly for RPA-coated ssDNA^9^. It is interesting to speculate that the 18-mer form of Mgm101 could be analogous to the 10-mer open-ring form of RAD52, but further studies are needed to establish its functional capabilities.

In summary, our oligomerization and cryo-EM data clearly indicate that Mgm101 anneals two complementary DNAs by a *cis* mechanism in which the two DNAs are paired within the same deep and narrow groove on a single protein oligomer. The extended and unwound structure of the duplex DNA intermediate, and the proposed *cis* mechanism of annealing, are strikingly similar to the intermediates recently captured by cryo-EM for two bacteriophage SSAPs^29,30^. Although RAD52 has so far only been visualized with ssDNA bound to the inner or the outer site, the conformation of the ssDNA, and the shape of the deep and narrow groove, are closely similar to what is seen in Mgm101 and the phage SSAPs. Moreover, modeling of RAD52 suggests that the second ssDNA could be easily accommodated with its groove, paired on top of the first in a similar manner as for Mgm101. We therefore suggest that the Mgm101 annealing model presented above is likely to be conserved and unifying for the entire RAD52 family of SSAPs.

## Methods

Residues 23-269 of *Saccharomyces cerevisiae* Mgm101 (Genbank #: X68482.1) were expressed from a pMal-C2E vector as an N-terminal maltose binding protein (MBP) fusion with an intervening PreScission protease cleavage site, as described previously^31^. Residues 1-22 of the natively expressed protein contain the mitochondrial targeting sequence and are cleaved off during translocation^43^. We therefore use the residue numbering scheme according to the mature protein, Val-1 to Lys-247, following Zuo *et. al*, 2007^43^. The Mgm101-pMal-C2E plasmid was transformed into BL21-DE3-RIL *E. coli* cells and cultured at 37 °C in 4 x 1L LB to an optical density at 600 nm of 0.6 and induced with 0.2 mM IPTG. At 5 hours after induction the cells were harvested by centrifugation, resuspended in MBP Buffer (20 mM Tris-HCl, 50 mM D-Glucose, 1 mM EDTA, pH 7.5) and frozen at -80 °C. Cells were thawed and incubated with Protease Inhibitor Cocktail Set I (Pierce, cat #: 539131) and lysozyme (1 mg/mL) for 30 minutes on ice. After sonication on ice using a Branson Sonifier (5 x 1-minute cycles at output 6, 30% duty), NaCl was added to 500 mM, and the crude lysate was centrifuged at 48,400 g for 30 minutes. The resulting supernatant was loaded onto a 2 x 5 mL MBP-trap HP column (Cytiva, cat #: 28918779) at 0.5 ml/min and washed with 200 mL of MBP Buffer. Bound protein was eluted with a 200 mL gradient into MBP Buffer with added 10 mM maltose. Fractions containing eluted protein were analyzed by SDS-PAGE, pooled, and dialyzed at 4°C overnight into 4 L of Heparin Buffer (50 mM Tris-HCl, 200 mM KCl, 0.5 mM EDTA, 2mM β-mercaptoethanol, pH 7.5) with added 10% Glycerol. The dialyzed protein was collected, mixed with Pierce HRV C3 protease (Thermo Scientific, cat #: 88946) at a ratio of 1 unit/100 ng protein, and incubated at 4 °C for 48-72 hours. The cleaved protein was centrifuged at 10,000 x g for 10 minutes and loaded at 0.5 ml/min back onto the 2 x 5 ml MBP column equilibrated in Heparin Buffer. The MBP column flow-through fractions containing un-tagged Mgm101 were analyzed by SDS-PAGE, pooled, and dialyzed into 4 L of fresh Heparin Buffer at 4 °C overnight. The dialyzed protein was collected, centrifuged at 10,000 x g for 10 minutes, and loaded onto a 2 x 5 mL Heparin column (Cytiva, cat #: 17040701) equilibrated in Heparin Buffer. The column was washed with 20 mL of Heparin Buffer, and bound protein was eluted with a 200 mL gradient of Heparin Buffer with added 2 M KCl. Eluted fractions containing protein were analyzed by SDS-PAGE, pooled, and dialyzed at 4°C overnight into 4 L of fresh Heparin Buffer. The dialyzed protein was collected and concentrated to 20 mg/ml (Vivaspin 20, 10 kDa MWCO, Sartorius cat #: VS2002). Protein concentration was measured by O.D. at 280 nM using a Nanodrop and an extinction coefficient of 42,400 M^-1^ cm^-1^ which was calculated from the amino acid sequence (247 amino acids, 28.48 kDa)^61^. Concentrated aliquots were stored at -80 °C.

### Site-directed mutagenesis

Oligonucleotides used for mutagenesis were purchased in desalted form from Integrated DNA Technologies (Corralville, IA). Primer sequences are shown in Supplementary Table 2. Site-directed mutations were introduced into the pMal-C2E-Mgm101 expression vector using the QuikChange method (Agilent technologies) and confirmed by Sanger dideoxy DNA sequencing (Ohio State University Comprehensive Cancer Center). The resulting plasmids were transformed into BL21-DE3-RIL *E. coli* cells and expressed as described above. Mgm101 protein variants were purified as described above, except the second MBP column step (after PreScission protease cleavage) was omitted.

### Mass photometry

Mass photometry data were collected using a Refeyn TwoMP instrument and analyzed using Refeyn AquireMP software. Ultrapure water and isopropanol were used sequentially to clean the glass coverslips and silicone gaskets. Both coverslips and gaskets were then dried with pure nitrogen. A clean coverslip with a 6- or 8-well silicone gasket on top was placed on the oil immersion objective. Briefly, 20 μL of poly-lysine was added to the coverslip and incubated at room temperature for 1 minute, then washed out with ultrapure water. For all MP measurements, purified Mgm101 was thawed from -80°C and diluted into Heparin Buffer to the indicated concentration. For Mgm101 without DNA, 100 nM, 200 nM, and 500 nM Mgm101 were incubated at 37 °C for 15 minutes to reach equilibrium and directly measured by the buffer-free method. For the Mgm101 83+ complex, Mgm101 was incubated with a stoichiometric amount of 83+ at 37 °C for 15 minutes. For the complex of Mgm101 with duplex intermediate, an equivalent amount of the complementary 83-strand was added to the Mgm101-83+ complex, and incubated for an additional 15 minutes at 37 °C. All oligonucleotides used for biochemical and structural studies were purchased HPLC-purified from Integrated DNA Technologies and dissolved in ddH2O. Both complexes were prepared at 10 μM and then diluted to 500 nM for measurement by the droplet dilution method. Data processing was performed with Refeyn DiscoverMP software. Raw data were exported and replotted using a bin size of 5 kDa to generate the mass distribution. The data was fit to a Gaussian distribution using a Python script.

### Native mass spectrometry

Mgm101 was thawed from -80 °C and buffer exchanged into 200 mM ammonium acetate (AmAc, Millipore Sigma) using a 6-kDa cut-off spin column (Bio-Rad, Cat #: 7326221). All ssDNAs were purchased HPLC purified from Integrated DNA Technologies, dissolved at 100 μM (molecules) in ddH_2_O, and buffer exchanged into 200 mM AmAc by using 10 kDa mass cutoff microdialysis (Pierce, cat #: PI88260). For complexes of Mgm101 with ssDNA, 10 μM Mgm101 (monomer) was mixed with a stoichiometric amount (4 nt per monomer) of ssDNA (83+, 83-, etc.) and incubated for 15 minutes at 37 °C. For complexes of Mgm101 with two strands of ssDNA, an equivalent amount of the second ssDNA (complementary strand or non-complementary control) was added and incubated for an additional 15 minutes at 37 °C. For complexes of Mgm101 with pre-formed dsDNA, 10 μM of Mgm101 (monomer) was mixed with 0.5 μM of pre-annealed 75-mer or 83-mer dsDNA, and incubated for 15 minutes at 37 °C. The pre-formed dsDNA was formed by mixing equal amounts of each strand, incubating at 95 °C for 5 minutes, and cooling slowly down to room temperature.

After complexes were formed, native MS experiments were performed on a Q Exactive Ultra-High Mass Range Orbitrap (UHMR) (ThermoFisher Scientific) modified with a 3-cm surface-induced dissociation (SID) device. 5 μL of sample was loaded into an in-house pulled borosilicate filament capillary (OD 1.0 mm, ID 0.78 mm) using a Sutter P-97 micropipette puller. The nanoelectrospray ionization voltage was held at a constant value of ∼1.1 kVs for the entire acquisition. The MS settings used for all measurements are listed below and kept consistent throughout the entire acquisition unless specified elsewhere: capillary temperature: 250 °C; ion transmission: high m/z; detector optimization: low-range of in-source trapping voltages was optimized, and -90 V was used to obtain the most accurate mass measurement; trap gas flow: 7 (UHV readback shows approximately 10^-10^ mbar). All apo data shown were collected under in-source trapping - 50V and trapping gas setting 5 to better represent the abundance of monomers. The resolution was set to 6,520 and defined at 400 m/z. Data processing was performed with UniDec 7.0.2. All settings remain the same across all data processing: m/z range: 2000-12000, charge range: 1 to 100; mass range: 10000 to 650000; Peak FWHM (Th): 1; Peak Shape Function: Gaussian; Beta: 5.0; Charge Smooth Width: 1.0; Point Smooth Width: 1.0; Mass Smooth Width: 0.0.

### Cryo-EM sample preparation

For the Mgm101-ssDNA complex, 1.0 mg/mL Mgm101 (in Heparin Buffer) was mixed with a stoichiometric amount (4 nucleotides/monomer) of 83+ and incubated at 37 °C for 10 minutes. The complex was stored on ice and diluted to 0.5 mg/mL protein with Heparin Buffer just prior to grid application. Within a Vitrobot Mark IV chamber set to 4 °C and 100% humidity, 4 μL of the complex was applied to a Quantifoil R1.2/1.3 Au 300 mesh holey carbon film that had been glow discharged for 60 s at 20 mA in a Pelco easiGlow. The grid was blotted for 4 s at blot force 1 and plunged into liquid ethane. The grid was clipped and stored under liquid nitrogen until imaging.

For the complex of Mgm101with duplex intermediate, 10 mg/ml Mgm101 (in Heparin Buffer) was incubated with a stoichiometric amount (4 nt/monomer) of 75+ oligonucleotide at 37 °C for 10 minutes. An equivalent amount of the complementary 75-oligonucleotide was added and incubated for an additional 10 minutes at 37 °C, before storing on ice. Just prior to grid application, 1 μL of 16 mM CHAPSO (Anatrace, cat #: C317) was added to 3 μL of 10 mg/mL complex. The 4 μL sample was applied to the surface of a Quantifoil R1.2/1.3 Cu 300 mesh holey carbon film that had been glow discharged for 60 s at 20 mA in a Pelco EasiGlow. The Vitrobot conditions were identical to those used for the above grid, except for a 6 s blot time and a blot force of -2.

The complex of Mgm101 with pre-formed dsDNA was prepared as described above for the complex with 75-mer duplex intermediate, except that 10 mg/ml Mgm101 was mixed with a stoichiometric amount (4.3 bp/monomer, equating to one 83+/83-dsDNA per one 19-mer) of pre-formed 83+/83- dsDNA (instead of the two being added sequentially) and diluted to 5 mg/mL before adding CHAPSO to a final concentration of 2 mM. Grids were prepared as described above for the 75-mer duplex intermediate sample.

### Cryo-EM data acquisition

All movies were collected on a 300 kV Thermo Scientific Krios G3i microscope using the nanoprobe EFTEM, a Cs corrector, Gatan BioQuantum energy filter (slit width 20 eV), and a Gatan K3 direct electron detector in counting mode. All datasets were collected using EPU at defocus ranges of -0.5, -1.0, -1.5, -2.0 and -2.5 um. Movies were acquired at 81,000x nominal magnification in super-resolution mode, to result in a pixel size of 0.426 Å. The detailed collection parameters of each of the three datasets are listed in Table S4.

### Cryo-EM data processing of Mgm101 complex with 83+ ssDNA

A set of 2,723 images with applied 30° tilt and a set of 3,010 images at 0° tilt collected from the same grid were imported as separate jobs in CryoSPARC v4.6.2. Both sets were patch motion corrected with ½ Fourier cropping, and patch CTF estimated. Exposures were then curated, and CTF fits ≤ 6.0 Å were retained. Blob picker with a diameter range set to 100-175 Å selected 1,653,892 particles from the 30° tilt set, and 2,142,621 particles from the 0° tilt set. 2D classification of each set separately resulted in selection of 23,442 particles (30° tilt set) and 70,762 particles (0° tilt set). A decoy set of 91,234 ‘junk’ particles was separately selected from the 2D classification of the tilted particles. The 23,442 and 70,762 selected good particles from 2D classification were fed into Ab-initio reconstruction, and then into homogenous refinement with C19 symmetry averaging. The resulting refined volume, along with 3 volumes created from an Ab-initio reconstruction of the ‘junk’ particles, was fed into heterogenous refinement with the blob-picked particles from both the 30° tilt and 0° tilt sets. The particles from the best resulting class (162,590) were subjected to a final round of 2D classification, in which two classes were selected that clearly represented the 30° tilt and 0° tilt rings. These particles were then used for Ab-initio reconstruction with C19 symmetry averaging, followed by homogenous refinement, and non-uniform refinement (with minimization over per particle scales, and optimization of defocus and CTF parameter optimization) with C19 symmetry. Particles were then reference-based motion corrected and subjected to final rounds of homogenous and non-uniform refinement (using the same settings as non-uniform refinement just prior), again with enforcement of C19 symmetry for a final reconstruction of 2.54 Å. Orientation diagnostics indicated the final map was significantly anisotropic, with a sampling compensation factor (SCF) value of 0.208.

### Cryo-EM data processing of Mgm101 complex with 75+ and 75-added sequentially

Movies were imported into CryoSPARC v4.6.2, and patch motion corrected with an output Fourier crop factor of ½. Following patch CTF estimation, manual exposure curation was used to select images with CTF fits ≤ 6.0 Å. Blob picker selected particles within a diameter range of 150-200 Å, and the resulting particles were subjected to 2D classification. Selected classes were fed into ab-initio reconstruction. A low-pass filtered (15 Å) non-uniform refinement map of the ab-initio volume was used to generate templates for picking, which resulted in 2,187,892 particles. Following another round of 2D classification, 417,609 particles were fed into ab-initio reconstruction, resulting in one class with 197,736 particles. This volume, along with 3 other “decoy” ab-initio volumes created from junk particles of the 2D classification job, were fed into heterogenous refinement, with the 2,187,892 particles selected from template picking. One class with 824,881 particles was selected for homogenous refinement, followed by re-extraction of the particles at a box size of 352 pixels, and another round of homogenous refinement. The non-uniform refinement volume of the un-binned 824,881 particles was used as input for 3D classification for 10 classes at a filter resolution of 5 Å.

The reconstruction for the complex with duplex intermediate, corresponding to Class 2, had 106,146 particles and was subjected to homogenous refinement, followed by non-uniform refinement with minimization over per-particle scales (min. PPS) and optimization of per-particle defocus settings. Reference-based motion correction was used before another round of homogenous refinement and non-uniform refinement with min. PPS, optimization of per-particle defocus and per-group CTF parameters. Flex data prep removed particles with per-particle scale factors of 0.75 and below, resulting in 81,000 particles for another round of non-uniform refinement. The output was then used for 3D variability (4 modes, at a filter resolution of 3 Å), which were grouped into 4 clusters. The best cluster containing 40,943 particles was homogenously refined, followed by a final round of non-uniform refinement with C19 symmetry (with min. PPS, and optimization of per-particle defocus) for a final GSFSC resolution of 2.54 Å. DeepEMhancer^62^ was used to sharpen the final map for use in model building in refinement.

For the structure with apparent B-form DNA, class 8 of the 3D classification job, which contained 116,779 particles, was subjected to homogenous refinement and non-uniform refinement with min. PPS. After optimization of per-particle scale factors and per-group CTF parameters, the particles were reference-based motion corrected. Following another round of homogenous refinement and non-uniform refinement with min. PPS, and optimization of per-particle scale factors and per-group CTF parameters, flex data prep removed particles with scale factors below 0.75, and non-uniform refinement (the same settings as above) was completed before another round of 3D classification (89,000 particles, 10 classes at filter resolution of 4 Å). The best volumes and particles (88,613 particles total) were grouped together for homogenous refinement and were then non-uniformly refined before 3D variability (3 modes, at a filter resolution of 7 Å). The modes were filtered into 3 clusters, the best of which (40,203 particles) was homogenously refined, followed by a final round of non-uniform refinement (same settings as above), for a final GSFSC resolution of 3.16 Å. DeepEMhancer was not used to sharpen the final map, as the sharpening process removed the density for B-form DNA.

For the lock-washer reconstruction, class 6 of the 3D classification, which contained 77,402 particles, was further processed. After homogenous refinement followed by non-uniform refinement with min. PPS, and optimization of per-particle scale factors and group CTF parameters on, particles were referenced-based motion corrected. The resulting 77,109 particles were non-uniform refined with the same settings as before, followed by Flex data prep with a cutoff of 0.75 for the scale factor. The final stack of 64,000 particles was subjected to a final round of non-uniform refinement, with the above settings, to a GSFSC resolution of 3.3 Å.

### Cryo-EM data processing of Mgm101 complex with preformed 83-mer dsDNA

Movies were imported into cryoSPARC v4.6.2, patch motion corrected with a Fourier-cropping at ½, and patch CTF estimated. Exposures were curated, and images with CTF fits ≤ 6.0 Å were kept for further processing. From an initial subset of images, templates were generated by blob picking and 2D classification and used to pick from the curated exposures. An initial 6,910,292 particles were extracted and 2D classified, from which 2,156,659 particles were further selected and fed into ab-initio reconstruction. The best-looking ab-initio class, which contained 1,570,915 particles, was subjected to homogenous refinement. This volume and 3 other “decoy” volumes generated by an ab-initio reconstruction from junk particles within the initial 2D classification job were fed into heterogenous refinement, with the entire 6,910,292 stack produced from template picking. The best class contained 2,840,752 particles that were then homogenously refined, followed by non-uniform refinement with min. PPS and optimization of per-particle defocus. The entire stack was then 3D classified into 10 classes at a filter resolution of 5 Å. Classes 1 (631,759 particles), 5 (470,314 particles), and 9 (606,716 particles) were each processed identically with the following (in order): homogenous refinement, NU-refinement (min. PPS, optimize defocus, optimize CTF), reference-base motion correction, another round of homogenous and a final round of NU-refinement (same settings as before). Class 5 and 9 were additionally subjected to flex data prep to remove particles with scales > 0.75, and then non-uniformly refined (class 5 was refined with C19 symmetry averaging). The final maps had resolutions of 2.97 Å (class 1, lock washer), 2.69 Å (class 5, apo 19-mer ring, with 383,000 particles) and 2.96 Å (class 9, lock washer, with 493,000 particles). The final maps of the lock-washer (class 9) and apo 19-mer ring (class 5) were sharpened with DeepEMhancer and were used for real-space refinement.

### Model building and refinement

For the ssDNA structure, an AlphaFold 3.0 prediction of monomeric Mgm101 was fit into the map, adjusted according to the observed density, and C19 symmetry expanded in ChimeraX (version 1.9) to complete the ring. The ssDNA was built and fit using Coot (version 0.9.8.93)^63^, and the model was real-space refined in PHENIX (version 1.21.2)^64^. The C19 ring from the Mgm101-ssDNA complex was used as a starting model for the duplex intermediate and apo-ring structures, while Mgm101 monomers from the ssDNA complex were individually fit into the lock-washer map in ChimeraX as a starting model. For maps produced with C19 symmetry averaging (complex with 83+ ssDNA, complex with 75-mer duplex intermediate, apo 19-mer ring) and the lock-washer reconstruction, NCS constraints were applied during Phenix refinement. For the map with B-form DNA, a 35-mer dsDNA (dA35/dT35) was built in Coot and manually fit into the density. Isolde^65^ was used to first refine the dsDNA conformation, followed by Phenix real-space refinement.

### Fluorescence polarization

For DNA binding measurements, 50 nM of 5’-fluoroscein-labeled ssDNA probe was incubated with serially diluted Mgm101 protein at room temperature for 30 minutes in PBS (137 mM NaCl, 2.7 mM KCl, 10 mM NaH_2_PO_4_, 1.8 mM KH_2_PO_4_, pH 7.4). To prepare the dsDNA probes, the 5’-fluoroscein labeled strand was mixed with an equimolar amount of the unlabeled complementary strand, and the mixture was incubated at 95 °C and cooled in stepwise increments of 2 °C every 2 minutes to a final temperature of 10 °C in a thermocycler. Fluorescence polarization was measured using a Tecan Spark plate reader at 485 nm excitation wavelength and 535 nm emission wavelength, in triplicate 20 μL aliquots in a Corning round-bottom black polystyrene 384 well plate (Corning, cat#: 3676) at 25 °C. Using Kaleidagraph (v4.03), mean fluorescence polarization (1/1000 mP) was plotted against protein concentration in nM and fit to the Hill equation to extract the dissociation constant and Hill coefficient as follows:

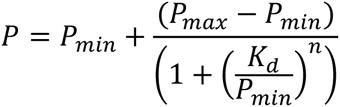

where P is the mean polarization (from triplicate measurements at each titration point), P_min_ is the fitted minimum polarization, P_max_ is the fitted maximum polarization, and n is the Hill coefficient. At least three such titrations were performed for each protein variant to determine values for the mean and standard deviation for the *K*_d_ and Hill coefficient (*n*) for each protein. All fits were subject to an R^2^ value ≥ 0.90 criteria to be considered for further analysis, and significant figures for the mean and standard deviation were considered to two digits.

## Reporting summary

### Data availability

The structural coordinates generated from this study have been deposited in the Protein Data Bank under accession code 9YI6 for the complex with 83+ ssDNA, 9YI7 for the complex with 75-mer duplex intermediate, 9YI8 for the complex with B-form dsDNA, 9YI9 for the unbound (apo) 18-subunit open lock washer, and 9YIA for the unbound (apo) closed ring 19-mer. The volumes generated in this study have been deposited in the EMDB data base under accession codes EMD-72979 for the complex with 83+ ssDNA, EMD-72980 for the complex with 75-mer duplex intermediate, EMD-72981 for the complex with B-form dsDNA, EMD-72983 for the unbound (apo) 18-subunit open lock washer, and EMD-72984 for the unbound (apo) closed ring 19-mer. The cryo-EM micrographs used in this study have been deposited in the EMPIAR database under accession code EMPIAR-13023 for Mgm101 mixed with 83+ ssDNA, EMPIAR-13025 for Mgm101 mixed with 75+:75-added sequentially, and EMPIAR-13024 for Mgm101 mixed with 83-mer pre-formed dsDNA. All unique biological materials, including plasmids used for protein expression, are available from the authors upon request.

## Supporting information

Supplementary Inforamtion

## Acknowledgments

The authors thank Dr. Xin Jie Chen of the State University of New York, Upstate Medical University, Syracuse NY, for kindly providing the Mgm101-pMal-C2E plasmid for protein expression. This work was funded by grants from the National Science Foundation (MCB-2212951 to C.E.B.) and the National Institutes of Health (T32GM141955 to C.T.W. and RM1GM149374 to V.H.W.). All cryo-EM data were collected at the Center for Electron Microscopy Analysis (CEMAS) at The Ohio State University, Columbus OH. The authors thank Yoshie Narui, Giovanna Grandinetti, and Binbin Deng for their assistance with cryo-EM data collection at CEMAS. The authors thank Joseph C. Sudar (NYU) for valuable discussions regarding cryo-EM data processing. The authors thank Sneha Kannan (OSU) for her assistance with the FP experiments. Molecular graphics and analyses performed with ChimeraX, developed by the Resource for Biocomputing, Visualization, and Informatics at the University of California, San Francisco, with support from National Institutes of Health R01-GM129325 and the Office of Cyber Infrastructure and Computational Biology, National Institute of Allergy and Infectious Diseases.

## Author Contributions

C.T.W., C.E.B, Z.Q, and V.H.W. designed research, C.T.W., Z.Q., K.Z and M.H. performed research, C.T.W., Z.Q., and M.H analyzed data, and C.T.W., Z.Q., V.H.W., and C.E.B. wrote the paper.

