## Supplementary Inforamtion for "Mechanism of single-strand annealing from native mass spectrometry and cryo-EM structures of RAD52 homolog Mgm101"

This document includes:

Supplementary Tables 1 to 5
Supplementary Figures 1 to 25
Captions to Supplementary Movies 1-4
SI References

**Table S1. Masses of Mgm101-DNA species identified by MP.** For each sample, the masses (in kDa) from three independent replicates are given with the sigma values from the fit. The mean and standard deviation from the three replicates are shown in the far right column. As the monomer is below the MP detection limit (30 – 40 kDa), the peak at the low mass region (Peak 1) could arise from cumulative histogram artifact peaks. Peak 2 is the Mgm101 oligomer shown in main text Fig. 1.

| Sample | Species | Rep 1 (kDa +/- σ) | Rep 2 (kDa +/- σ) | Rep 3 (kDa +/- σ) | Mean +/- S.D. (kDa) |
|---|---|---|---|---|---|
| **500 nM Mgm101** | Peak 1 | 63 ± 12 | 79 ± 14 | 61 ± 11 | 69 ± 8 |
| | Peak 2 | 557 ± 23 | 545 ± 30 | 544 ± 22 | 549 ± 6 |
| **200 nM Mgm101** | Peak 1 | 41 ± 8 | 45 ± 8 | 43 ± 8 | 43 ± 2 |
| | Peak 2 | 540 ± 16 | 548 ± 14 | 552 ± 14 | 547 ± 5 |
| **100 nM Mgm101** | Peak 1 | 58 ± 8 | 55 ± 9 | 50 ± 9 | 53 ± 2 |
| | Peak 2 | 548 ± 24 | 552 ± 18 | 554 ± 16 | 551 ± 3 |
| **500 nM Mgm101 with 83+ ssDNA** | Peak 1 | 65 ± 16 | 53 ± 15 | 63 ± 10 | 60 ± 5 |
| | Peak 2 | 559 ± 24 | 567 ± 23 | 570 ± 27 | 565 ± 5 |
| **500 nM Mgm101 with 87+ ssDNA** | Peak 1 | 65 ± 14 | 62 ± 16 | 76 ± 19 | 68 ± 7 |
| | Peak 2 | 585 ± 30 | 582 ± 27 | 577 ± 37 | 581 ± 4 |
| **500 nM Mgm101 with 83+:83-** | Peak 1 | 72 ± 15 | 93 ± 23 | 78 ± 19 | 81 ± 9 |
| | Peak 2 | 595 ± 34 | 594 ± 42 | 617 ± 39 | 602 ± 11 |
| **500 nM Mgm101 with 83+/83- dsDNA** | Peak 1 | 83 ± 21 | 97 ± 25 | 94 ± 27 | 91 ± 6 |
| | Peak 2 | 619 ± 44 | 602 ± 52 | 613 ± 47 | 611 ± 7 |

**Table S2. Sequences of oligonucleotides used in this study.**

| Oligonucleotide | Sequence (5’-3’) |
|---|---|
| 83- | TTGCATATTTAAAACATGTTGAGCTACAGCACCAGATTCAGCAATTAAGCTCTAAGCCATCCGCAAAAATGACCTCTTATCAA |
| 83+ | TTGATAAGAGGTCATTTTTGCGGATGGCTTAGAGCTTAATTGCTGAATCTGGTGCTGTAGCTCAACATGTTTTAAATATGCAA |
| 75+ | TTGATAAGAGGTCATTTTTGCGGATGGCTTAGAGCTTAATTGCTGAATCTGGTGCTGTAGCTCAACATGTTTTAA |
| 75- | TTAAAACATGTTGAGCTACAGCACCAGATTCAGCAATTAAGCTCTAAGCCATCCGCAAAAATGACCTCTTATCAA |
| 87+ | TTGATAAGAGGTCATTTTTGCGGATGGCTTAGAGCTTAATTGCTGAATCTGGTGCTGTAGCTCAACATGTTTTAAATATGCAATGAC |
| 87+ NC | GGTTATGTTTCAAGCGCACCTAATGCTAGAGTTATAGGTGGCAACTGCGACTCACGCCGCTAATCAGGCCGCACTTCCATAGCTCGA |
| Seq-F | GGTCGTCAGACTGTCGATGAAGCC |
| Seq-R | CGCCAGGGTTTTCCCAGTCACGAC |
| K111A-F | CTGGACCCTAAGGATATAGAAATCGCGCCAGATGGTTTGATATACTT |
| K111A-R | AAGTATATCAAACCATCTGGCGCGATTTCTATATCCTTAGGGTCCAG |
| Y117A-F | CTAAGGATATAGAAATCAAGCCAGATGGTTTGATAGCCTTGCCTGAAATTAAATAC |
| Y117A-R | GTATTTAATTTCAGGCAAGGCTATCAAACCATCTGGCTTGATTTCTATATCCTT |
| K122A-F | GCCAGATGGTTTGATATACTTGCCTGAAATTGCATACCGTAGAATACTGAA |
| K122A-R | TTCAGTATTCTACGGTATGCAATTTCAGGCAAGTATATCAAACCATCTGGC |
| F131A-F | ACCGTAGAATACTGAACAAAGCTGCTGGCGCAGGCGG |
| F131A-R | CCGCCTGCGCCAGCAGCTTTGTTCAGTATTCTACGGT |
| R193A-F | GCAAAAGTAATGCATTAATGGCGTGTTGCAAAGATCTCGGCG |
| R193A-R | CGCCGAGATCTTTGCAACACGCCATTAATGCATTACTTTTGC |
| K196A-F | GCATTAATGAGGTGTTGCGCAGATCTCGGCGTCGGTTC |
| K196A-R | GAACCGACGCCGAGATCTGCGCAACACCTCATTAATGC |
| 5’F 83+ | TTGATAAGAGGTCATTTTTGCGGATGGCTTAGAGCTTAATTGCTGAATCTGGTGCTGTAGCTCAACATGTTTTAAATATGCAA |
| 5’F 48 pT | TTTTTTTTTTTTTTTTTTTTTTTTTTTTTTTTTTTTTTTTTTTTTTTT |
| 48 pA | AAAAAAAAAAAAAAAAAAAAAAAAAAAAAAAAAAAAAAAAAAAAAAAA |
| 5’F 48+ | TGACGTAGATTCGAATCGTTAGTGAGAGCTTAATTGCTGAATCTGGTG |
| 48- | CACCAGATTCAGCAATTAAGCTCTCACTAACGATTCGAATCTACGTCA |

**Table S3. Masses of Mgm101-DNA species identified by nMS.**

| Figure | Observed mass (Da) | Expected mass (Da) | Error | Species |
|---|---|---|---|---|
| S3a | 28481 | 28482 | -1 | 1[Mgm101] |
| | 513427 | 512676 | 751 | 18[Mgm101] |
| | 541927 | 541158 | 769 | 19[Mgm101] |
| S3b | 28481 | 28482 | -1 | 1[Mgm101] |
| | 513550 | 512676 | 874 | 18[Mgm101] |
| | 542222 | 541158 | 1064 | 19[Mgm101] |
| S5a | 540105 | 539631.6 | 473.4 | 18[Mgm101]1[87+] |
| | 568670 | 568113.6 | 556.4 | 19[Mgm101]1[87+] |
| S5b | 538307 | 538108 | 199 | 18[Mgm101]1[83-] |
| | 566807 | 566590 | 217 | 19[Mgm101]1[83-] |
| S5c | 536337 | 535916.2 | 420.8 | 18[Mgm101]1[75+] |
| | 559616 | 559156.4 | 459.6 | 18[Mgm101]2[75+] |
| | 564856 | 564398.2 | 457.8 | 19[Mgm101]1[75+] |
| | 588094 | 587638.4 | 455.6 | 19[Mgm101]2[75+] |
| S5d | 535874 | 535646 | 228 | 18[Mgm101]1[75-] |
| | 558889 | 558616 | 273 | 18[Mgm101]2[75-] |
| | 564404 | 564128 | 276 | 19[Mgm101]1[75-] |
| | 587391 | 587098 | 293 | 19[Mgm101]2[75-] |
| S6a | 559121 | 558886.2 | 234.8 | 18[Mgm101]1[75+]1[75-] |
| | 587623 | 587368.2 | 254.8 | 19[Mgm101]1[75+]1[75-] |
| S6b | 565324 | 565063.6 | 260.4 | 18[Mgm101]1[83-]1[87+] |
| | 593815 | 593545.6 | 269.4 | 19[Mgm101]1[83-]1[87+] |
| S6c | 563468 | 562601.6 | 866.4 | 18[Mgm101]1[87+]1[75-] |
| | 591963 | 591083.6 | 879.4 | 19[Mgm101]1[87+]1[75-] |
| S6d | 561621 | 561365 | 256 | 18[Mgm101]1[75-]1[83+] |
| | 590119 | 589847 | 272 | 19[Mgm101]1[75-]1[83+] |
| S7 | 51153 | 51151 | 2 | 1[83+:83-] |
| | 564123 | 563827 | 296 | 18[Mgm101]1[83+]1[83-] |
| | 592679 | 592309 | 370 | 19[Mgm101]1[83+]1[83-] |
| S8a | 563750 | 563540 | 210 | 18[Mgm101]2[83-] |
| | 565121 | 564865 | 256 | 18[Mgm101]1[87NC]1[83-] |
| | 566468 | 566190 | 278 | 18[Mgm101]2[87NC] |
| | 568200 | 567915 | 285 | 19[Mgm101]1[87NC] |
| | 593627 | 593347 | 280 | 19[Mgm101]1[87NC]1[83-] |
| | 595002 | 594672 | 330 | 19[Mgm101]2[87NC] |
| S8b | 539987 | 539631 | 355 | 18[Mgm101]1[87+] |

| | | | | |
|---|---|---|---|---|
| | 559533 | 558616 | 917 | 18[Mgm101]2[75+] |
| | 563262 | 562601 | 660 | 18[Mgm101]1[87+]1[75+] |
| | 568553 | 568113 | 439 | 19[Mgm101]1[87+] |
| | 588072 | 587098 | 974 | 19[Mgm101]2[75+] |
| | 591762 | 591083 | 678 | 19[Mgm101]1[87+]1[75+] |
| S9a | 564650 | 564398.2 | 251.8 | 19[Mgm101]1[75+] |
| | 588185 | 587638.4 | 546.6 | 19[Mgm101]2[75+] |
| S9b | 536128 | 535916.2 | 211.8 | 18[Mgm101]1[75+] |
| | 559445 | 559156.4 | 288.6 | 18[Mgm101]2[75+] |
| | 564636 | 564398.2 | 237.8 | 19[Mgm101]1[75+] |
| | 587901 | 587638.4 | 262.6 | 19[Mgm101]2[75+] |
| S10a | 629478 | 628877 | 601 | 18[Mgm101]5[75+] |
| | 634562 | 634118.8 | 443.2 | 19[Mgm101]4[75+] |
| | 657909 | 657359 | 550 | 19[Mgm101]5[75+] |
| S10b | 593884 | 593542.8 | 341.2 | 18[Mgm101]3[87+] |
| | 622399 | 622024.8 | 374.2 | 19[Mgm101]3[87+] |
| S11 | 539787 | 539631.6 | 155.4 | 18[Mgm101]1[87+] |
| | 562997 | 562601.6 | 395.4 | 18[Mgm101]1[87+]1[75-] |
| | 568289 | 568113.6 | 175.4 | 19[Mgm101]1[87+] |
| | 591429 | 591083.6 | 345.4 | 19[Mgm101]1[87+]1[75-] |
| S12 | 559380 | 558886.2 | 493.8 | 18[Mgm101]1[75+:75-] |
| | 588039 | 587368.2 | 670.8 | 19[Mgm101]1[75+:75-] |

### Table S4. Cryo-EM Image Acquisition and Refinement Statistics

| **Dataset** | **83-mer ssDNA** | **75+/75- mer duplex intermediate and B DNA** | | **83-mer dsDNA** | |
|---|---|---|---|---|---|
| Micrographs | 3,010 – 0° tilt<br>2,723 – 30° tilt | 3,701 | | 8,199 | |
| EMPIAR ID | EMPIAR-13023 | EMPIAR-13025 | | EMPIAR-13024 | |
| Magnification | 81,000 X | | | | |
| Voltage (kV) | 300 | | | | |
| Total exposure ($e^{-}$/ Å$^2$) | 50 | | | | |
| Defocus ($\mu$m) | -0.5, -1.0, -1.5, -2.0, -2.5 | | | | |
| Pixel size (collection) (Å) | 0.426 | | | | |
| Pixel size (reconstruction) (Å) | 0.852 | | | | |
| **Structure** | **ssDNA** | **Duplex intermediate** | **Apparent B-form DNA** | **Lock-washer (Apo)** | **19-mer ring (Apo)** |
| PDB ID | 9YI6 | 9YI7 | 9YI8 | 9YI9 | 9YIA |
| EMDB ID | EMD-72979 | EMD-72980 | EMD-72981 | EMD-72983 | EMD-72984 |
| # Particles used in reconstruction | 162,066 | 40,943 | 40,203 | 493,000 | 383,000 |
| Map resolution (0.143 GSFSC, tight mask) (Å) | 2.54 | 2.54 | 3.16 | 2.96 | 2.69 |
| Symmetry | C19 | C19 | C1 | C1 | C19 |
| **Refinement Statistics** | | | | | |
| CC (mask) | 0.63 | 0.82 | 0.86 | 0.73 | 0.80 |
| r.m.s deviation bond lengths (Å) | 0.006 | 0.004 | 0.013 | 0.003 | 0.003 |
| r.m.s deviation bond angles (°) | 0.963 | 0.826 | 0.866 | 0.702 | 0.658 |
| MolProbity score | 0.84 | 1.50 | 1.93 | 1.47 | 1.62 |
| Clashscore | 1.20 | 9.48 | 23.96 | 8.66 | 7.35 |
| C$\beta$ outliers (%) | 0.0 | 0.0 | 0.0 | 0.0 | 0.0 |
| CaBLAM outliers (%) | 1.33 | 0.58 | 0.58 | 0.52 | 1.74 |
| Rotamer outliers (%) | 0.0 | 0.0 | 0.0 | 0.0 | 0.0 |
| Ramachandran favored (%) | 99.35 | 98.86 | 97.73 | 98.86 | 96.59 |
| Ramachandran allowed (%) | 0.65 | 1.14 | 2.27 | 1.14 | 3.41 |
| Ramachandran outliers (%) | 0.0 | 0.0 | 0.0 | 0.0 | 0.0 |

**Table S5. Summary of prior genetic and mutational data on Mgm101.**

| Mutation | Effects on Activity | Location in structure |
|---|---|---|
| | ***Mgm101-ts1, Chen et al., 1993*** [1] | |
| P119S | Failed to replicate mtDNA at 35 °C | At beginning of $\alpha$2, contacts a phosphate of inner ssDNA. |
| | ***Mgm101-ts2, Meeusen et al., 2003*** [2] | |
| D109N | Failed to replicate mtDNA at 35 °C | On surface of $\beta$1, forms loose ion pair with R240 of C-lobe. |
| | ***Zuo et al., 2007*** [3] | |
| L127P | Failed to complement Mgm101-ts1 (P119A) at 35°C in vivo | On $\alpha$2, buried in hydrophobic core. |
| E224G | Failed to complement Mgm101-ts1 (P119A) at 35°C in vivo | On $\beta$6, exposed on surface of C-lobe, near K229 and K247. |
| I179L | Failed to complement Mgm101-ts1 (P119A) at 35°C in vivo | At beginning of $\alpha$3, buried in apolar pocket near subunit interface. |
| G114D † | Failed to complement Mgm101-ts1 (P119A) at 35°C in vivo | At tip of $\beta$1-$\beta$2 hairpin, between D113 and L115, near inner ssDNA. |
| Y117N † | Failed to complement Mgm101-ts1 (P119A) at 35°C in vivo | On $\beta$1, at bottom of $\beta$1-$\beta$2 hairpin, packs against a ribose of inner ssDNA. |
| S188G † | Failed to complement Mgm101-ts1 (P119A) at 35°C in vivo | On $\alpha$3, forms H-bond with phosphate of inner ssDNA. |
| L198H † | Failed to complement Mgm101-ts1 (P119A) at 35°C in vivo | At rnd of $\alpha$3, buried in the hydrophobic core. |
| R124G † | Failed to complement Mgm101-ts1 (P119A) at 35°C in vivo | On $\alpha$2, near E120 and D197 at subunit interface. |
| K231E † | Failed to complement Mgm101-ts1 (P119A) at 35°C in vivo | On $\beta$6, exposed on surface of C-lobe (away from DNA). |
| R153W † | Failed to complement Mgm101-ts1 (P119A) at 35°C in vivo | On $\beta$3, near E154 across subunit interface. |
| K196E † | Failed to complement Mgm101-ts1 (P119A) at 35°C in vivo | End of $\alpha$3, forms close ion pair with phosphate of inner ssDNA strand |
| L99I † | Failed to complement Mgm101-ts1 (P119A) at 35°C in vivo | On $\alpha$2, buried in hydrophobic core. |
| L118S † | Failed to complement Mgm101-ts1 (P119A) at 35°C in vivo | At end of $\beta$2, buried in hydrophobic core, near inner ssDNA backbone. |
| | ***Mbantenkhu et al., 2011*** [4] | |
| N128A | Failure to replicate mtDNA at 37 °C, aggregation *in vitro* | On $\alpha$2 helix, H-bonds at subunit interface |
| F131A | Failure to replicate mtDNA at 35 and 37 °C, aggregation *in vitro* | Atend of $\alpha$2, buried in hydrophobic core. |
| F213A | Failure to replicate mtDNA at 35 and 37 °C, aggregation *in vitro* | On $\alpha$4, buried in hydrophobic core at bottom of ring. |
| | ***Nardozzi et al., 2012*** [5] | |
| C194A, C194A, C194D, C194E | Mutations destabilize mtDNA in vivo and lead to defects in ring formation or aggregation in vitro. | On $\alpha$3 helix, buried in hydrophobic core, near R193 and K196 that contact phosphates. |
| C195A, C195S, C195D, C195E | Mutations destabilize mtDNA in vivo and lead to defects in ring formation or aggregation in vitro. | On $\alpha$3 helix, buried in hydrophobic core, near R193 and K196 that contact phosphates. |
| | ***Mbantenkhu et al., 2013*** [6] | |

| | | |
|---|---|---|
| F222A | Group 3, no defects in vivo. | On s β6, surface of C-lobe, not near DNA. |
| K229A | Group 2, affected mtDNA maintenance under stress conditions. | At beginning of β7, surface of C-lobe, poised to contact B-form DNA. |
| R230A | Group 2, affected mtDNA maintenance under stress conditions. | At beginning of β7, 7.9 Å from outer strand, 5 Å from B-form DNA |
| K231A | Group 1, did not support mtDNA maintenance in vivo. | At beginning of β7, surface of C-lobe, close to B-form DNA. |
| W235A | Group 1, did not support mtDNA maintenance in vivo. | At end of β7, partially buried within C-lobe, not near DNA. |
| R237A | Group 1, did not support mtDNA maintenance in vivo. | At end of β7, buried into interface with DBD. |
| K248A | Group 2, affected mtDNA maintenance under stress conditions. | At bottom surface of C-lobe, contacts backbone C=O of D216 |
| Y244A | Group 2, affected mtDNA maintenance under stress conditions. | At C-terminal tail of C-lobe, within 4 Å of outer strand. |
| Y246A | Group 1, did not support mtDNA maintenance in vivo. | At C-terminal tail of C-lobe, within 4 Å of outer strand. |
| K247A | Group 3, no defects in vivo. | At C-terminal residues of C-lobe, exposed on surface away from DNA. |

†: The indicated mutation occurred together with one or more other mutations to give the indicated phenotype but was indicated by the authors to be the change likely to affect function based on sequence conservation analysis.

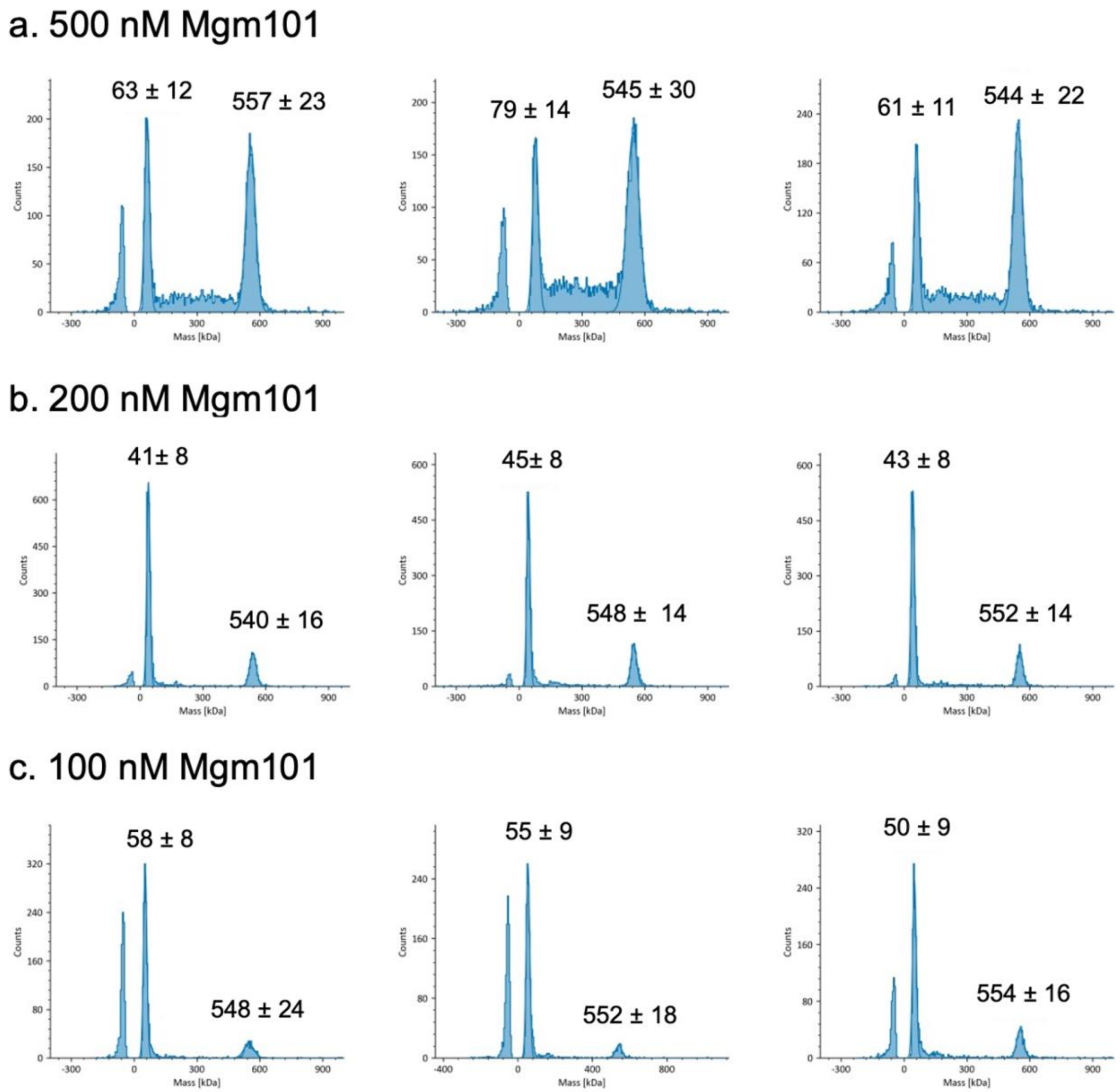


**Figure S1. Mass photometry of Mgm101 without DNA at different concentrations**. Three replicates are shown for each concentration. Each peak is labeled with the measured mass and fitting error (sigma) in kDa. From the three replicates, the mean and standard deviation of the mass of each species is reported in main text Table 1. The peaks to the left of zero kDa on the x-axis are the unbinding event. The theoretical masses of Mgm101 monomer and 19-mer are 28 and 541 kDa, respectively. As the monomer is below the MP detection limit (30 – 40 kDa), the peak at the low mass region could arise from cumulative histogram artifact peaks.

a. 500 nM Mgm101 with 83+

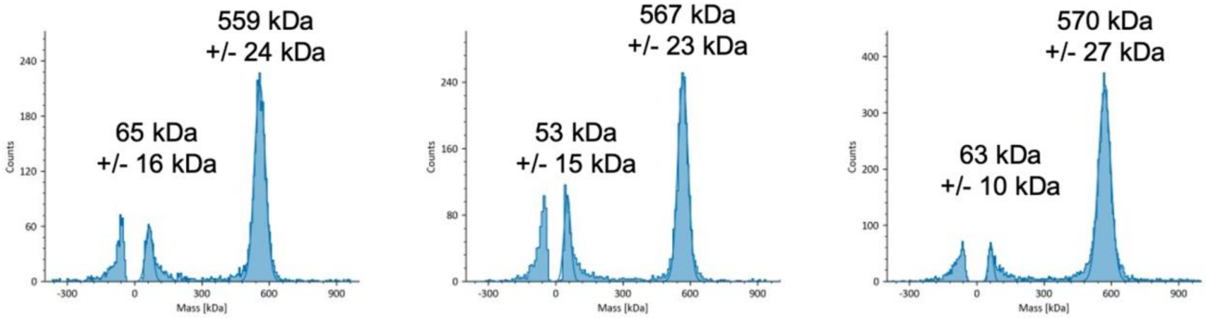


b. 500 nM Mgm101 with 87+

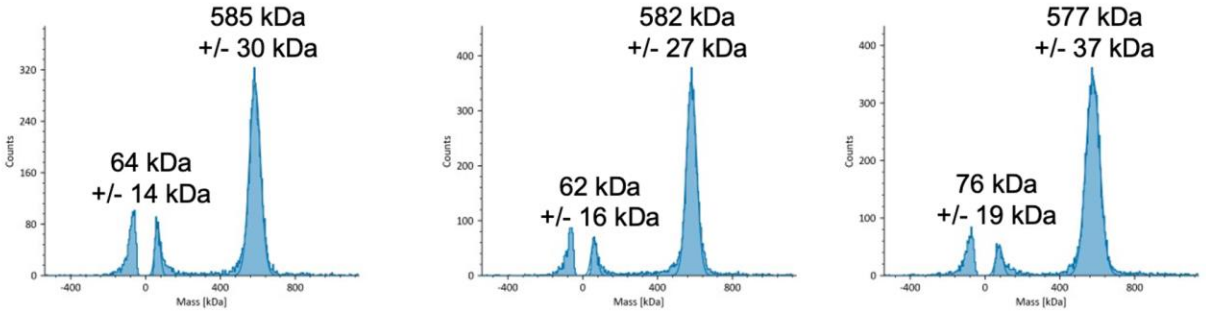


c. 500 nM Mgm101 with 83+:83-

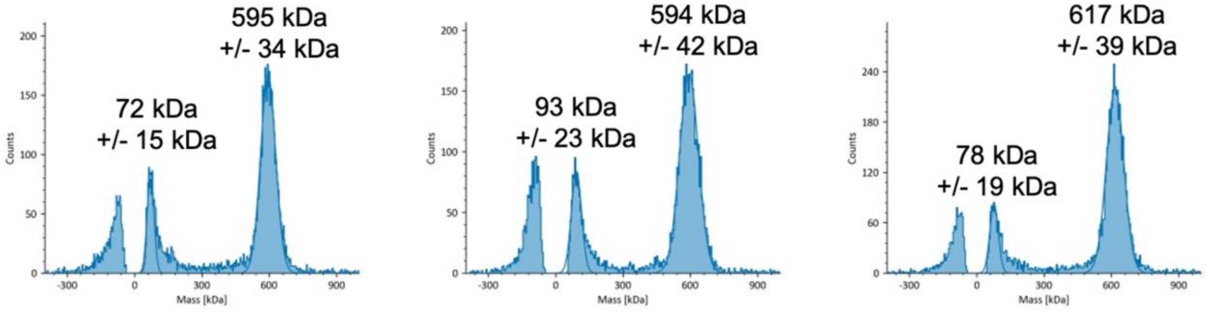


d. 500 nM Mgm101 with pre-formed 83-mer dsDNA

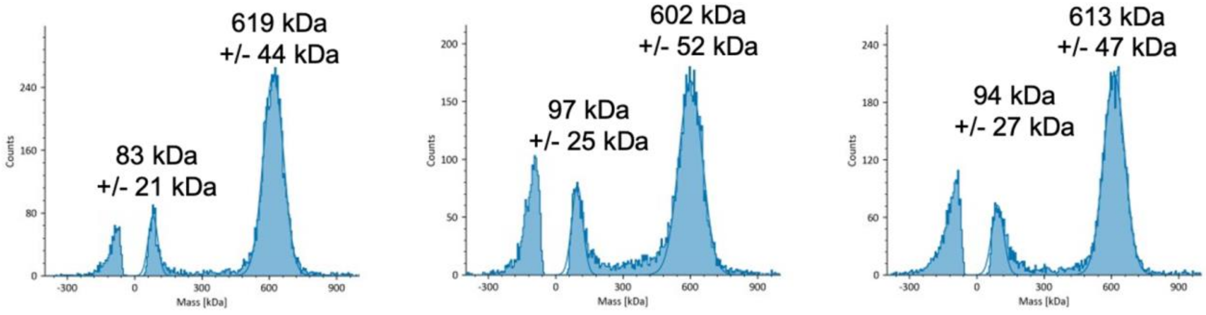

**Figure S2. Mass photometry of Mgm101-DNA complexes**. Three replicates are shown for each different type of complex: 500 nM Mgm101 with a stoichiometric amount of 83+ ssDNA (a), 87+ ssDNA (b), with 83+:83- added sequentially (c), and with pre-formed 83+/83- dsDNA (d). Each peak is labeled with the measured mass and fitting error (sigma) in kDa. From the three replicates, the mean and standard deviation of the mass of each species are reported in the main text Table 1. The theoretical masses of an Mgm101 19-mer with 83+/87+ and 83+:83- (or 83+:83+) are 567 and 598 kDa, respectively. As the Mgm101 monomer (28 kDa) is below the MP detection limit (30 – 40 kDa), the peak at the low mass region could arise from cumulative histogram artifact peaks.

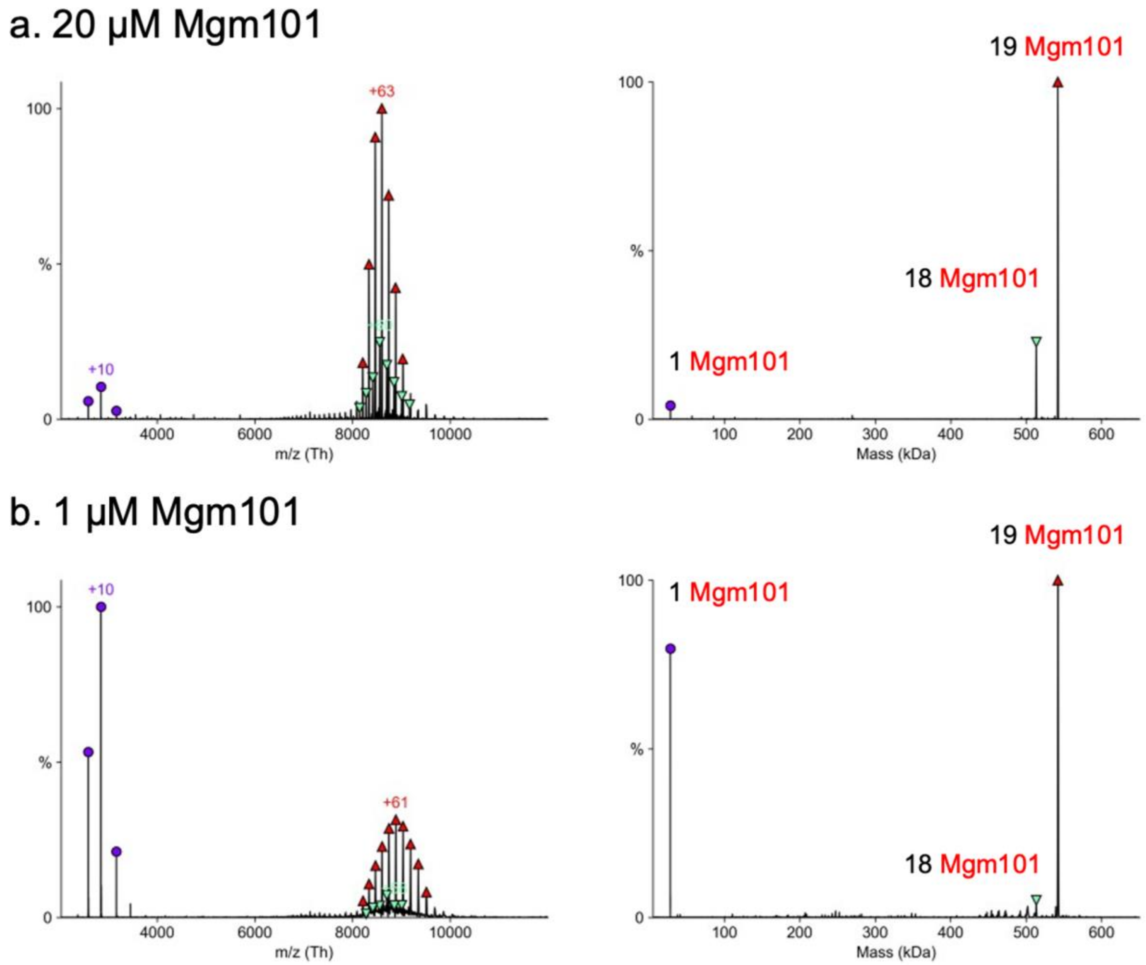


**Figure S3. Native MS of Mgm101 at different concentrations without DNA.** Raw (left) and deconvolved (right) spectra of Mgm101 alone at (**a)** 20 μM and (**b**) 1 μM monomer. Together with main text Fig. 2a (at 10 μM), notice that the ratio of oligomer to monomer increases at higher concentrations, indicating a cooperative assembly. The experimental and theoretical masses of the labeled species are provided in Supplementary Table 3.

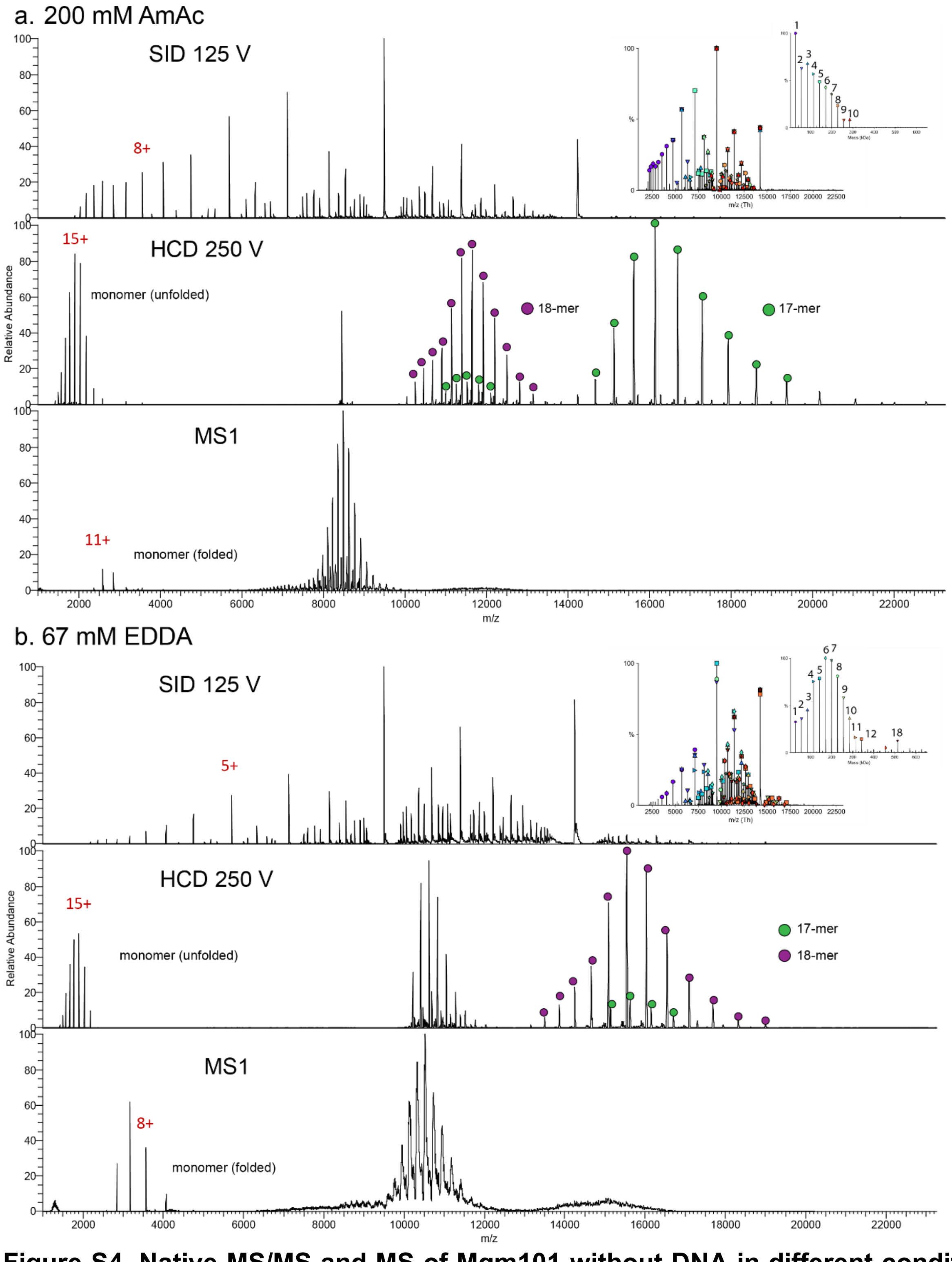


**Figure S4. Native MS/MS and MS of Mgm101 without DNA in different conditions.** Surface induced dissociation (SID) 125 V (top), higher energy collisional induced dissociation (HCD) 250 V (middle), and without any fragmentation, MS1 (bottom) in (**a**) 200 mM ammonium acetate (AmAc) and (**b**) 67 mM Ethylenediamine-N, N'-diacetic acid (EDDA). EDDA is expected to reduce the charge states of species relative to AmAc and was used as a control. In both solvents, HCD generates completely unfolded monomers at a 15+ charge state, 17-mers from 18-mer precursors, and 18-mers from 19-mer precursors. SID generates a variety of fragments from monomer to high-order oligomeric species. The monomer charge state generated from both HCD (15+) and SID (8+ and 5+) differs from the charge state generated without any activation (11+ and 8+), suggesting that the monomer and 18-mer exists in solution and is not a byproduct of the experimental conditions.

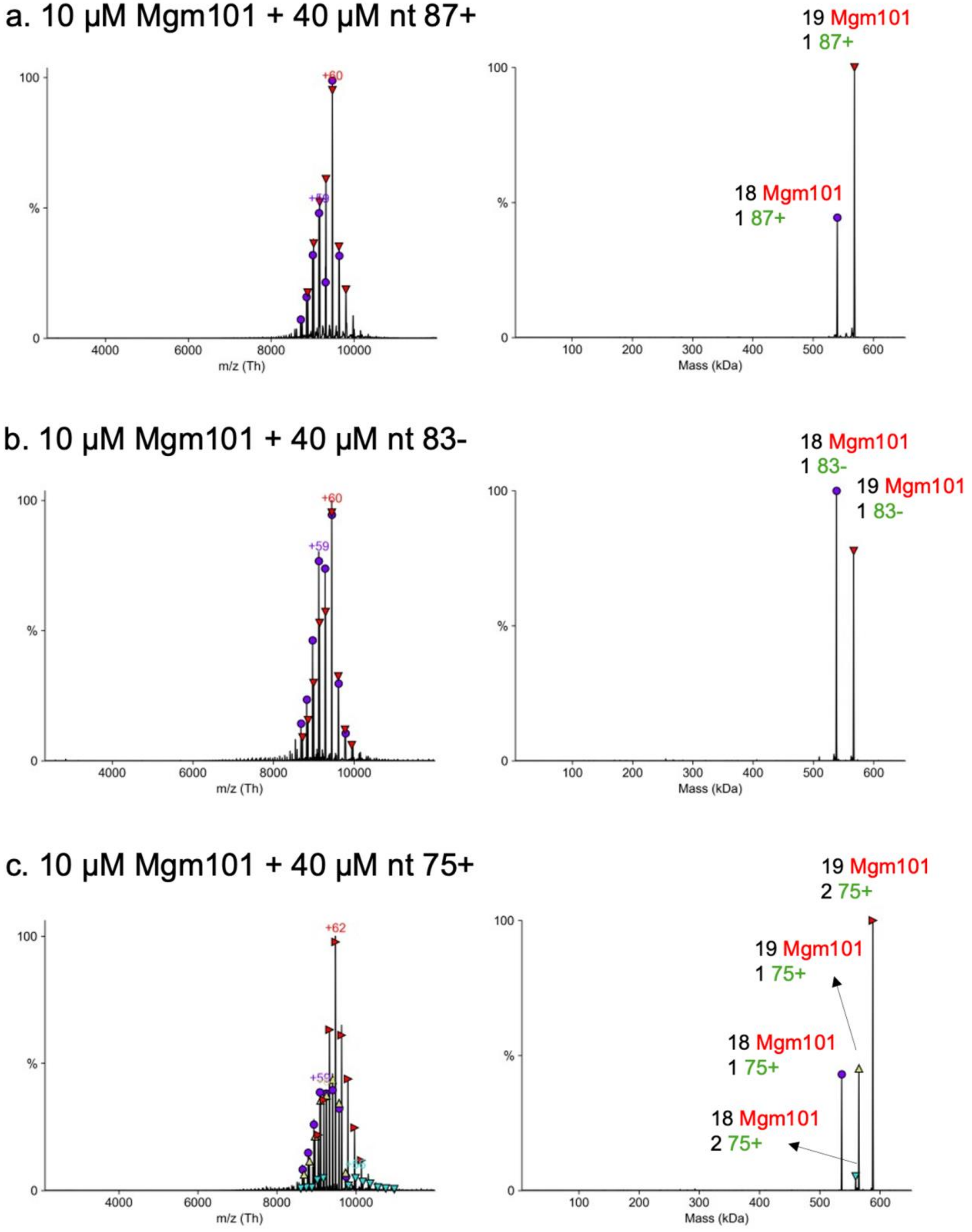
a. 10 µM Mgm101 + 40 µM nt 87+
19 Mgm101
1 87+
18 Mgm101
1 87+
+60
+59
m/z (Th)
Mass (kDa)
b. 10 µM Mgm101 + 40 µM nt 83-
18 Mgm101
1 83-
19 Mgm101
1 83-
+60
+59
m/z (Th)
Mass (kDa)
c. 10 µM Mgm101 + 40 µM nt 75+
19 Mgm101
2 75+
19 Mgm101
1 75+
18 Mgm101
1 75+
18 Mgm101
2 75+
+62
+59
m/z (Th)
Mass (kDa)

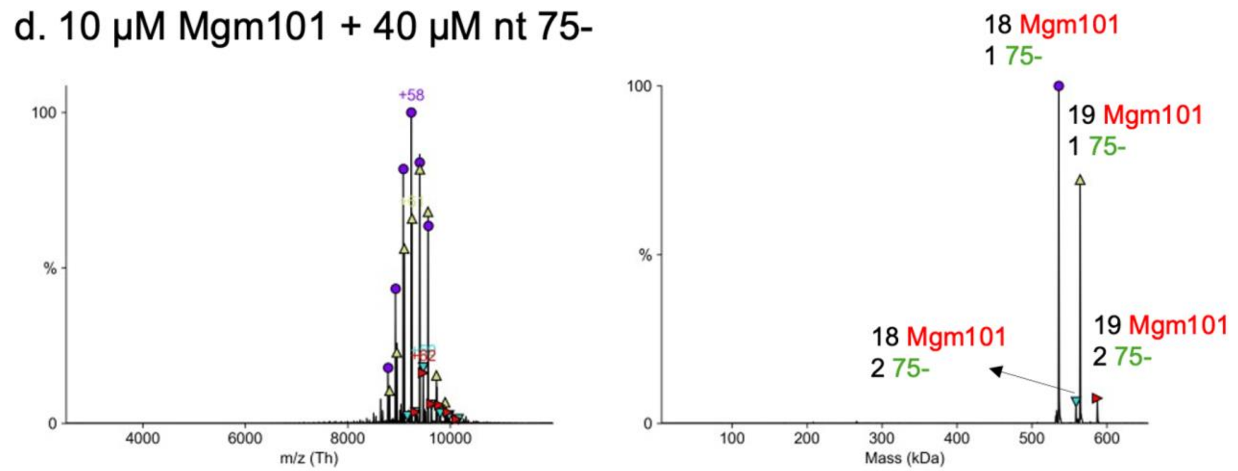


**Figure S5. Native MS of Mgm101 with additional sequences of ssDNA.** Raw (left) and deconvolved (right) spectra of Mgm101 mixed with stoichiometric amounts of (a) 87+, (b) 83-, (c) 75+, and (d) 75-. As also seen in Fig. 2b of the main text for 83+, dominant complexes containing Mgm101 18- or 19-mers bound to one copy of each ssDNA are observed. For 75- (panel d), minor peaks are seen for complexes with two copies of 75-, and for 75+ (panel c) more substantial peaks for two-strand complexes are observed. These two-strand complexes could arise from attempts at annealing at sites of partial complementarity. The experimental and theoretical masses of the labeled species are provided in Supplementary Table 3.

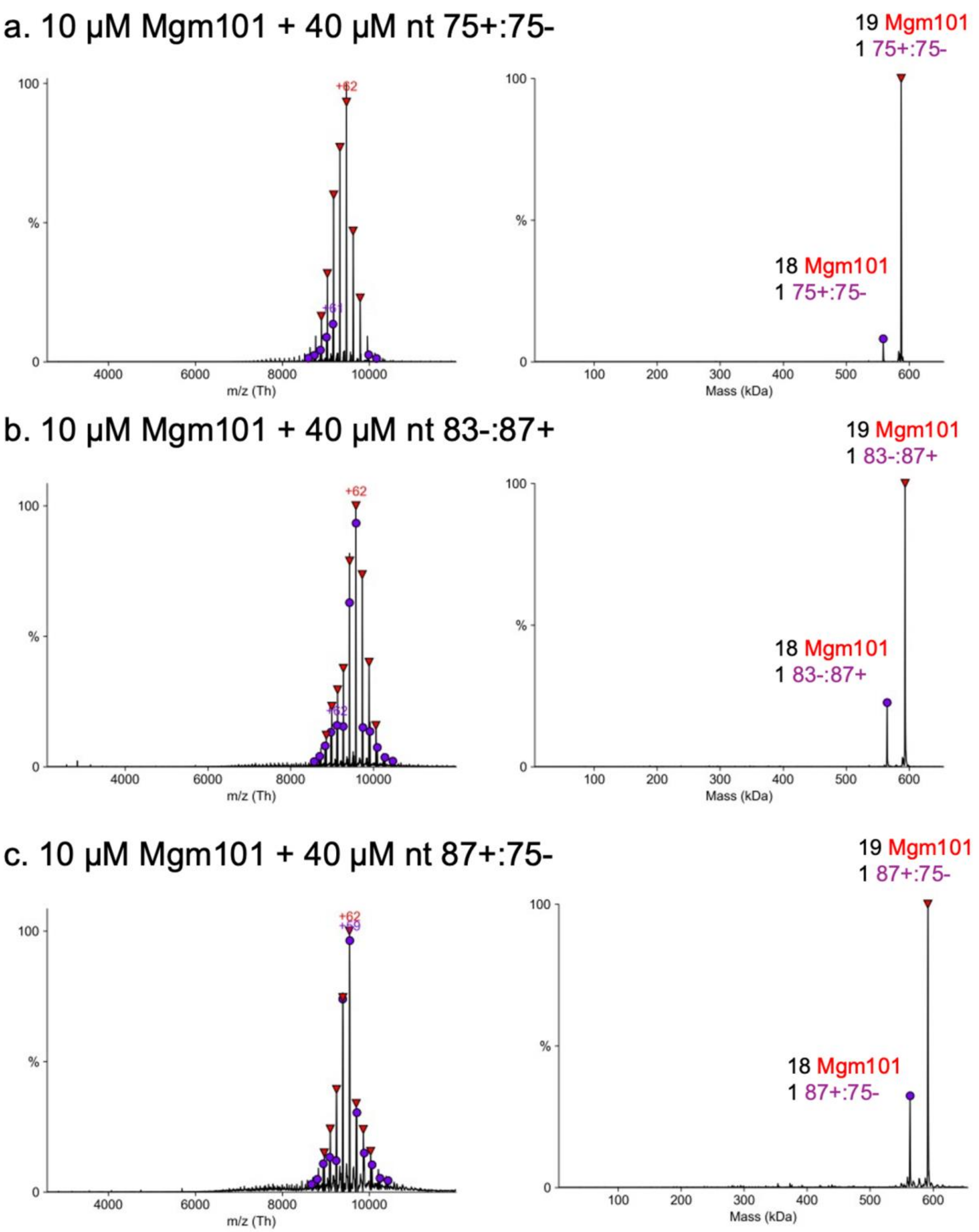

a. 10 µM Mgm101 + 40 µM nt 75+:75-
19 Mgm101
1 75+:75-
18 Mgm101
1 75+:75-
+62
+61
100
%
0
4000
6000
8000
10000
m/z (Th)
100
200
300
400
500
600
Mass (kDa)
b. 10 µM Mgm101 + 40 µM nt 83-:87+
19 Mgm101
1 83-:87+
18 Mgm101
1 83-:87+
+62
+62
m/z (Th)
Mass (kDa)
c. 10 µM Mgm101 + 40 µM nt 87+:75-
19 Mgm101
1 87+:75-
18 Mgm101
1 87+:75-
+62
+59
m/z (Th)
Mass (kDa)

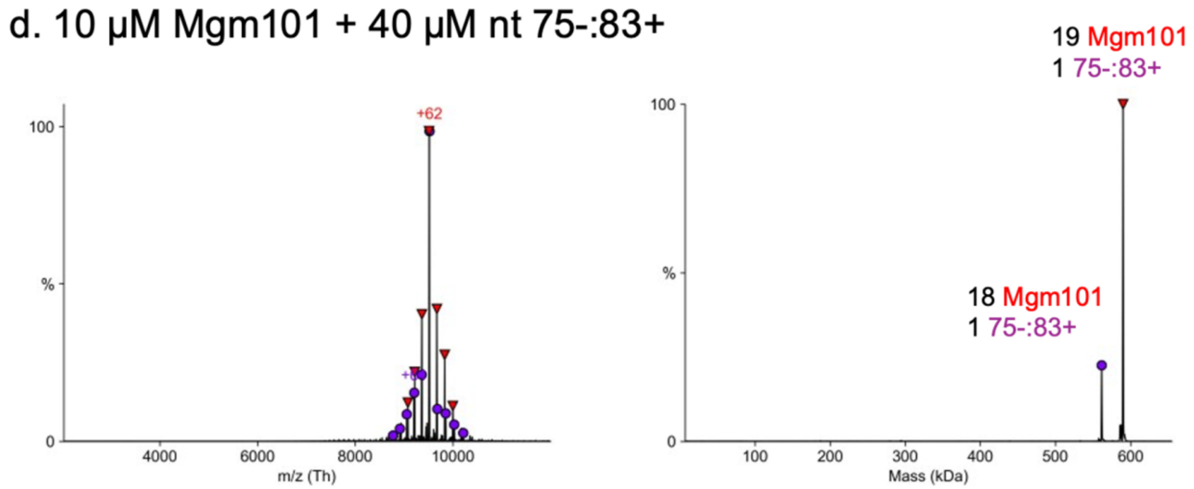


**Figure S6. Native MS of Mgm101 mixed with sequentially added complementary ssDNA to form a complex with a duplex intermediate of annealing.** Raw (left) and deconvolved (right) spectra of Mgm101 mixed with (a) 75+:75-, (b) 83-:87+, (c) 87+:75- and (d) 75-:83+. To form the complex, the first ssDNA (first: second) was added to the protein at 4 nt per monomer and incubated for 15 min at 37 °C, and then the second ssDNA was added and incubated for an additional 15 min at 37 °C. In all cases, dominant complexes containing one Mgm101 18-mer or 19-mer and one copy of each strand (i.e. two complementary strands) are observed. The experimental and theoretical masses of the labeled species are provided in Supplementary Table 3.

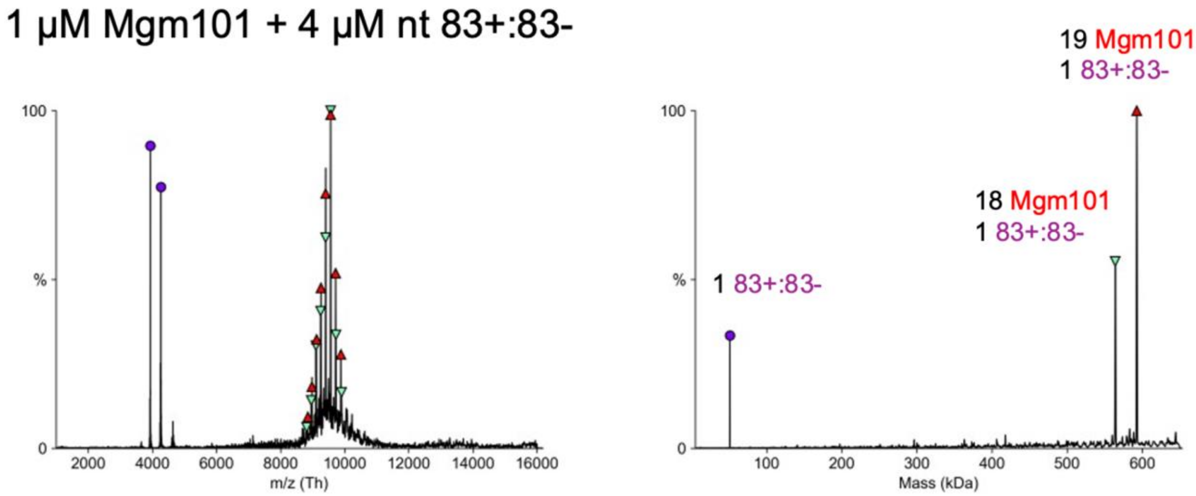


**Figure S7. Native MS of Mgm101 mixed with sequentially added 83+:83- at 10-fold lower (1 μM) concentration.** Raw (left) and deconvolved (right) spectra of Mgm101 mixed with 83+:83-. As seen at higher (10 µM) concentrations (main text Fig. 2c), the complex with two complementary strands again emerged as the dominant species. Although a small peak for unbound duplex was also observed, no peaks were observed for unbound Mgm101 oligomer or monomer, or Mgm101 monomer bound to DNA. The experimental and theoretical masses of the labeled species are provided in Supplementary Table 3.

a. 10 µM Mgm101 + 40 µM 87NC:83-

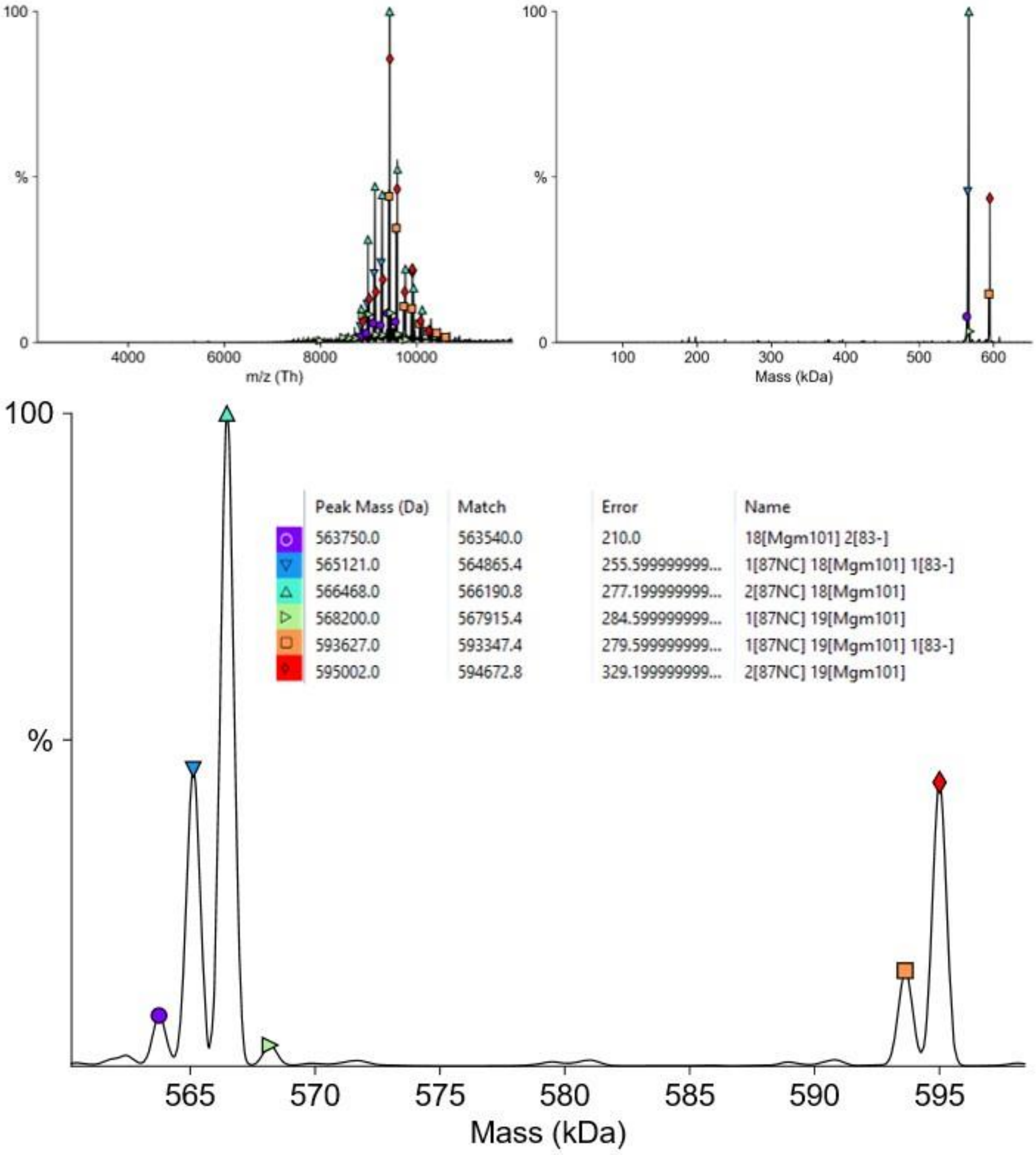

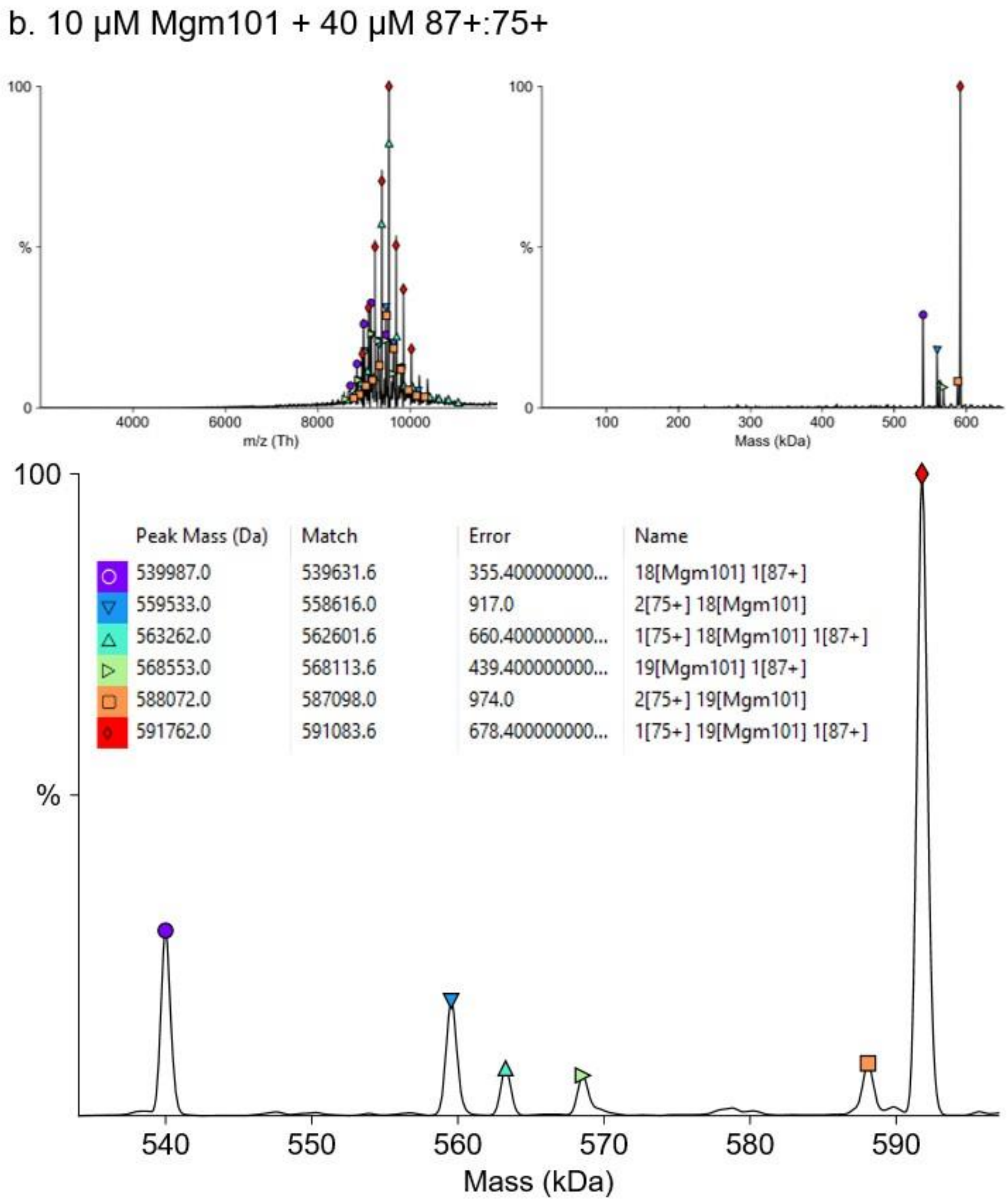


**Figure S8. Native MS of Mgm101 mixed with two non-complementary ssDNAs.** Raw (left) and deconvolved (right) spectra are shown for Mgm101 mixed with (**a**) 87NC:83-, and (**b**) 87+:75+. The lower panels show a blow up of the region of the deconvoluted spectrum containing the multiple oligomeric species of interest (far right of the deconvoluted spectrum), which are defined by the colored symbols and detailed in the legend with observed mass measurement (Peak Mass) and difference (Error) from theoretical value (Match) for each specified complex. Notice that in both panels, a mixture of multiple species containing complexes of Mgm101 18- or 19-mers bound to different combinations of one or two ssDNA strands are observed, as opposed to the dominant complexes observed when two complementary strands are added to Mgm101 sequentially. The experimental and theoretical masses of the labeled species are provided in Supplementary Table 3 (and in the lower panels).

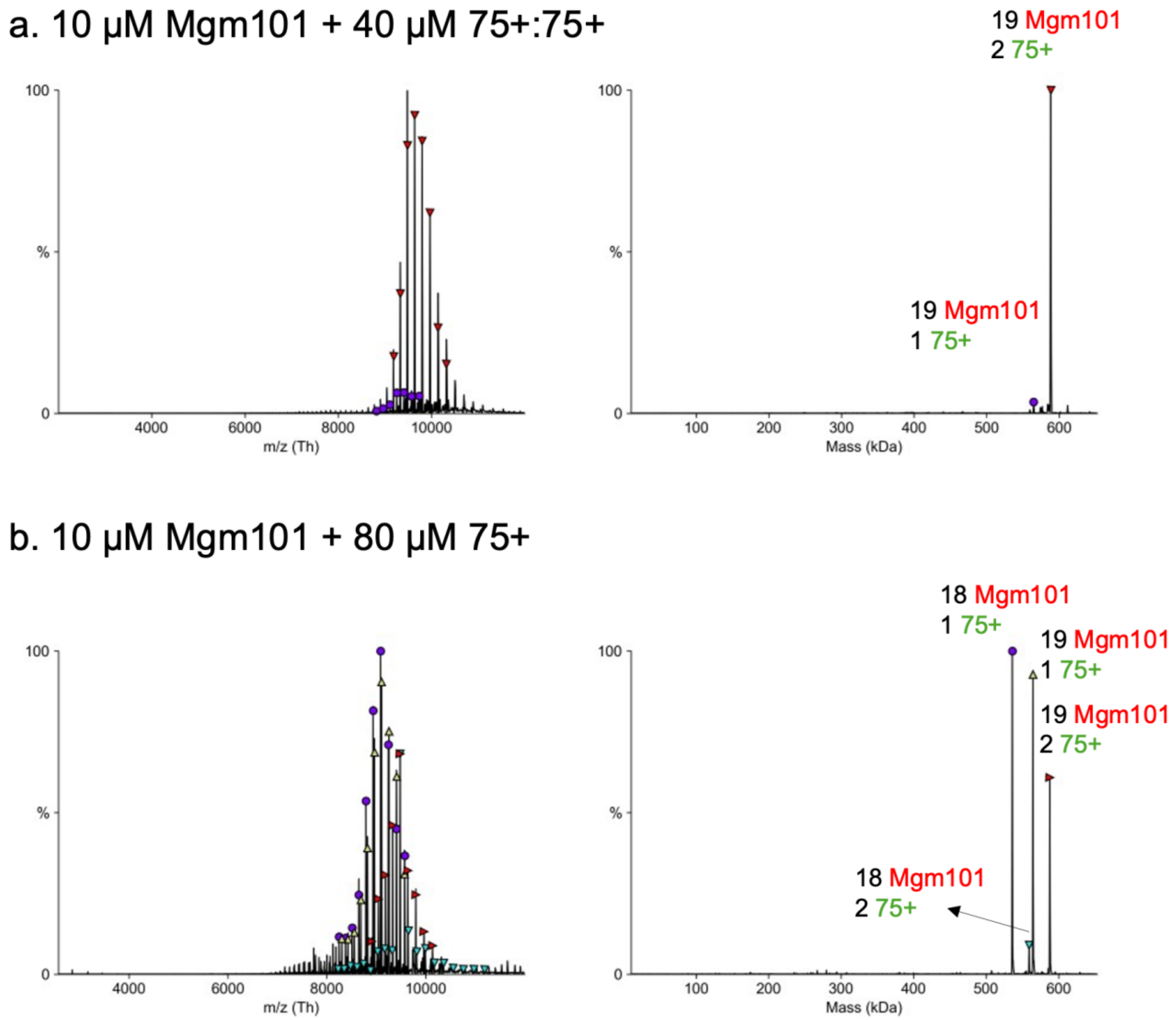


**Figure S9. Native MS of Mgm101 mixed with a 2-fold excess of 75+ ssDNA.** Raw (left) and deconvolved (right) spectra of Mgm101 mixed with (**a**) 75+:75+ (a second dose of the same 75+ is added after a first addition), and (**b**) 75+ added at 2x concentration for the first addition. The complexes of Mgm101 bound to two copies of the same ssDNA likely represent attempts at annealing at sites of partial complementarity. The experimental and theoretical masses of the labeled species are provided in Supplementary Table 3.

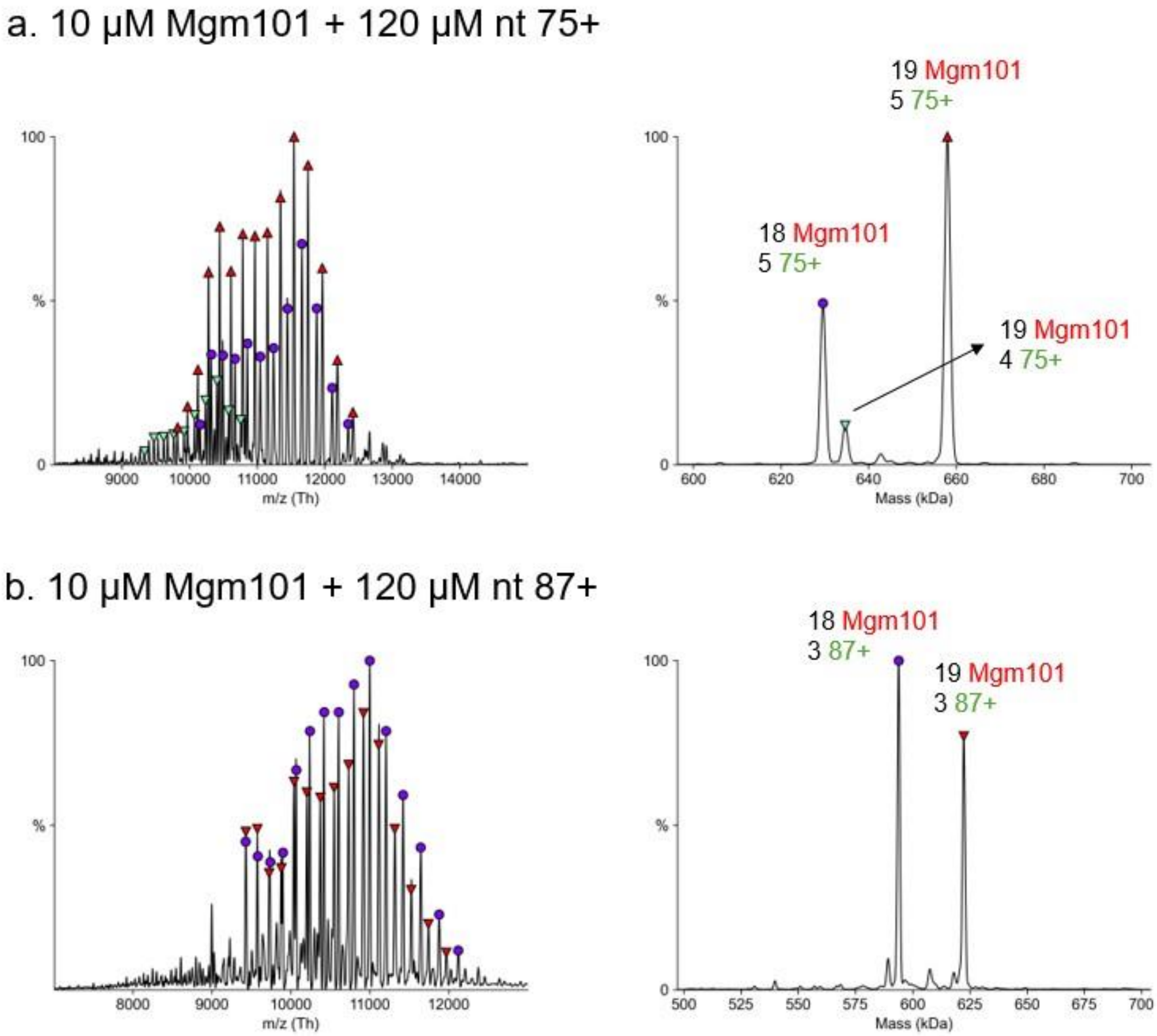


**Figure S10. Native MS of Mgm101 mixed with a 3-fold excess of ssDNA.** Raw (left) and deconvolved (right) spectra of Mgm101 mixed with 3x concentration of (**a**) 75+, and (**b**) 87+. The complexes with >2 copies of ssDNA could conceivably arise from two strands binding in the main (inner) DNA-binding groove, and additional strands binding non-specifically to the C-lobes. Alternatively, strands of ssDNA could only partially occupy the main DNA-binding groove (i.e. not fully encircle), such that >2 strands can bind. The experimental and theoretical masses of the labeled species are provided in Supplementary Table 3.

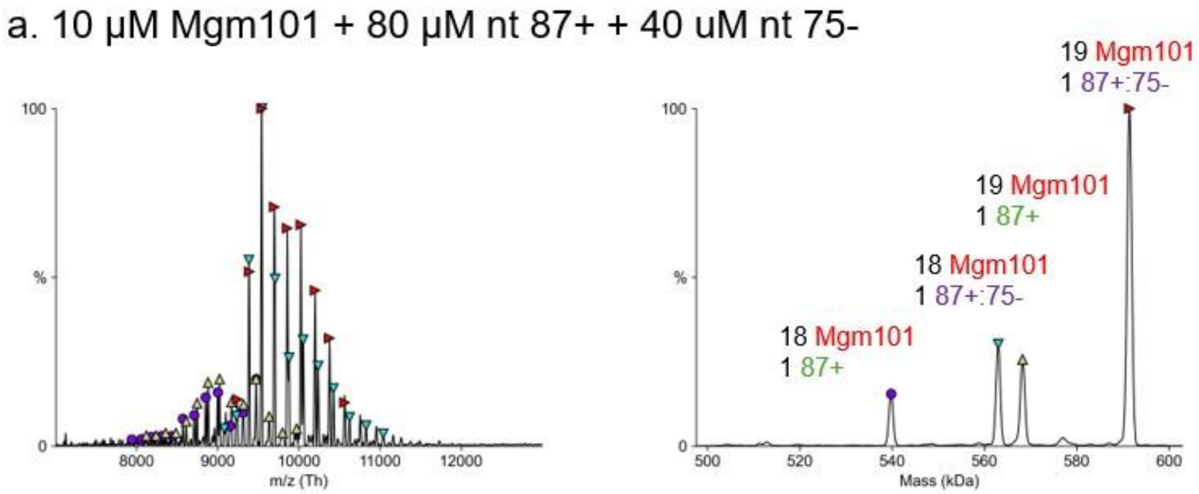


**Figure S11. Native MS of Mgm101 mixed with an excess of one ssDNA followed by an equivalent of the complementary ssDNA.** Raw (left) and deconvolved (right) spectra of Mgm101 mixed with the first strand (87+) in 2x molar excess, and the complementary strand (75-) in a molar equivalent. Notice that while multiple species are observed, the complexes with one copy of each strand (i.e. two complementary strands) emerge as dominant. This shows that the complementary strand can replace the non-complementary strand. The experimental and theoretical masses of the labeled species are provided in Supplementary Table 3.

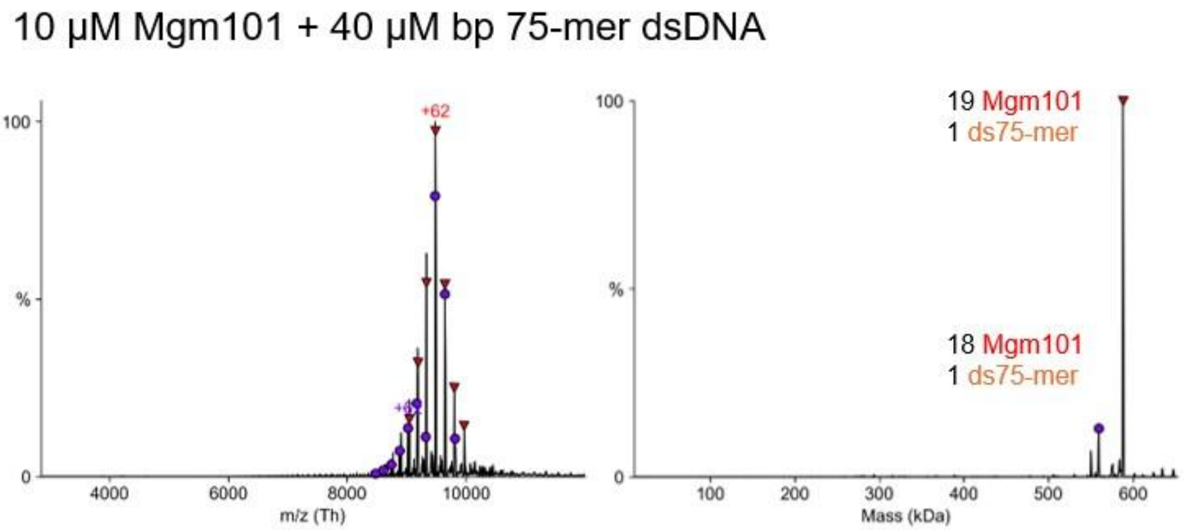


**Figure S12. Native MS of Mgm101 mixed with pre-formed dsDNA.** Raw (left) and deconvolved (right) spectra of Mgm101 mixed with 75-mer pre-formed dsDNA. A dominant complex with one copy of the dsDNA is observed, as was the case for the 83-mer dsDNA shown in main text Fig. 2d. The experimental and theoretical masses of the labeled species are provided in Supplementary Table 3.

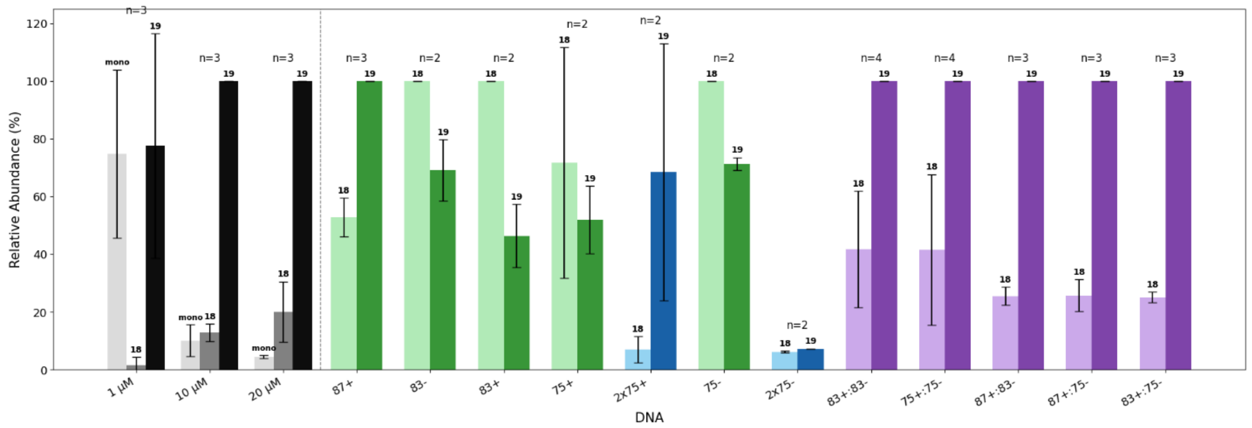


**Figure S13. Relative amounts of the Mgm101 18 and 19-mer species observed in nMS**. Histograms in grey and black are species observed without DNA, where the concentrations are listed below. Histograms in green are species produced when only one equivalent of a ssDNA strand is added. Histograms in blue represent species observed when two molar equivalents of one ssDNA were added. Histograms in purple represent species observed when two different strands were added sequentially.

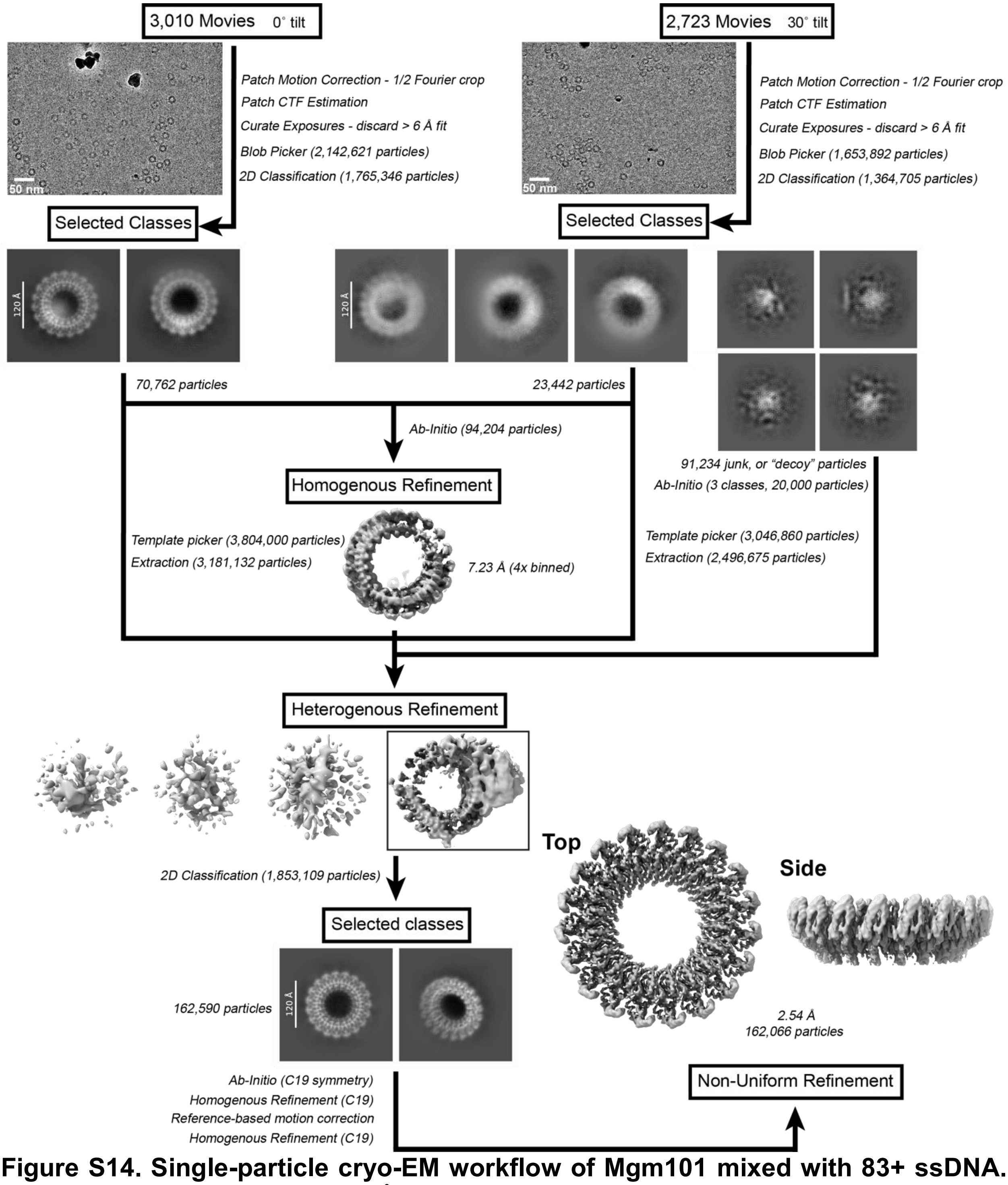


**Figure S14. Single-particle cryo-EM workflow of Mgm101 mixed with 83+ ssDNA.** The workflow resulted in a 2.54 Å stucture of Mgm101 bound to 83+ ssDNA (PDB ID: 9YI6, EMD-72979, EMPIAR-13023).

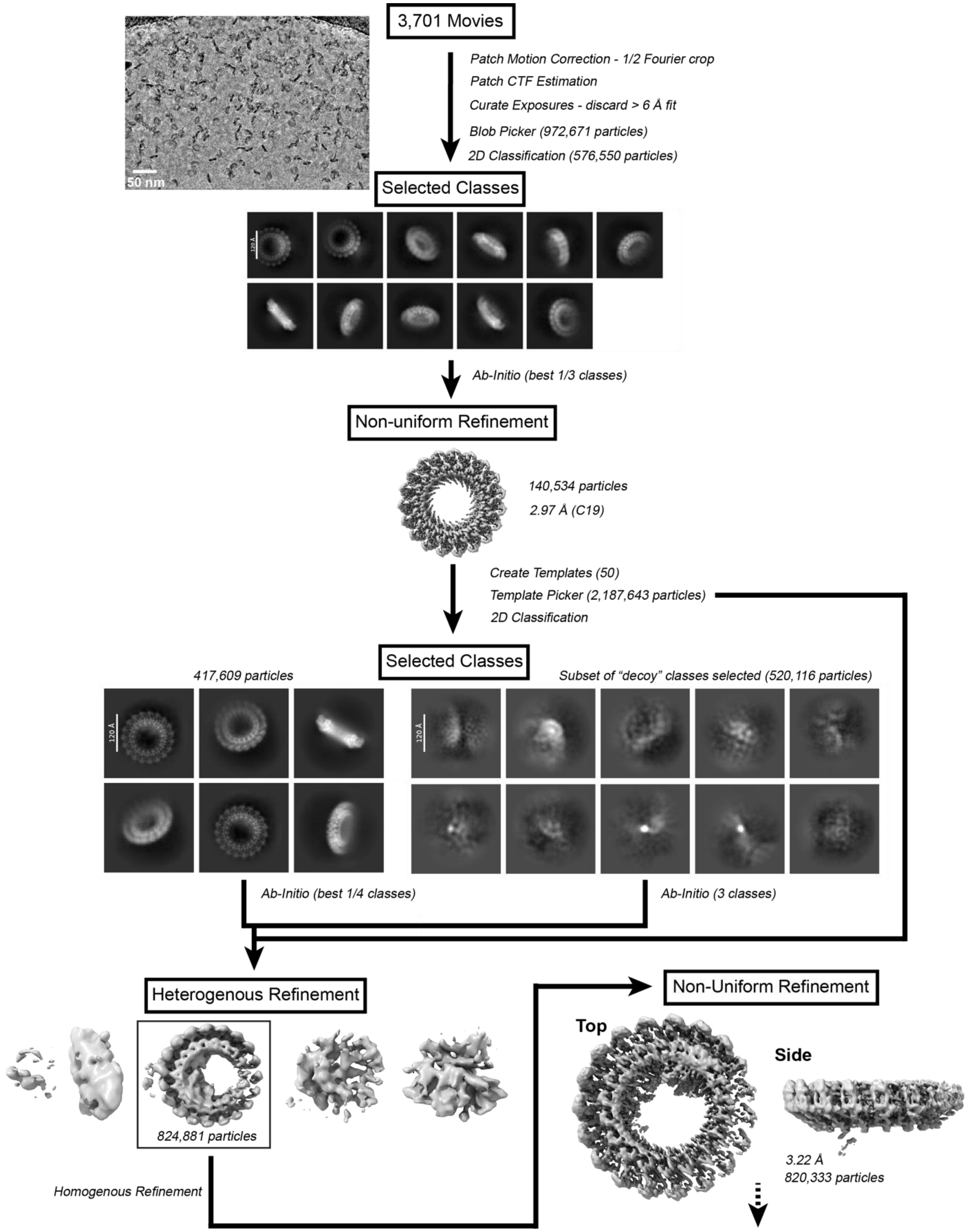

50 nm
3,701 Movies
Patch Motion Correction - 1/2 Fourier crop
Patch CTF Estimation
Curate Exposures - discard > 6 Å fit
Blob Picker (972,671 particles)
2D Classification (576,550 particles)
Selected Classes
120 Å
Ab-Initio (best 1/3 classes)
Non-uniform Refinement
140,534 particles
2.97 Å (C19)
Create Templates (50)
Template Picker (2,187,643 particles)
2D Classification
Selected Classes
417,609 particles
Subset of “decoy” classes selected (520,116 particles)
120 Å
120 Å
Ab-Initio (best 1/4 classes)
Ab-Initio (3 classes)
Heterogenous Refinement
824,881 particles
Homogenous Refinement
Non-Uniform Refinement
Top
Side
3.22 Å
820,333 particles

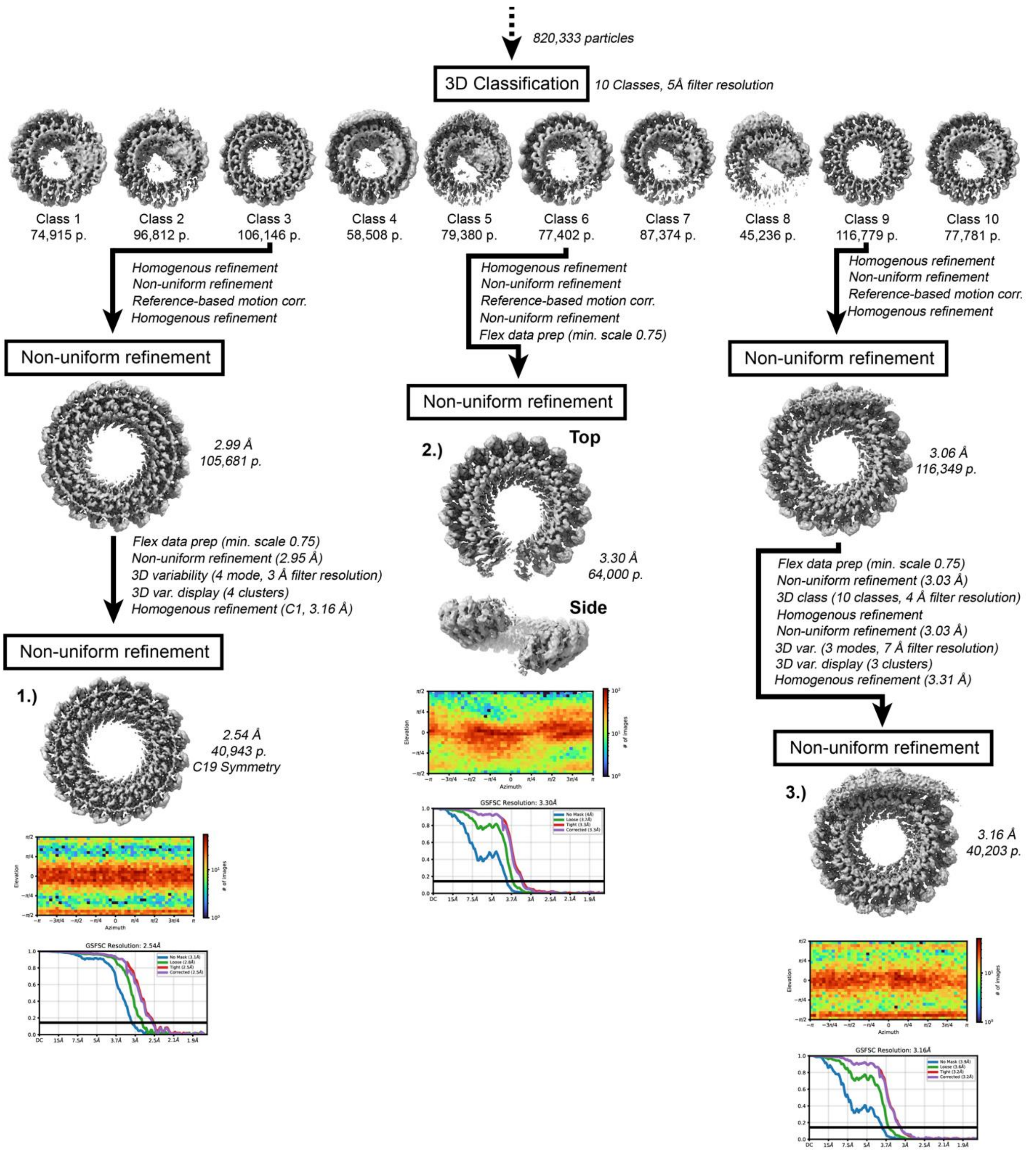


**Figure S15. Single-particle cryo-EM workflow of Mgm101 mixed with two complementary ssDNAs (75+ and 75-) added sequentially.** The workflow resulted in (1) Mgm101 19-mer bound to 75+:75- duplex annealing intermediate (left, PDB ID: 9YI7, EMD-72980, EMPIAR-13025), (2) apo Mgm101 18-mer lock washer (middle), and (3) Mgm101 19-mer bound to B-form dsDNA (right, PDB ID: 9YI8, EMD-72981, EMPIAR-13025).

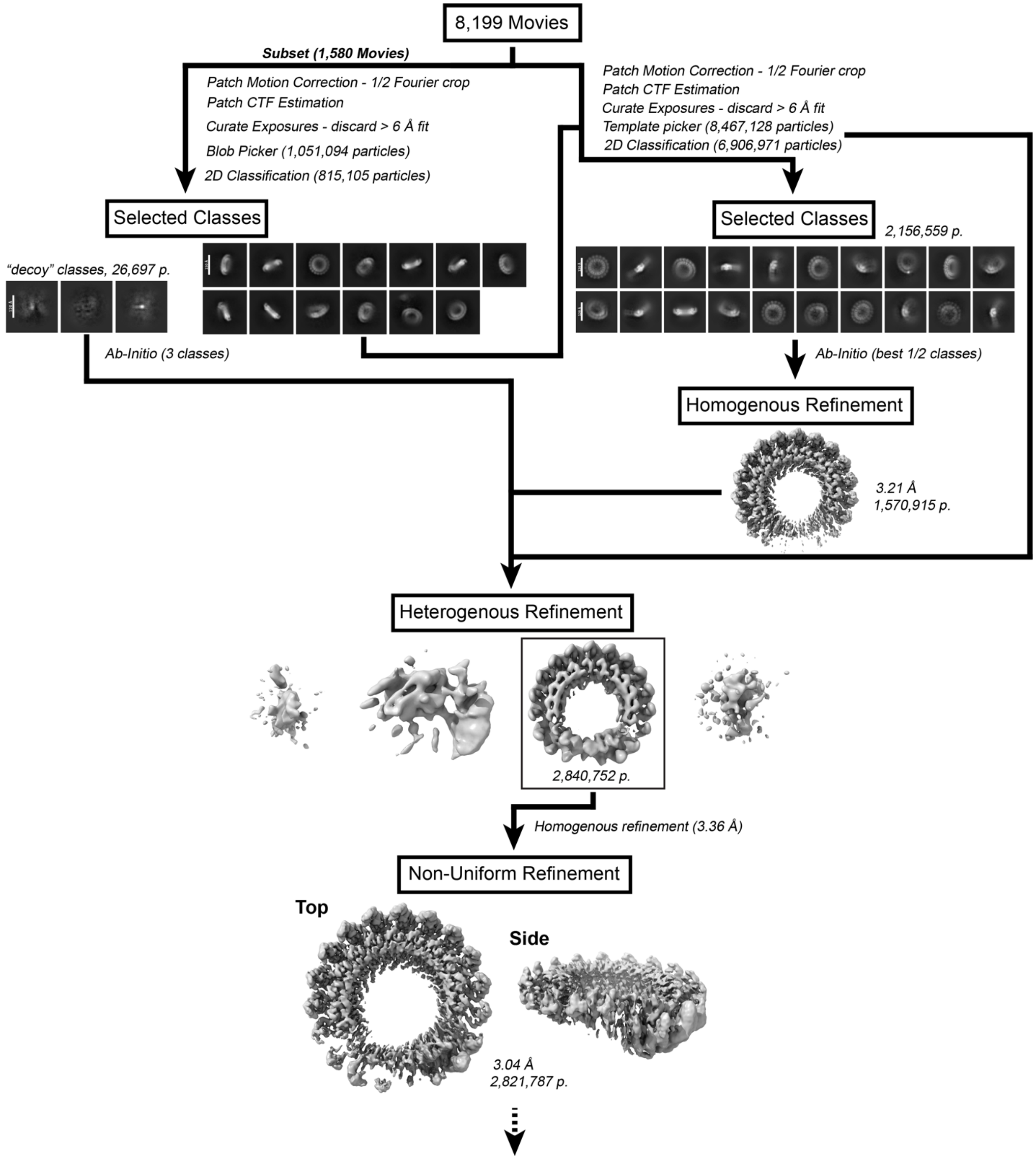
8,199 Movies
Subset (1,580 Movies)
Patch Motion Correction - 1/2 Fourier crop
Patch CTF Estimation
Curate Exposures - discard > 6 Å fit
Blob Picker (1,051,094 particles)
2D Classification (815,105 particles)
Patch Motion Correction - 1/2 Fourier crop
Patch CTF Estimation
Curate Exposures - discard > 6 Å fit
Template picker (8,467,128 particles)
2D Classification (6,906,971 particles)
Selected Classes
"decoy" classes, 26,697 p.
Ab-Initio (3 classes)
Selected Classes
2,156,559 p.
Ab-Initio (best 1/2 classes)
Homogenous Refinement
3.21 Å
1,570,915 p.
Heterogenous Refinement
2,840,752 p.
Homogenous refinement (3.36 Å)
Non-Uniform Refinement
Top
Side
3.04 Å
2,821,787 p.

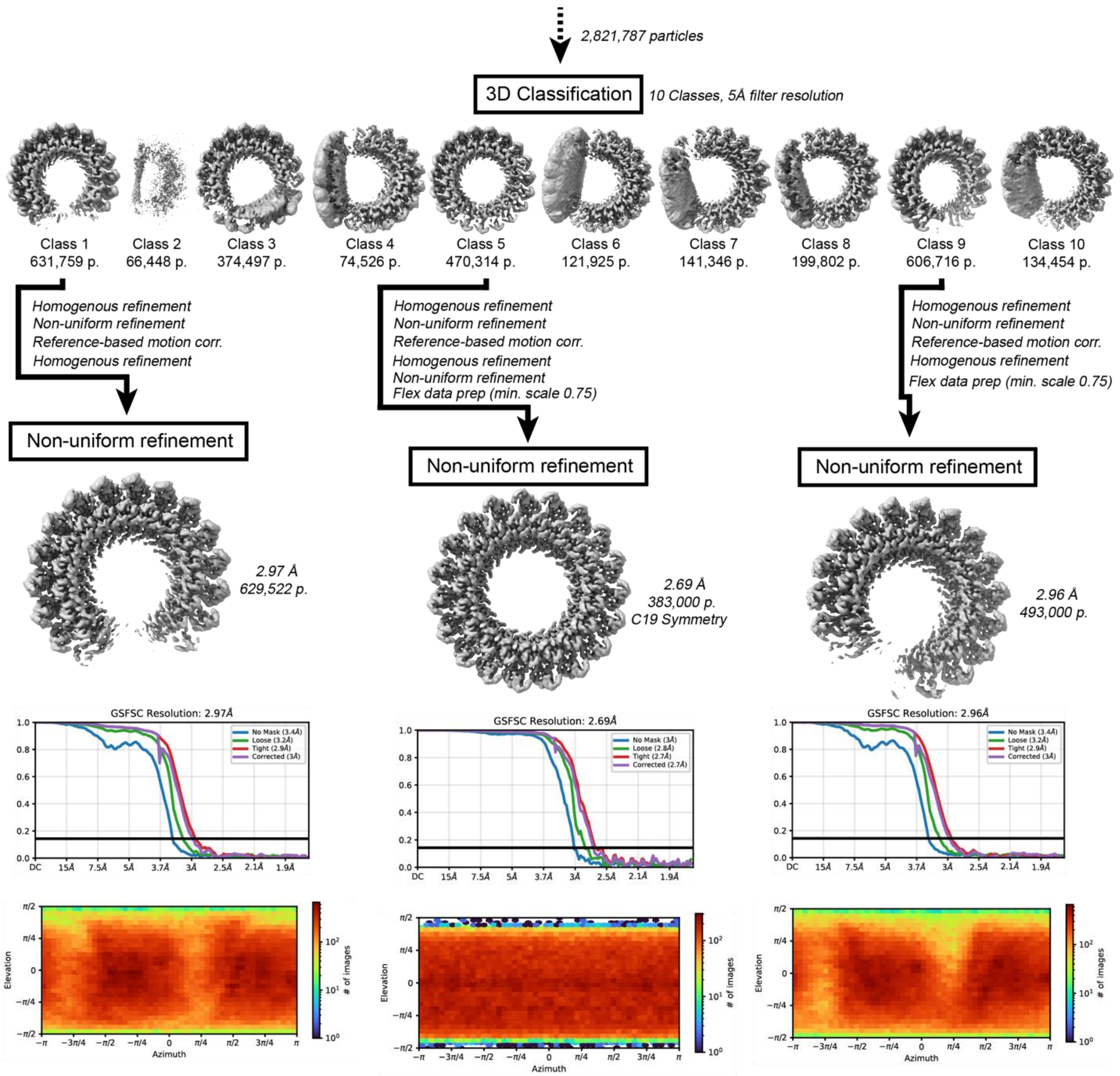


**Figure S16. Single-particle cryo-EM workflow of Mgm101 mixed with pre-formed 83-mer dsDNA.** The workflow resulted in two structures of an apo Mgm101 18-subunit open lock washer (left and right, PDB ID: 9YI9, EMD-72983, EMPIAR-13024), and one structure of apo Mgm101 closed ring 19-mer (middle, PDB ID: 9YIA, EMD-72984, EMPIAR-13024).

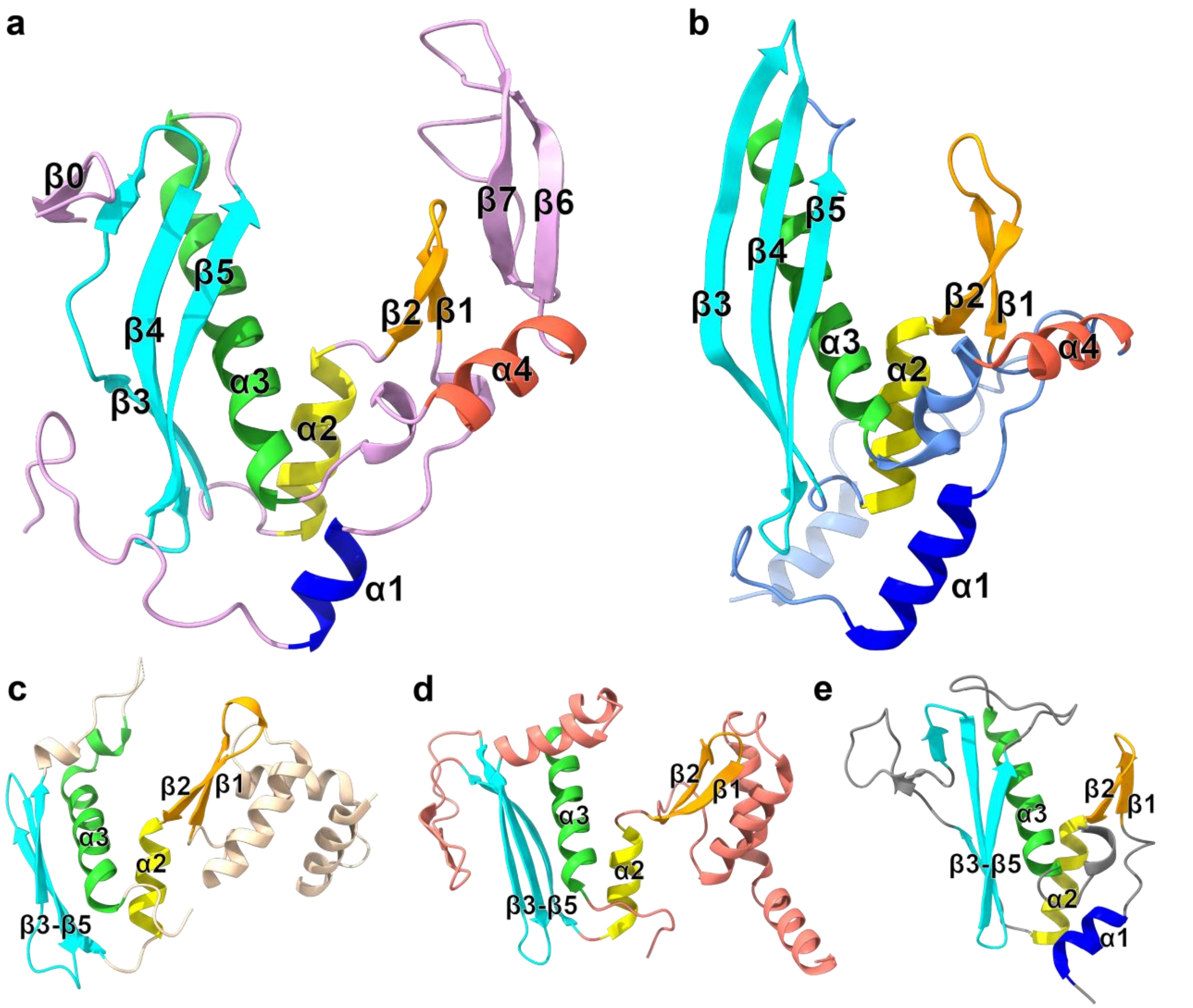


**Figure S17**. **Mgm101 monomer structure comparison to RAD52 and related SSAPs.** **(a)** Mgm101 (PDB ID 9YIA) **(b)** RAD52 (PDB ID 5XRZ) **(c)** Red$\beta$ (PDB ID 7UJL) **(d)** LiRecT (PDB ID 7UB2) **(e)** sak80$\alpha$ (PDB ID 8PQ8). Notice that $\alpha$2-$\alpha$3 in the center (yellow and green), and the $\beta$3-$\beta$5 sheet (cyan) and $\beta$1-$\beta$2 hairpin (orange) on either side of it, are conserved for the family to form a central DNA-binding groove.

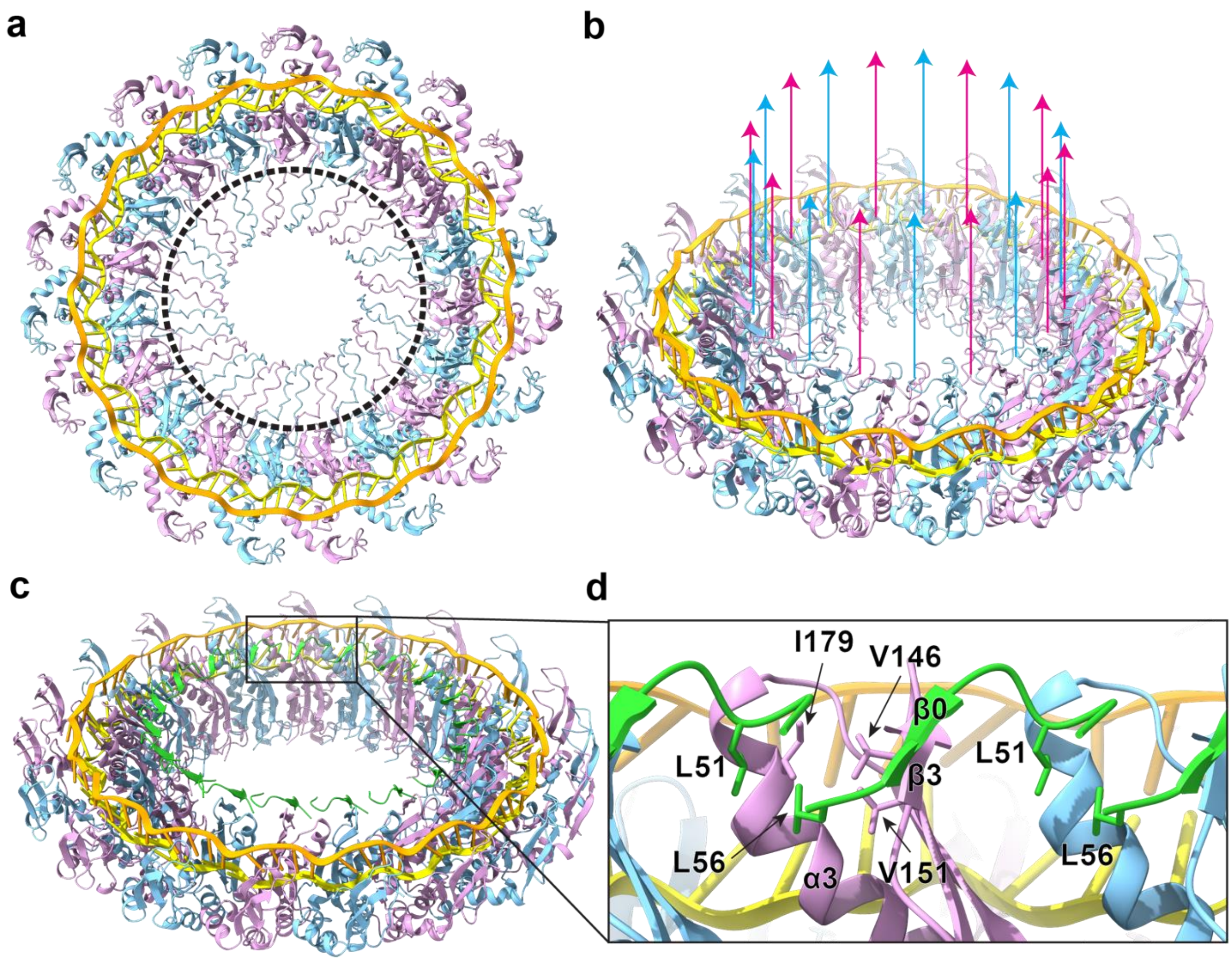


**Figure S18. Plausible locations of missing residues not observed in the Mgm101 structures and the packing of β0 onto β3 at the inner rim of the Mgm101 ring. (a)** Modeling of residues 57-75 (within dotted circle) residing within the Mgm101 ring interior. The disordered segments were taken from an AF3 prediction of monomeric Mgm101 and grafted onto the structure between residues 56-76 of each subunit. (**b**) An oblique view of (a) showing that the N-terminal tail residues 1-48 of each subunit, shown as upwards arrows, are likely to project out from the upper surface of the ring, based on the location of S49, the first ordered residue of the β0 segment. **(c)** An oblique view of the structure of the Mgm101 ring with duplex intermediate, with alternating subunits colored in plum and cyan. N-terminal residues 49-56 containing β0 (residues 53-55) are colored lime green. **(d)** A close-up view of the region boxed in (a), showing the hydrophobic interactions that L51 and L56 make at the subunit interface.

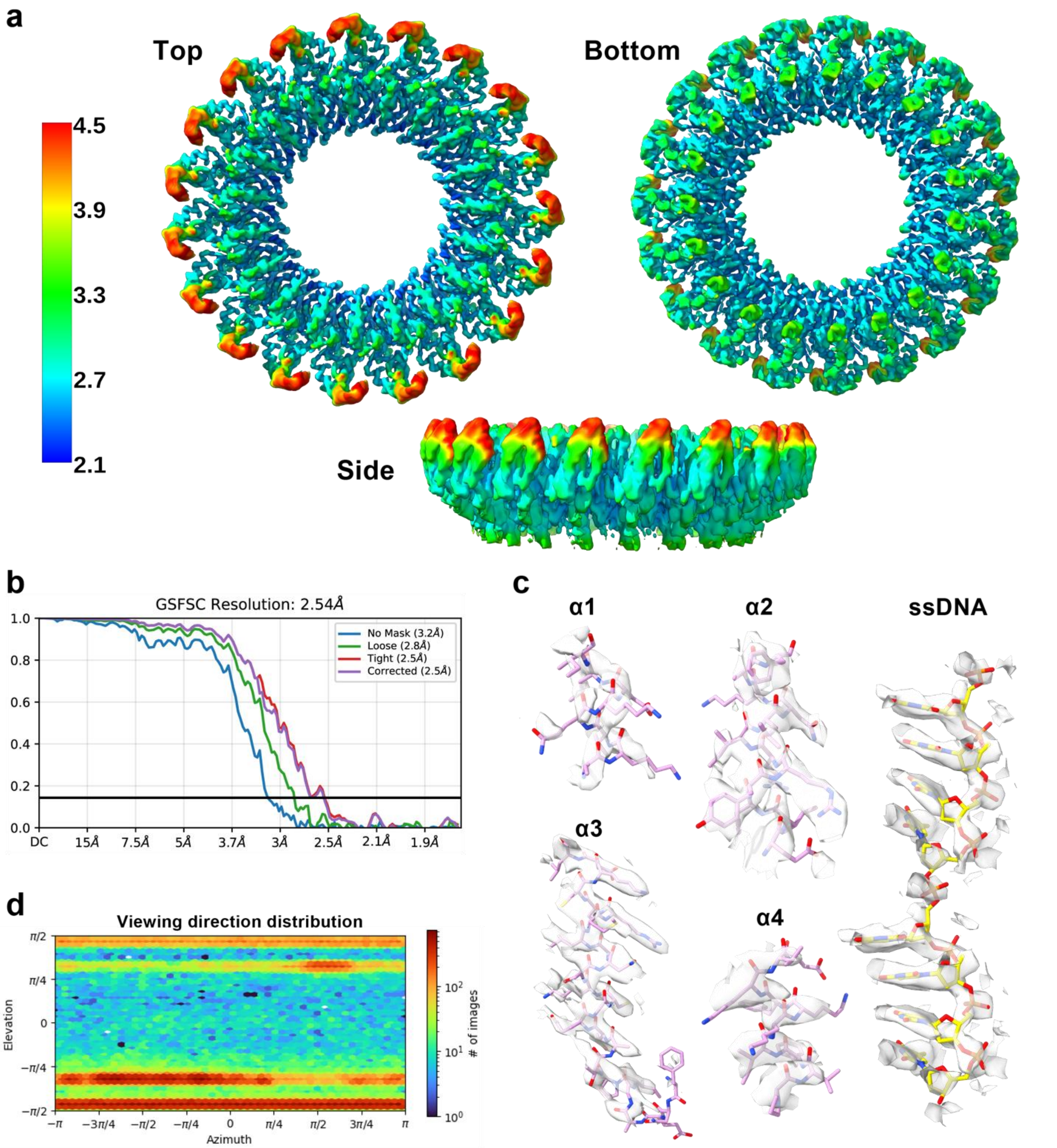


**Figure S19. Cryo-EM structure of Mgm101 complex with 83+ ssDNA.** (**a**) Local resolution estimation of the reconstruction (PDB ID: 9YI6, EMD-72979, EMPIAR-13023). (**b**) GSFSC determination. (**c**) The Mgm101 α-helices and their fit within the density. (**d**) Particle viewing direction distribution.

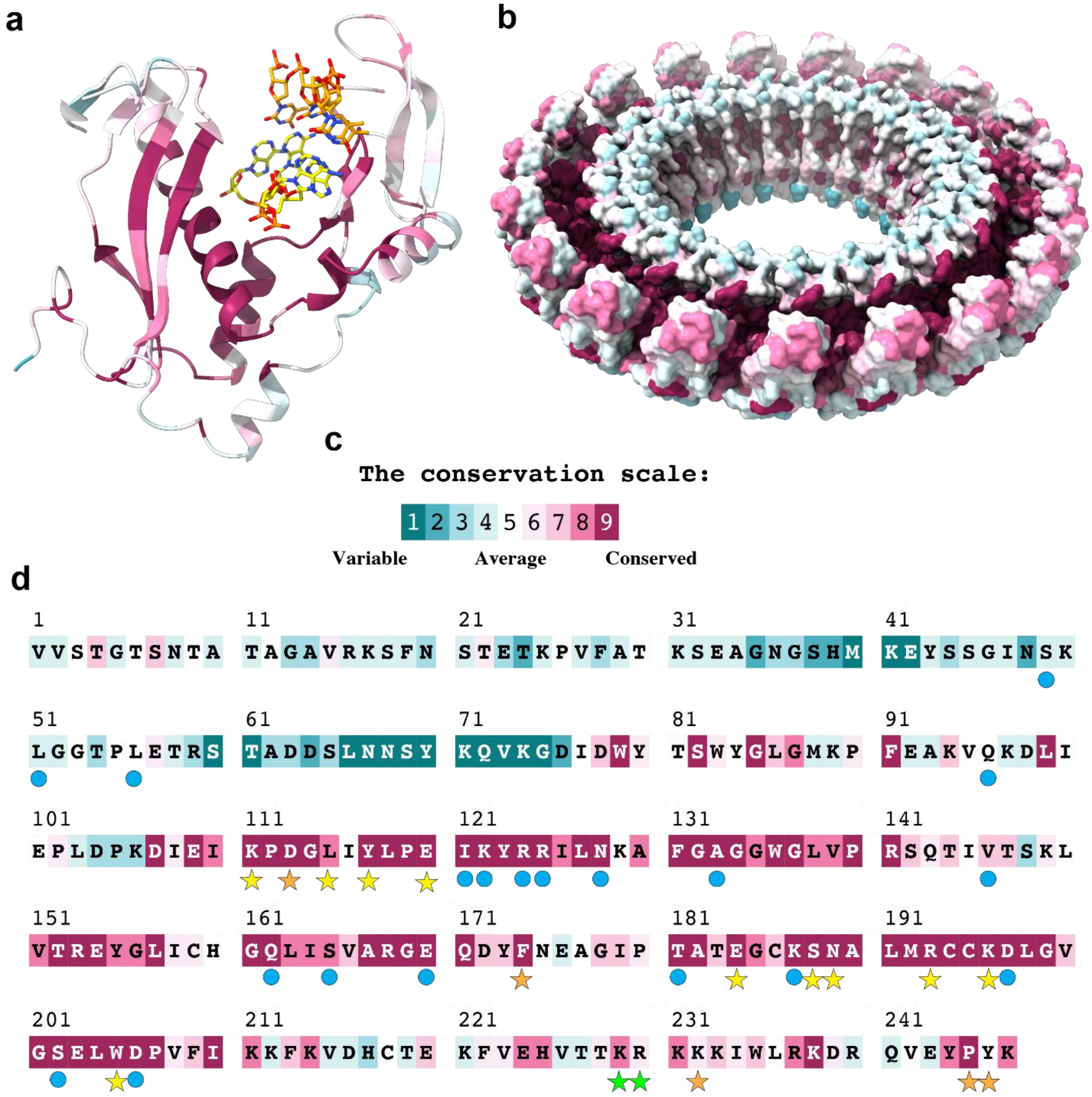


**Figure S20. Structure and sequence of Mgm101 colored by conservation.** (**a**) Monomeric Mgm101, (**b**) 19-mer surface representation, (**c**) the conservation scale, (**d**) and the mature sequence, colored by conservation according to (c), where symbols below correspond to residues for: inner site DNA binding (yellow star), outer site DNA binding (orange star), B-form DNA binding (green star) or inter-subunit interactions (cyan circle). The analysis was performed with ConSurf[7] using default settings.

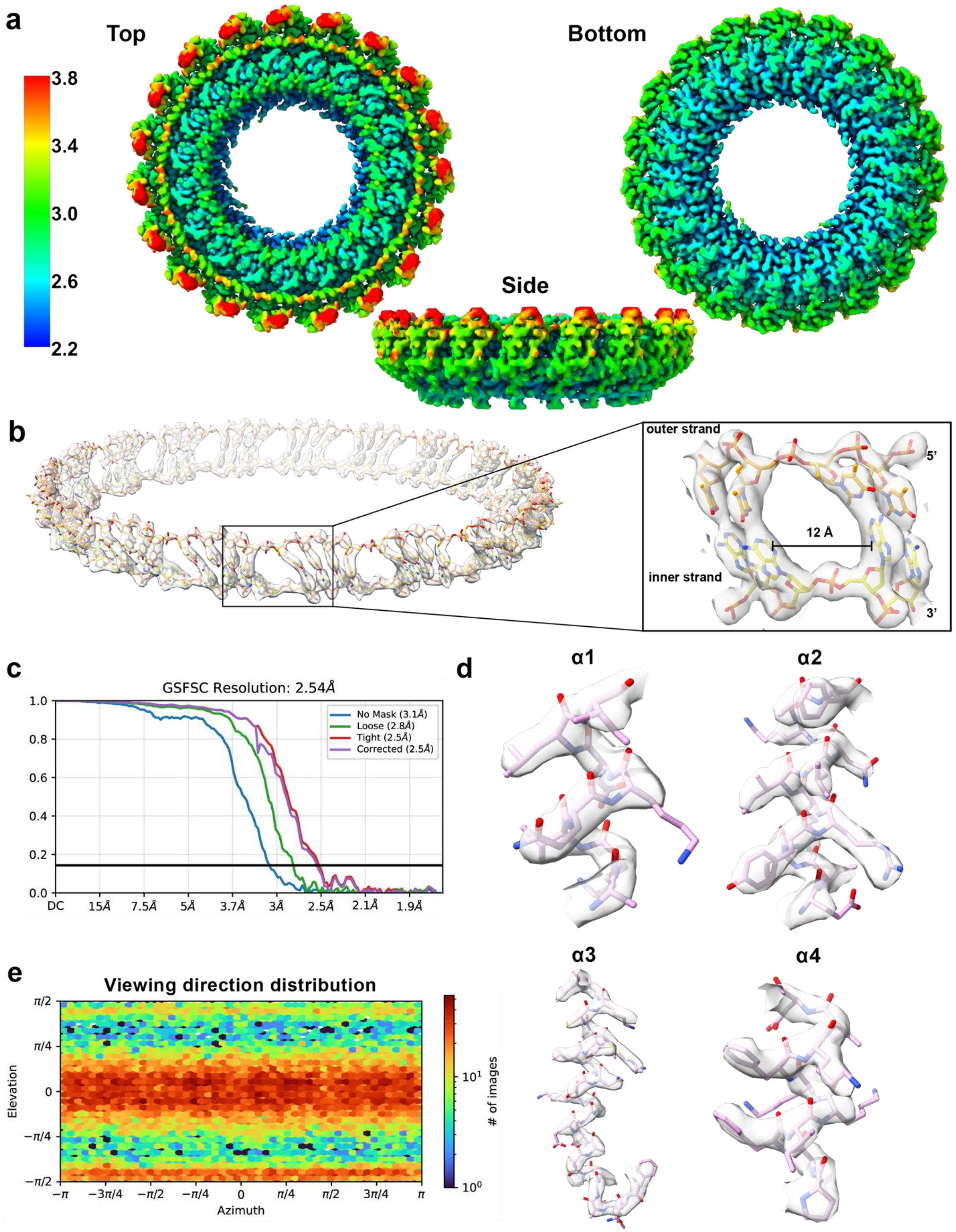


**Figure S21. Cryo-EM structure of Mgm101 complex with 75+ and 75- added sequentially.** (**a**) Local resolution estimation of the reconstruction (PDB ID: 9YI7, EMD-72980, EMPIAR-13025). **(b)** Density corresponding to the DNA duplex annealing intermediate. **(c)** GSFSC resolution. **(d)** Mgm101 α-helices and their fit within the density. **(e)** Particle orientation distribution.

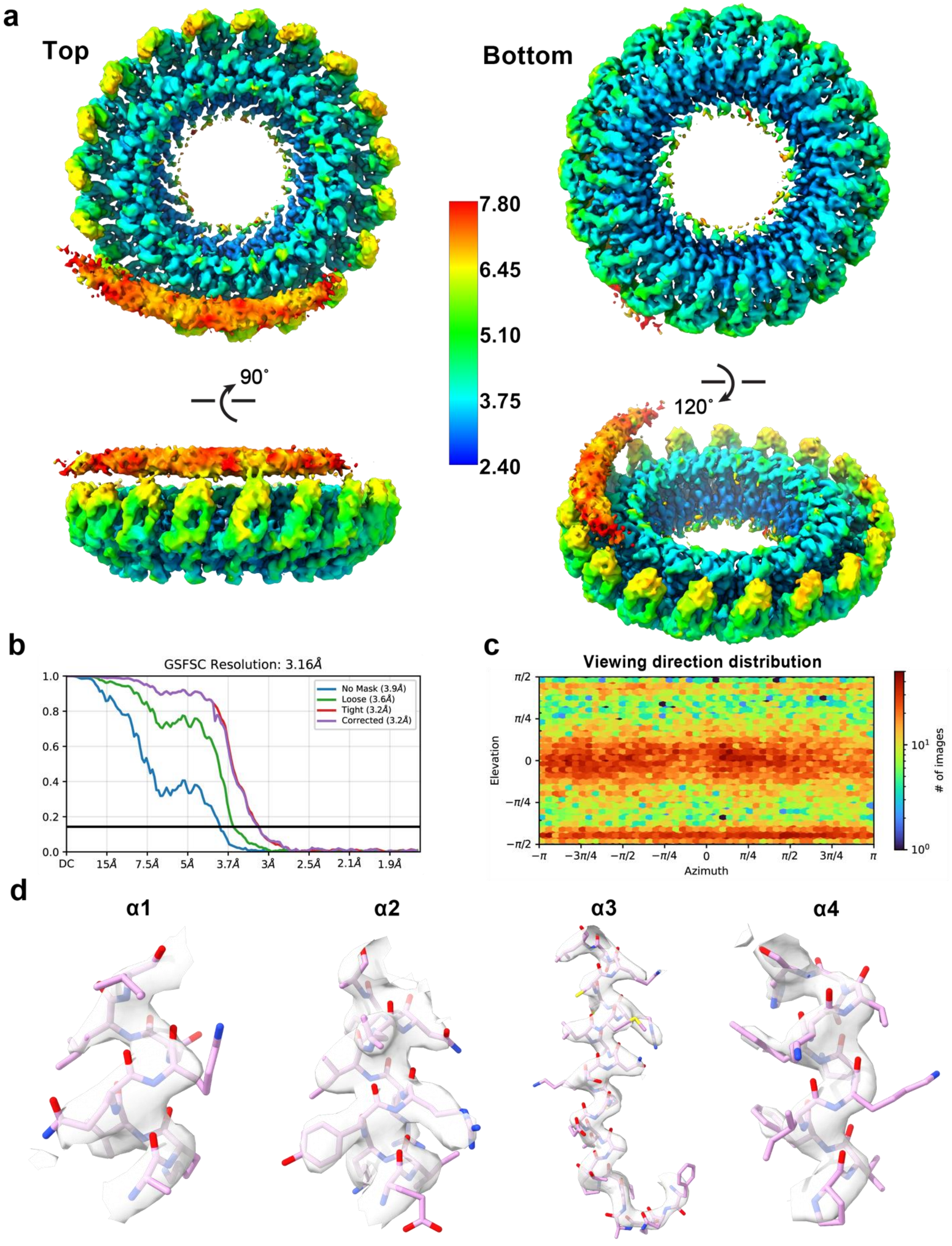

**Figure S22. Cryo-EM structure of Mgm101 with B-form DNA. (a)** The reconstruction (PDB ID: 9YI8, EMD-72981, EMPIAR-13025) colored by local resolution estimation. (**b**) GSFSC resolution. (**c**) Particle viewing direction distribution (**d**) Mgm101 $\alpha$-helices and their fit within the density.

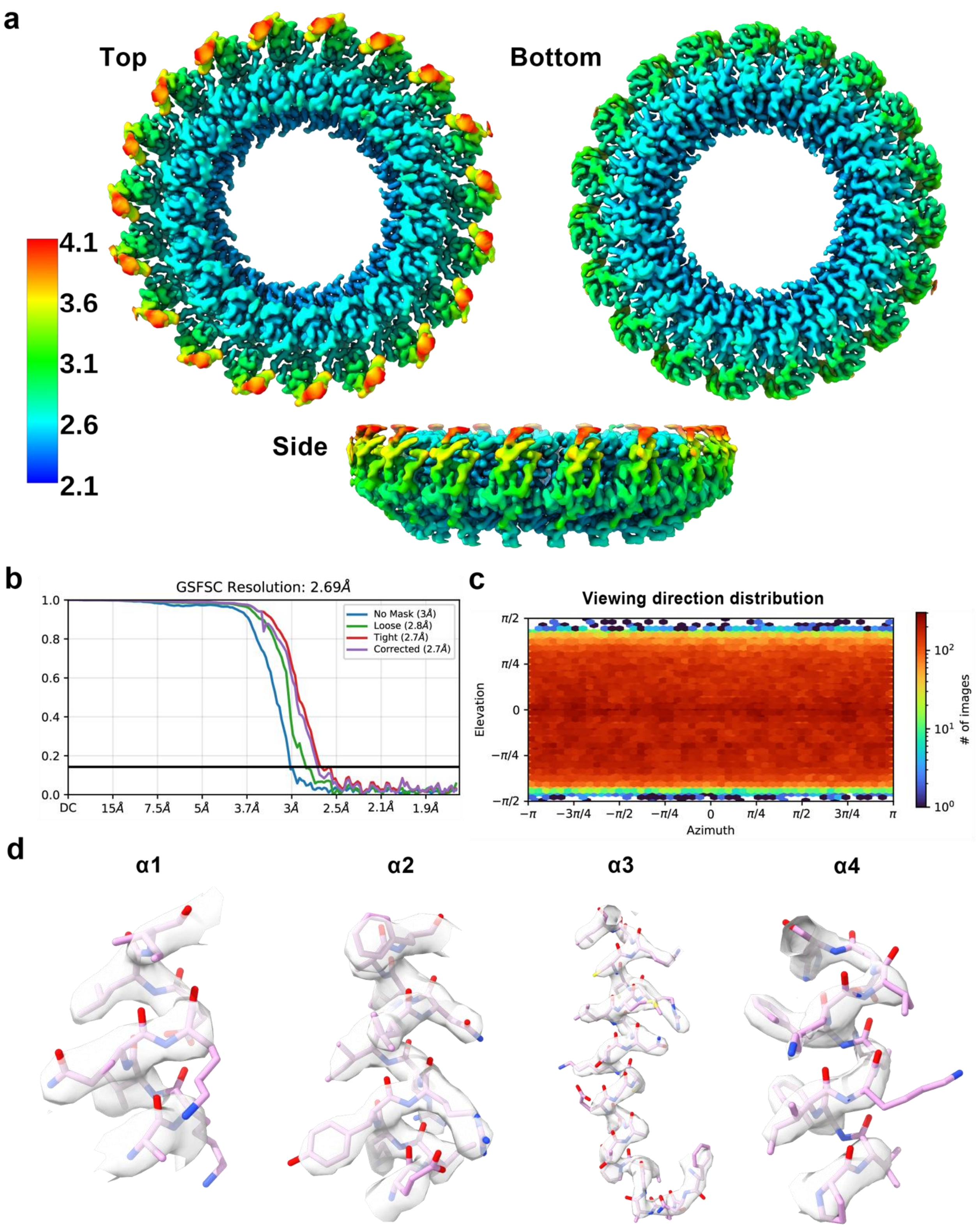


**Figure S23. Cryo-EM structure of the apo Mgm101 ring** (**a**) The reconstruction (PDB ID: 9YIA, EMD-72984, EMPIAR-13024) colored by local resolution estimation. (**b**) GSFSC resolution. (**c**) Particle viewing direction distribution. (**d**) Mgm101 α-helices and their fit within the density.

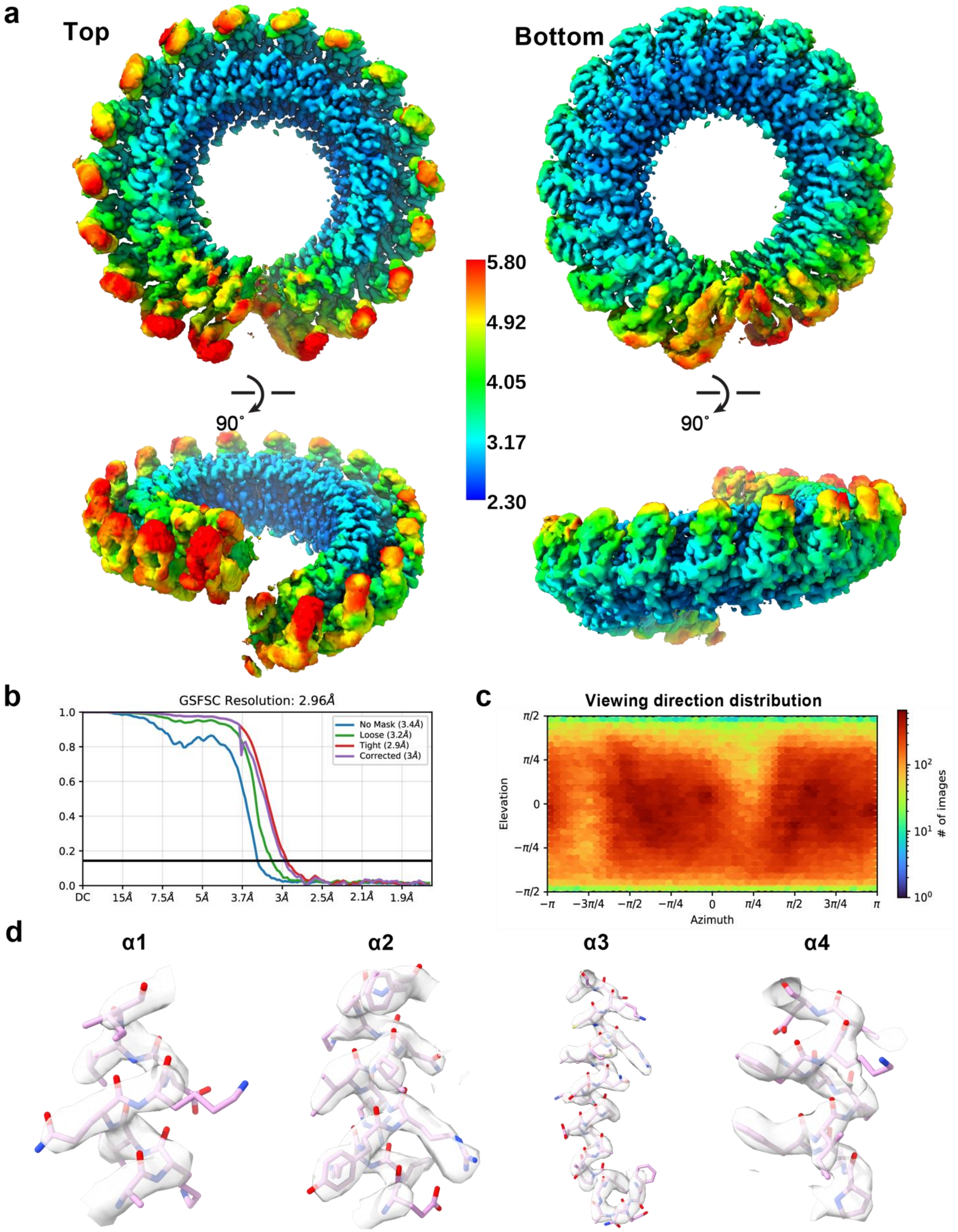

**Figure S24. Cryo-EM structure of the Mgm101 open-ring lock-washer.** (**a**) The lock-washer reconstruction (PDB ID: 9YI9, EMD-72983, EMPIAR-13024) colored by local resolution estimation. (**b**) GSFSC resolution. (**c**) Lock-washer particle viewing direction distribution. (**d**) Mgm101 $\alpha$-helices and their fit within the density.

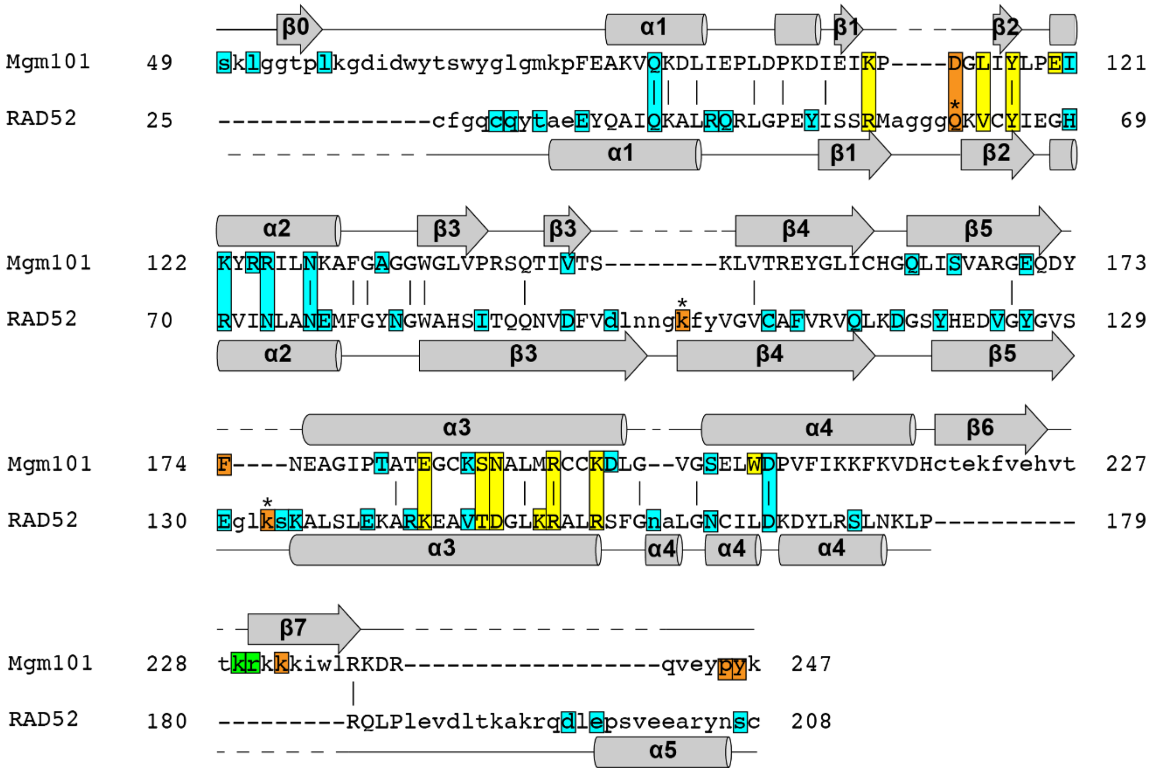


**Figure S25. Mgm101-RAD52 structure-based sequence alignment.** Residues are colored according to their contacts with the inner ssDNA strand (yellow), the outer ssDNA strand (orange), B-form DNA (green), or inter-subunit interactions (cyan), where * represents possible interactions of RAD52 with an outer ssDNA strand. The alignment was performed with the DALI server[8].

**Movie S1. Continuous modeling of the lock-washer from 3D variability component 1.** Variability in the diameter of the lock-washer configuration.

**Movie S2. Continuous modeling of the lock-washer from 3D variability component 2.** Variability along one end of the lock-washer termini, showing the appearance of density for additional subunits.

**Movie S3. Continuous modeling of the lock-washer from 3D variability component 3.** Variability along the other end of the lock-washer termini, again showing the appearance of density for additional subunits.

**Movie S4. Continuous modeling of the lock-washer from 3D variability component 4.** Variability in the planarity of the lock-washer termini, indicating flexibility.
